# Adaptation to seasonal drought in *Arabis alpina* is linked to the demographic history and climatic changes since the last glacial maximum

**DOI:** 10.1101/2025.09.09.671043

**Authors:** Bastiaan Tjeng, Mehak Mehak, Helene Bråten Grindeland, Jasmin Zohren, Andrea Dalla Libera, Jörg Wunder, Carlos Alonso-Blanco, George Coupland, Andrea Fulgione

## Abstract

Understanding how species adapt to new environments is a central goal in evolutionary biology, and a topical question in climate change research. Here, we sequenced the genomes of 426 individuals of the perennial, Arctic-alpine herb *Arabis alpina* to study demography and adaptation, with a focus on populations in Northern Spain, that experience warm and dry summers. Our inference supports a scenario in which *A. alpina* colonized Northern Spain in a range expansion event that started near the Alps around 216 thousand years ago (kya). During the last glacial episode (115 to 12 kya), this expansion proceeded Westward, and effective population sizes were large across Europe, likely due to a larger suitable habitat for *A. alpina*. These ancient demographic events gave rise to a highly diverged genetic lineage in Northern Spain. In the present interglacial (between 12 kya and present), populations became increasingly fragmented, and lost genetic diversity across Europe. Furthermore, we detected signatures of selection at genes associated with responses to abiotic stress, including drought stress, and regulation of growth, for instance at *SC5D* and *NAC055*, which reflects the climatic changes since the last glacial period. Notably, an ancient polymorphism at the gene *FRL1* emerged as a candidate for conferring variation in flowering behavior, and for contributing to adaptation to drought. Our study suggests that the combination of ancestral variation in flowering behavior, and positive selection on new mutations involved in drought responses, underlies the evolution of a new trait syndrome, and adaptation to climate change.

## Introduction

Understanding how species shifted and expanded their range through time, and whether and how they can adapt to new environments are central goals of evolutionary biology, and topical questions in climate change research (Waldvogel et al. 2020; Aguirre-Liguori et al. 2021). In the current Quaternary period (2.6 million years ago to present), species experienced repeated glacial-interglacial cycles that influenced their distribution range and pattern of genomic variation (Hewitt 2000; Petit et al. 2003). As species move around the landscape, they can encounter new environmental conditions, they can adapt, and colonize new habitats (Davis and Shaw 2001; Luqman et al. 2023). Depending on the genetic architecture of the traits under selection, population genetic models describe how adaptation can happen through a rapid increase in population frequency of a beneficial allele (a selective sweep) from single or recurrent *de novo* mutations, or from standing genetic variation, or through infinitesimal allele frequency shifts at many loci (Stephan 2016). However, a particularly challenging scenario is when multiple environmental conditions change simultaneously, which can result in selective pressures acting on several traits. For instance, current climate change is characterized by simultaneous changes in average temperature, seasonality, precipitation, and in climatic extremes such as heatwaves, storms, and droughts (Calvin et al. 2023). In this scenario, adaptation may require the evolution of new combinations of trait values (new trait syndromes), which may be considerably more complex than a selective sweep. If adaptation proceeds from new genetic variants that influence variation at multiple traits, the waiting time may be long. Alternatively, adaptation might proceed from standing genetic variation. In this case, the covariance structure among traits, shaped by the past history of a population, may align or misalign with multivariate selection, which can respectively increase or constrain the potential for adaptation (Lovell et al. 2013; Etterson and Shaw 2001; Schluter 1996; Kirkpatrick 2009). With multivariate selective pressures, it is unclear which dynamics would lead to adaptation in natural populations.

An interesting case of adaptive trait covariance, is when species segregate for alternative seasonal strategies (Moran et al. 2016). For instance, in annual plants like *Arabidopsis thaliana*, traits affecting life history, such as the timing of germination (partly regulated by seed dormancy) and of flowering onset, growth rates, and stress responses underlie local adaptation and often co-vary (Donohue 2002; Alonso-Blanco et al. 2009; Picó 2012; Chiang et al. 2013; Krämer 2015; Vasseur et al. 2018; Takou et al. 2019). Correlations among these traits create alternative, seasonal life-history strategies, such as the winter annual and the spring (or summer) annual habits, which segregate within species (Laibach 1951; Rédei 1970). Winter annual plants usually germinate in autumn, overwinter as vegetative rosettes, and flower and set seeds the next spring. In winter annuals, early flowering is inhibited through the vernalization pathway, mainly by the action of *FLOWERING LOCUS C (FLC)*, which is upregulated by *FRIGIDA (FRI)*, until flowering is induced by a prolonged period of cold temperatures (vernalization) (Andrés and Coupland 2012). Conversely, spring annuals germinate in spring, and flower and set seeds in the same growing season, which results in a faster life cycle. The spring annual habit is associated with multiple nonsense mutations at *FRI*, and in rare cases at *FLC*, which impair the vernalization pathway and accelerate flowering (Johanson et al. 2000; Méndez-Vigo et al. 2011; Fulgione et al. 2022; Stinchcombe et al. 2004; Zhang and Jiménez-Gómez 2020). This habit tends to occur in disturbed, ruderal landscapes, and in Mediterranean climates, where the ability to set seeds before the onset of drought or disturbance enables an escape strategy (Ludlow 1989; Mckay et al. 2003; Franks et al. 2007; Kooyers 2015). These alternative seasonal strategies are characterized by the co-variation of flowering time, germination time, and possibly various other traits including growth rates (Debieu et al. 2013), defense response (Glander et al. 2018; Davila Olivas et al. 2017), and stress responses (Ludlow 1989; Mckay et al. 2003; Franks et al. 2007; Des Marais et al. 2012; Lovell et al. 2013; Easlon et al. 2014; Kooyers 2015; Davila Olivas et al. 2017). Overall, life-history traits can evolve jointly in annual plants, and whether current trait correlations aid or constrain adaptation to future multivariate selection pressures, is still an open question.

The adaptive value of seasonal strategies differs in perennial plants, which account for about 90% of all plants (The Angiosperm Phylogeny Group 2016). In perennials, seasonal strategies can derive from life-history trade-offs, for instance among vegetative growth, survival, and reproductive effort in the current or future growing seasons (Stearns 1989; Reich 2014). A growth-survival trade-off is for instance well established in perennial grasses with a seasonal endo-dormant phase (Gillespie and Volaire 2017; Volaire 2018). In the grass *Dactilis glomerata*, individual plants that are winter- and summer-dormant have higher survival with winter frost and summer drought, respectively (Bristiel et al. 2018). In the ryegrass *Lolium perenne*, some individuals have high growth rates and are sensitive to drought and frost stress, others are drought-tolerant and have low summer growth rates, and others are frost-tolerant and have low winter growth rates (Keep et al. 2021). Growth-survival trade offs have been further described along altitudinal (Benavides et al. 2015), and latitudinal clines (Koehler et al. 2012) in trees. A trade-off between vegetative growth and reproductive effort has been described for instance in the genus *Fragaria*, in which a deletion in the *GA20ox* gene regulates the fate of axillary meristems between vegetative growth through the production of runners, and flowering (Tenreira et al. 2017). In *Arabis alpina*, an emerging model for perennial Brassicaceae (Wötzel et al. 2022), the *FLC* ortholog *PERPETUAL FLOWERING 1 (PEP1)* represses flowering in the absence of vernalization and facilitates a return to vegetative growth after flowering (Wang et al. 2009). Furthermore, the gene *TFL1* regulates the juvenility phase, the development of the inflorescence, and whether the axillary shoots commit to flowering or remain vegetative (Wang et al. 2011). This gene has similar functions in some tree species (Jensen et al. 2001; Kotoda et al. 2006; Mohamed et al. 2010). In many perennials, the regulation of flowering varies within species (Friedman 2020), however its adaptive value differs from annuals, in which early flowering followed by senescence can directly translate into an escape strategy. For instance, in *A. alpina*, a screen of flowering behavior revealed that plants from most European regions varied in the time of flowering onset without vernalization, and in the duration of flowering after vernalization (Wunder et al. 2023). The most notable exception were populations from the Cantabrian Mountains in Northern Spain, in which all individuals were unable to flower without vernalization, and flowered for a short time after vernalization. This flowering behavior was hypothesized to contribute to adaptation to summer drought (Wunder et al. 2023).

Here, we study the perennial, Arctic-alpine herb *A. alpina*, and in particular the populations from Northern Spain, which are locally adapted on the basis of common garden experiments (Toräng et al. 2015). Furthermore, early flowering was associated with lower winter survival in these populations (Wunder et al. 2023), which suggests a trade-off between reproductive effort and survival. *A. alpina* is distributed across the alpine and amphi-Atlantic regions of Europe, the Middle East and Africa, and is thought to have colonized Europe 0.3 to 1.4 million years ago, and Scandinavia in the current interglacial (Ehrich et al. 2007; Koch et al. 2006; Ansell et al. 2011; Karl et al. 2012; Laenen et al. 2018). However, the history of colonization of Spain, and of adaptation to the local conditions are not yet resolved. In this study, we aim to understand the adaptive value of seasonal strategies and of flowering behavior in a perennial plant. To this end, we collected and sequenced the genomes of 211 individual plants (accessions) from Northern Spain, and of 215 from related populations in the French Alps for comparison. We characterized genomic diversity and population structure, we reconstructed the demographic history of colonization of Northern Spain, and we identified genomic signatures of adaptation.

## Results

### Plant material and SNP calling

We sequenced the whole genomes of 211 individual *A. alpina* plants collected in 14 populations in the Cantabrian Mountains (Northern Spain), and of 215 individual plants collected in 15 populations in the French Alps (Supplementary Table S1, Fig. 1a). A subset of these accessions were sourced from previous studies (Wunder et al. 2023; Tjeng et al. 2024). The average genome-wide sequencing depth was 19.7x (range across individuals: 14.8x - 26.0x). We identified over 7.2 million and 5.8 million high quality SNPs, with less than 30% missing genotypes in the populations from the Cantabrian Mountains and from the Alps, respectively. Using a newly developed approach, ParaMask (Tjeng et al. 2024), we identified a total of 48.6 Mbp and 65.3 Mbp of multicopy genomic regions out of the total 311.6 Mbp of assembled genome for the populations from the Cantabrian Mountains and from the Alps. Additionally, we created a 63.6 Mbp mappability mask using SNPable. After excluding these regions for a total of 125.9 Mbp, we retained over 4.6 million SNPs for the samples from the Cantabrian Mountains and over 3.2 million SNPs for the samples from the Alps. After further filtering steps that are specific for some of the analyses (described in Materials and Methods), we used 916,101 SNPs for the neighbour joining tree and the principal coordinate analysis (PCoA), 629,967 SNPs for ADMIXTURE, and 2,967,704 SNPs for demographic inference. Depending on the test set used in selection scans with *3P-CLR*, we used between 2.3 and 2.5 million SNPs.

**Figure 1.**
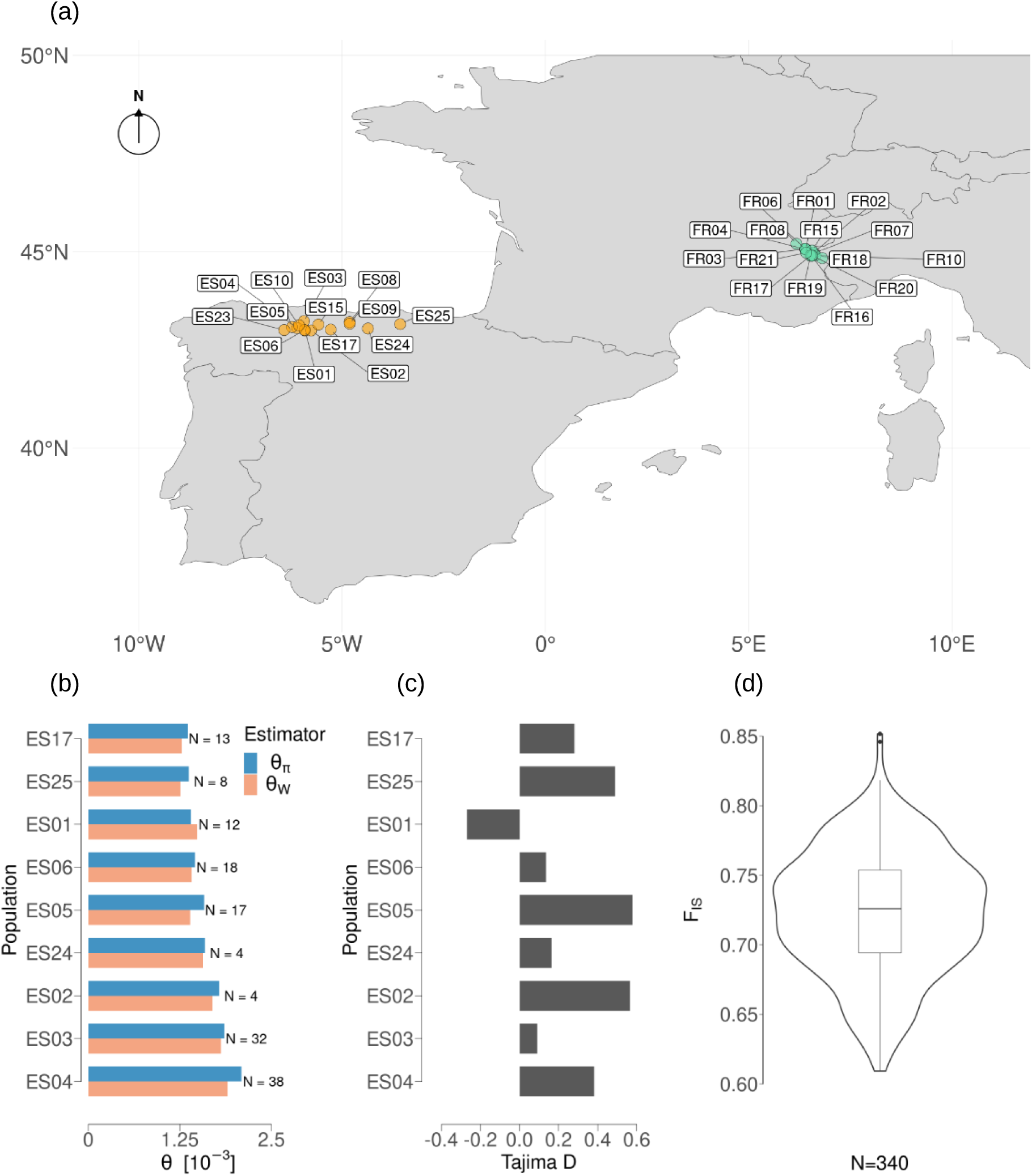
Map of the sampling sites and genomic summary statistics estimated from unrelated samples. **a:** Map of the sampling sites in Northern Spain (yellow) and in the French Alps (green). Each point represents one population. **b:** Estimators of the population mutation rate (*θ_π_* in blue, and *θ_W_* in red) for populations with a sample size *N* ≥ 4. Sample sizes are indicated on the right of each bar. **c:** Tajima’s D estimated for populations with a sample size *N* ≥ 4. **d:** Global inbreeding coefficient (*F_IS_*), estimated for populations with a sample size *N* ≥ 10 and calculated in bins of 10k SNPs.

### Populations in the Cantabrian Mountains (Northern Spain) are genetically differentiated from each other

In order to understand the general pattern of genetic variation in the populations from the Cantabrian Mountains, we characterized genetic diversity and population structure. Genetic diversity measured with Watterson’s estimator of theta (*θ_W_*), and the average pairwise differences (*θ_π_*), ranged from 1.26 × 10*^−^*^3^ and 1.36 × 10*^−^*^3^ in population ES25 to 1.9 × 10*^−^*^3^ and 2.1 × 10*^−^*^3^ in population ES04, respectively (Fig. 1b). Tajima’s D was positive in almost all populations, and ranged between 0.091 in ES03 and 0.48 in ES05, which indicates a slight excess of intermediate-frequency variants (Fig. 1c). Population ES01 was the only exception with a negative Tajima’s D of -0.268, suggesting a recent population expansion. The mean inbreeding coefficient (*F_IS_*) across bins of 10,000 SNPs was estimated at 0.722, with a standard deviation of 0.047 (Fig. 1d). Considering that these populations are self-compatible (Toräng et al. 2017; Laenen et al. 2018), this corresponds to an outcrossing rate of 16.1% if inbreeding is only due to selfing, or higher than that if other causes of inbreeding also play a role, for instance population structure.

The average density of genetic differences per pair of samples was 1.59-fold higher across populations than within populations (on average 3.11×10*^−^*^3^ across populations and 1.96×10*^−^*^3^ within populations; Supplementary Fig. S1a). The distribution within populations was unimodal, and had a tail of low-density pairwise differences due to closer relatedness among individuals. The distribution among populations, on the other hand, was multimodal with three main peaks of pairwise differences. The mode at 2.81 × 10*^−^*^3^ differences included pairs of individuals from Western locations. The mode at 3.50 × 10*^−^*^3^ differences included pairs with one individual from a Western population, and one from a Central-Eastern location, excluding the Eastern-most population, ES25. The mode at 4.00 × 10*^−^*^3^ differences included all sample pairs with one individual from ES25. Consistent with a high selfing rate, most individuals had low heterozygosity (median = 1.68 × 10*^−^*^4^), and few had heterozygosity in the range of the pairwise differences within populations (up to 2.2 × 10*^−^*^3^), likely due to rare outcrossing events (Supplementary Fig. S1b). The distribution of pairwise differences was strongly correlated with geographical distance, consistent with isolation by distance (IBD) (Mantel correlation = 0.799, p-value *<* 1 × 10*^−^*^4^; Supplementary Fig. S1c).

Clustering of the genomic data with a principal coordinate analysis (PCoA) based on a distance matrix identified three main genetic clusters (Fig. 2a). The first coordinate of the PCoA separated Central-Eastern populations (CE-CAN), including samples from locality Puebla de Lillo (ES17), Picos de Europa (ES08 and ES09), and Eastern locations (ES24 and ES25). The second coordinate differentiated two groups within the Western region (W-CAN1 and W-CAN2), one including localities between Angliru and Valle de Riotuerto (ES01, ES02, ES03, ES06, ES10, ES15), and the other including localities between Leitariegos and La Cueva (ES04, ES05, ES23). Similarly, a neighbor-joining tree separated the Central-Eastern populations with a long internal branch, and the two groups of Western populations with shorter internal branches (Fig. 2b). In both analyses, genetic clustering aligns with the geographic location of sample collection, suggesting clear differentiation among populations. We analyzed hierarchical population structure with ADMIXTURE, after sub-sampling the data set to 173 unrelated individuals (Materials and Methods). This analysis identified six ancestry groups based on cross-validation (CV) errors across 10 replicate runs (mean CV: 0.2534), and similarly low CV for up to nine groups (t-test: p-value = 0.2916, Supplementary Fig. S2). Across runs, populations from CE-CAN were assigned to the same ancestry group, despite a relatively large geographical distance between populations (at most around 65 km; Fig. 2c). Genetic ancestry was private to most other populations, with the exception of populations ES23 - ES04, and ES01 - ES06, respectively within W-CAN1 and W-CAN2, which shared genetic ancestry (Fig. 2c). Overall, the analysis of genetic diversity and population structure revealed clear differentiation among populations in the Cantabrian Mountains, and in particular among Central-Eastern locations and two genetic clusters in the Western area.

**Figure 2.**
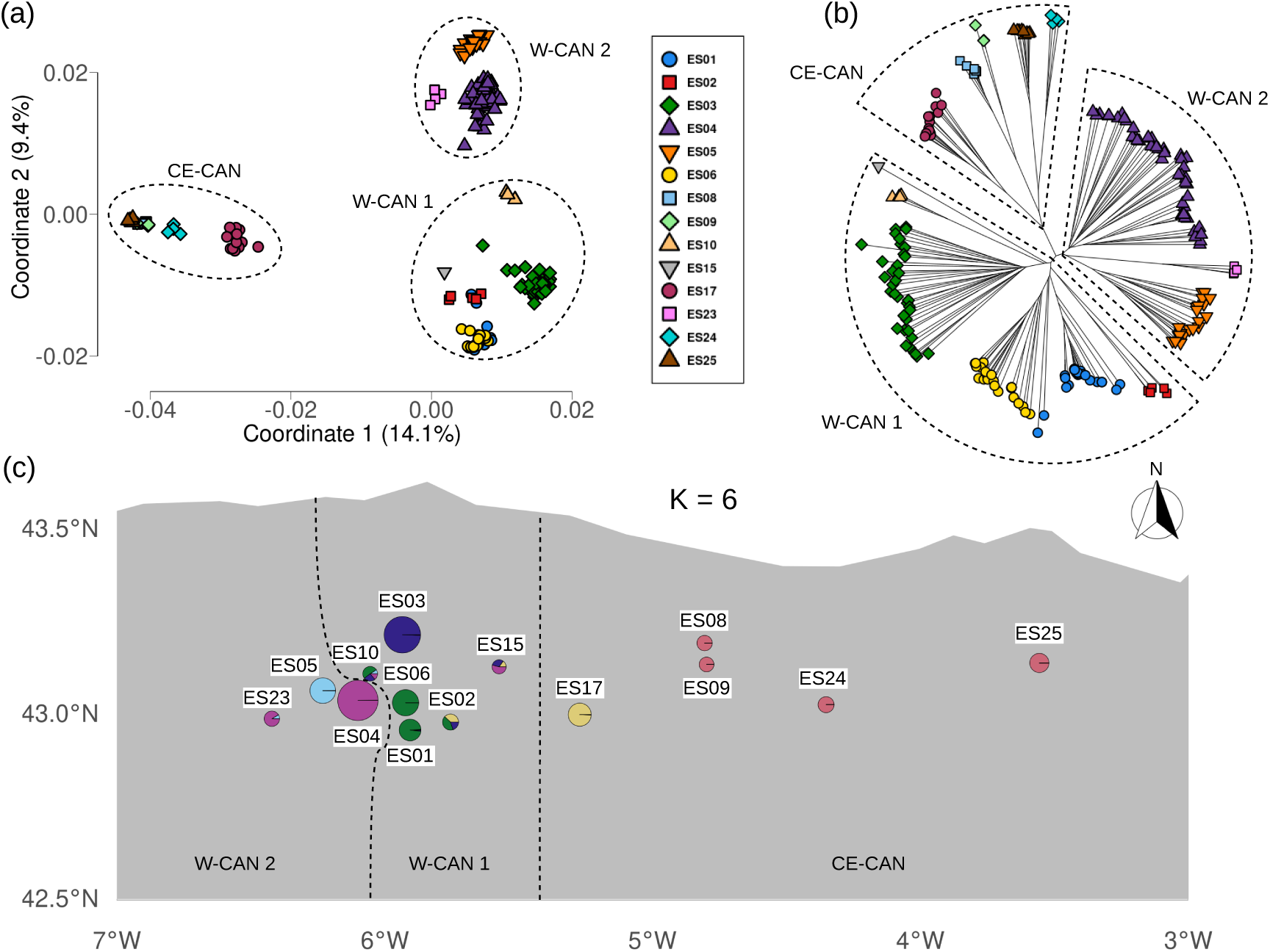
Analysis of genetic clustering among *A. alpina* populations from the Cantabrian Mountains. The three major structure groups Central-Eastern Cantabrian Mountains (CE-CAN), Western Cantabrian Mountains 1 (W-CAN1), and Western Cantabrian Mountains 2 (W-CAN2) are highlighted by dashed lines. **a:** Principal coordinate analysis (PCoA) based on genetic distances (1-IBS, identity by state). Samples are arranged in the first two coordinates, which capture the greatest variation of the original distances. Labels indicate the percentage of variation explained by each axis. Shapes and colors denote different populations. **b:** Unrooted neighbor-joining (NJ) tree. Shapes and colors denote different populations, as in panel (a). **c:** Geographic distribution of shared genetic ancestry inferred with ADMIXTURE. The number of ancestry groups is set to *K* = 6, based on cross-validation error. The pie charts represent the proportions of genetic ancestry, and circle diameters are proportional to sample size.

### Demographic inference reveals that effective population sizes were large during the last glacial period, and that the colonization of the Cantabrian Mountains proceeded from east to west

We reconstructed the demographic history of populations from the Cantabrian Mountains and from the French Alps using the hidden Markov model (HMM) based method Relate (Speidel et al. 2019). Consistent across populations, this inference revealed a period of low effective population size between around 400 and 200 thousand years ago (kya) (Fig. 3a). This was followed by a period of large effective population size that started between 100 and 200 kya, and lasted until between 10 and 20 kya. This period approximately coincides with the last glacial period, between 115 and 11.7 kya, but it may also have been influenced by the penultimate glacial period, between 194 and 135 kya (Dahl-Jensen et al. 2013). Subsequently, effective population sizes declined until present, in a period that roughly coincides with the current interglacial, between the last glacial maximum (LGM), around 21 kya, and present (Clark et al. 2009).

**Figure 3.**
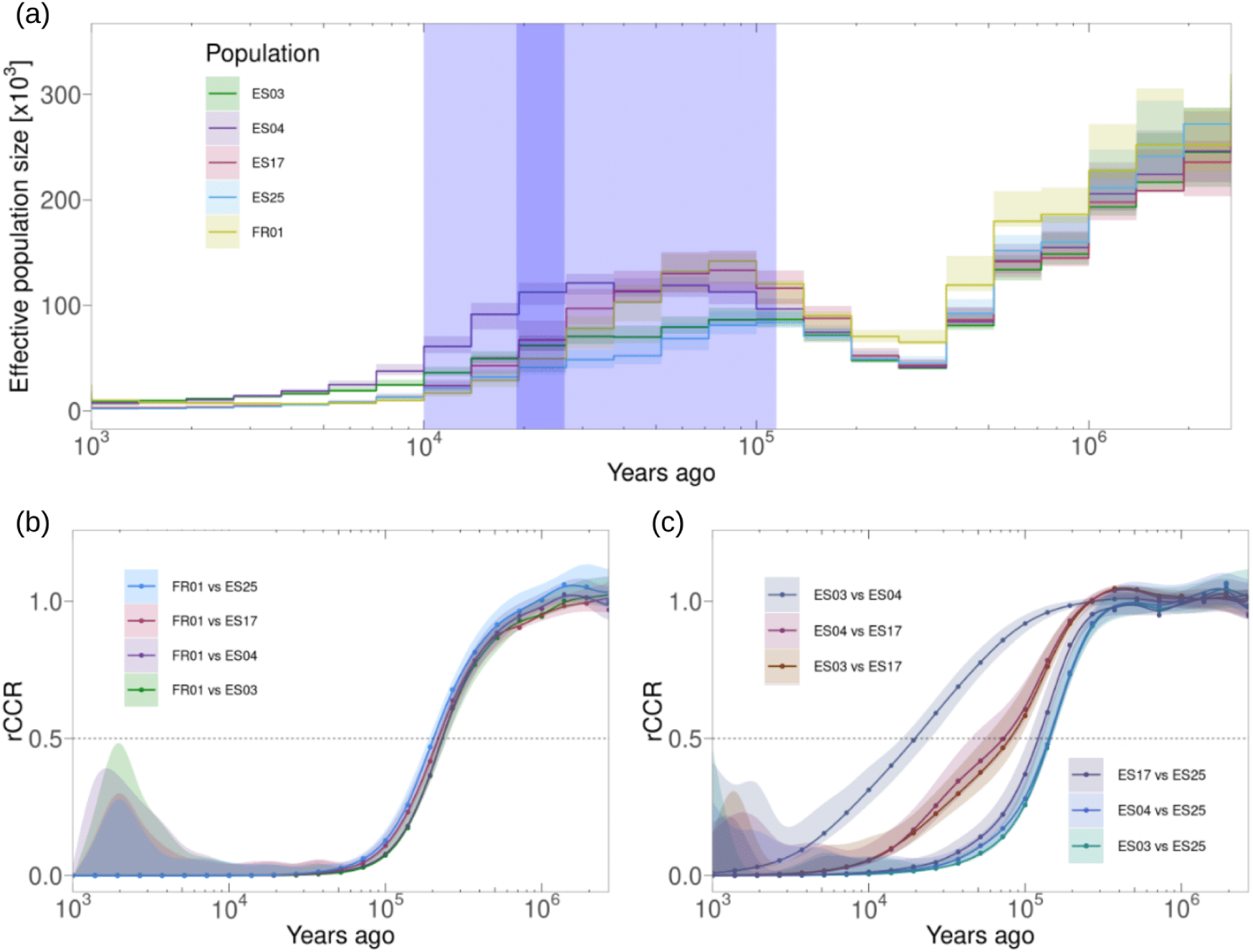
Estimates of effective population size as a function of time, and of split times among populations based on the relative cross coalescent rates (rCCR). Lines represent point estimates, and shaded colors represent the 95% confidence intervals based on 100 bootstrap replicates. Four representative populations are shown (ES03, ES04, ES17, and ES25), and the French population FR01 is used as an outgroup. Split times for all population pairs can be found in Supplementary Table S2. **a:** Effective population size across time. The plot area shaded in light blue represents the last glacial episode, and the area shaded in dark blue represents the last glacial maximum. **b:** rCCR between populations from the Alps and from the Cantabrian Mountains. A cutoff of *rCCR* = 0.5, indicated by the horizontal line, is used to infer split times. **c:** rCCR among populations from the Cantabrian Mountains. A cutoff of *rCCR* = 0.5, indicated by the horizontal line, is used to infer split times.

To infer the time in the past when populations split from each other, we used the relative cross coalescent rate (rCCR) statistic, and a threshold of 0.5 (Schiffels and Durbin 2014). Split times between populations in the Cantabrian Mountains and FR01, which we used as an outgroup from the French Alps, ranged between 198 and 230 kya (Fig. 3b), dating back to the start of the penultimate glacial period, around 194 kya. Split times among populations from the Cantabrian Mountains ranged between 6 and 150 kya (Fig. 3c, Supplementary Table S2), mostly during, or just predating the last glacial period. In comparisons between populations from CE-CAN and W-CAN1 or W-CAN2, split times were 1.1 to 8.4 (mean: 3.8) times older than the splits among W-CAN groups. Split times within ancestry groups ranged between 6 and 34 kya in W-CAN1, between 10 and 20 kya in W-CAN2, and between 23 and 140 kya in CE-CAN. These recent split times within ancestry groups suggest that fragmentation among populations mostly happened in the current interglacial, since the LGM. Split times were positively correlated with the longitude of the eastern-most population across population pairs (pearson correlation: *r* = 0.836; p-value = 2.2 × 10*^−^*^16^, Supplementary Fig. S3), implying that the Cantabrian Mountains were colonized from east to west.

### The Cantabrian Mountains are characterized by warmer temperatures year-around, and lower precipitation in summer, compared to other collection sites in Europe

Because our reconstruction of the history of *A. alpina* in the Cantabrian Mountains is consistent with an important role of glacial and interglacial periods, we characterized the climatic conditions at collection sites across Europe. We used temperature and precipitation records for the years 1970-2000 from https://www.worldclim.org/ at the collection sites described in Wunder et al. (2023), which were mostly in the Cantabrian Mountains, in the Alps, and in Scandinavia. In addition, we used the Aridity Index (AI) from the Global Aridity Index database *v*3 (Zomer et al. 2022), which is based on precipitation and potential evapotranspiration. The average monthly temperatures at the sites in the Cantabrian Mountains were above freezing year-around (Supplementary Fig. S4a). In contrast, average monthly temperature in the Alps and in Scandinavia was 5.4°C and 8.8°C lower than in the Cantabrian Mountains, and it was below freezing from November to April and from October to April, respectively. Annual precipitation was highest in the Alps (mean: 1322.0 mm), followed by the Cantabrian Mountains (mean: 1151.1 mm), and lowest in Scandinavia (mean: 890.8 mm). Moreover, the seasonal course of precipitation largely differed among regions (Supplementary Fig. S4b). In Scandinavia, precipitation peaked in summer (July to October), while in the Alps and in the Cantabrian Mountains, precipitation was at a minimum in summer (June to September). As a result of warmer temperatures year-around, and lower precipitation in summer, the sites in Northern Spain are more arid in summer, especially between mid-June and September (Supplementary Fig. S4c).

### Detection of genomic regions with signatures of positive selection

Our demographic inference revealed a decrease in genomic diversity concomitant with a change in climatic conditions in the current interglacial. To understand whether and how *A. alpina* adapted to the changing conditions, we screened the genomic data for signatures of positive selection using *3P-CLR* (Racimo 2016). This method can account for structured populations, and detect selective sweeps that happened in the two terminal branches that lead to each of two related focus populations, and sweeps that happened in the branch that is ancestral to both, compared to an outgroup. For each genetic cluster, we first obtained estimates of drift rates, which are rates of allele frequency changes due to genetic drift that depend on divergence times and on effective population size, *N_e_*, and are equivalent to F3 statistics (Racimo 2016). Estimated drift rates were highest for divergence between Northern Spain and France (2.78 to 3.35 across comparisons, which corresponds to a drift time of 5.56 to 6.70 *N_e_*generations), followed by divergence between CE-CAN (population ES17) and W-CAN1 and W-CAN2 (0.22 to 0.65), and between W-CAN1 and W-CAN2 (0.08 to 0.18; Supplementary Table S3). Drift rates between W-CAN and pooled CE-CAN were negative, which suggests that CE-CAN (excluding population ES17) is partly admixed with populations in the French Alps. As a consequence of this, we used as focus clusters W-CAN1, W-CAN2, and population ES17 as a representative of CE- CAN, and we used as outgroups the populations from the French Alps, and from CE-CAN, excluding ES17 (the sets of populations used are shown in Supplementary Table S3). Estimated drift rates were consistent with the results from demographic inference (more detail in Supplementary Materials).

The top five CLR peaks on the terminal branches that lead to W-CAN1, W-CAN2, and ES17 (CE-CAN) overlapped genes with functions in lipid biosynthesis and membrane fluidity (in the endoplasmic reticulum and chloroplast), cell-wall modification and scaffolding, and the synthesis and turnover of surface-secreted metabolites (Supplementary Table S4). Genes within the upper 0.5% tail of CLR scores were enriched in GO categories associated with energy generation, solute transport, core metabolism, and abiotic stress responses (p-value ≤ 0.5%; Supplementary Table S5). On the branch ancestral to W-CAN1, W-CAN2, and ES17, the top five CLR peaks overlapped genes involved in growth control, protein turnover, membrane assembly and metabolism, and defense mechanisms (Supplementary Table S4). The top 0.5% CLR peaks on the ancestral branch were enriched for genes related to the negative regulation of growth and positive regulation of transcription (Supplementary Table S5). Because divergence to the French Alps was ancient, we repeated the analysis for W-CAN1, W-CAN2, and ES17, using CE-CAN as outgroup (excluding ES17). In the resulting terminal branches, the top five CLR peaks overlapped genes with roles in sterol metabolism, RNA processing, abiotic stress responses, hormone- and light-mediated growth responses, and translational capacity (Supplementary Table S4). The upper 0.5% CLR peaks overlapped genes enriched in membrane trafficking and secretion, lipid and pigment metabolism, proteostasis, calcium signaling, and RNA splicing (Supplementary Table S5). On the branch ancestral to W-CAN1, W-CAN2 and ES17, the top five CLR peaks overlapped genes linked to hormone transport and metabolism, ion homeostasis, defense, abiotic stress responses, and meiotic recombination (Supplementary Table S4). The top 0.5% CLR peaks were significantly enriched in genes related to responses to environmental cues, including nutrient and oxidative stress, as well as developmental processes (Supplementary Table S5). Across all combinations of focus populations and outgroups, the top CLR peak was on the terminal branch leading to W-CAN2, and overlapped the gene *SC5D* (Fig. 4a). *SC5D* encodes a C-5 sterol desaturase, and a premature stop codon in *A. thaliana* impairs brassinosteroid biosynthesis, and underlies the dwarf mutant *dwf7* (Choe et al. 1999).

**Figure 4.**
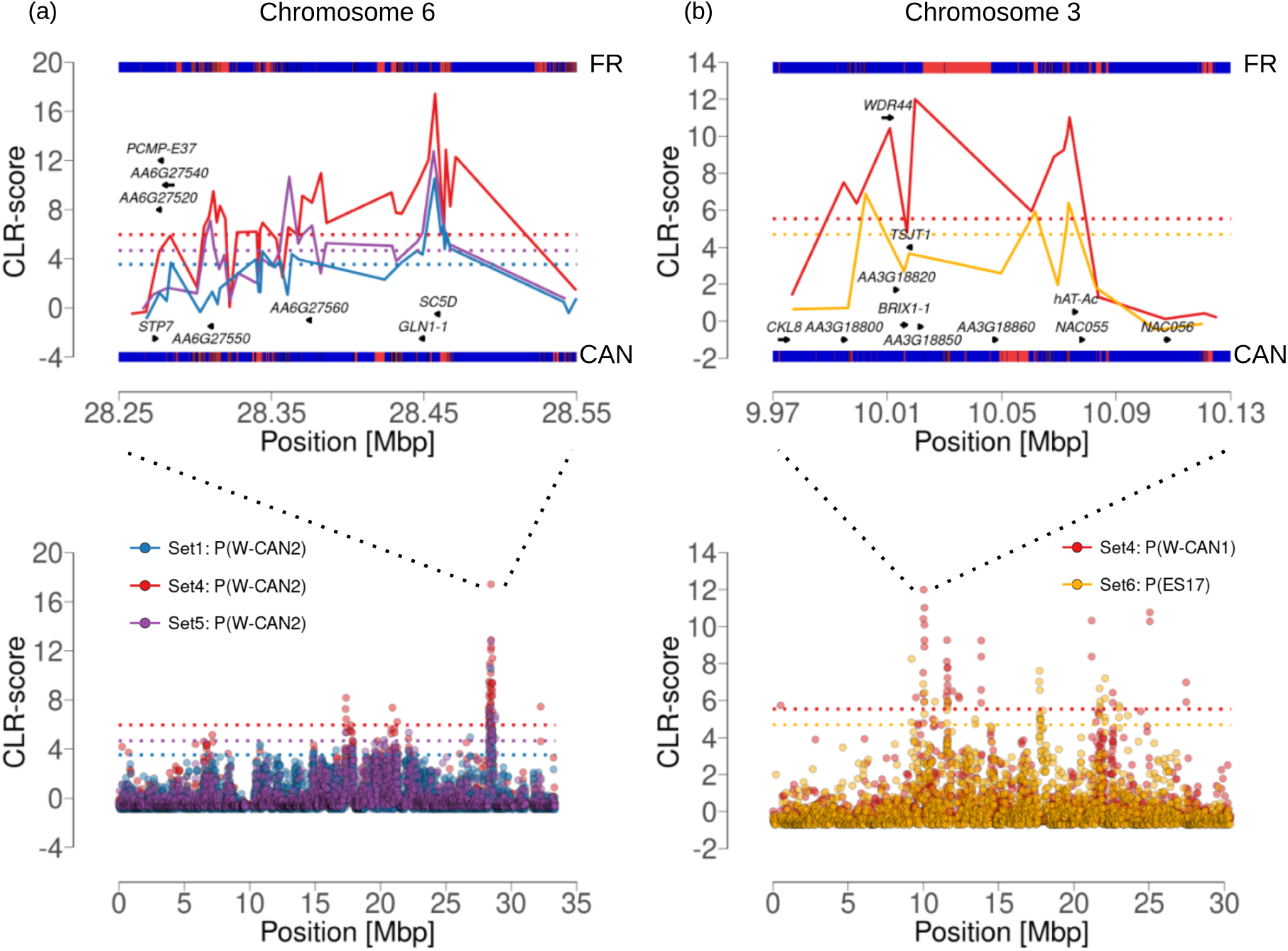
Standardized composite likelihood ratio scores at the *SC5D* (**a**) and *NAC055* (**b**) loci, shown zoomed in (top) and across the chromosome (bottom). Genes are indicated by arrows and labeled with their *A. thaliana* ortholog or, if no ortholog is known, by their *A. alpina* gene identifier. Dotted lines show the 0.5% most extreme CLR scores for each analysis. Segments in the top panels show multicopy (red) and single-copy (blue) regions in France (top segment) and in the Cantabrian Mountains (bottom segment) inferred by ParaMask. The sets in the legend represent different combinations of populations (explained in Supplementary Table S3). **a:** Scores for the private branch of W-CAN2 and for different group sets in the *3P-CLR* analysis (blue – Set1; red – Set4; purple – Set5). **b:** Scores for the private branch of W-CAN1 (red – Set4) and for ES17 (yellow – Set6). The DNA transposon (hAT-Ac) upstream of the gene is also indicated.

To find genes potentially involved in adaptation to summer drought, we analyzed candidate genes with signatures of positive selection, particularly in the terminal branch leading to the most arid population, ES17 (*AI_July_* = 0.250; *AI_August_* = 0.236). We found 16 genes associated with drought-related GO terms (response to water deprivation and heat; GO:0009414, GO:0009408), of which 11 were found on terminal branches, and five on the branch ancestral to W-CAN1, W-CAN2, and ES17 (Supplementary Table S6). Among these, we found transcription factors that are partially responsive to ABA (e.g.: *NAC055*, *ZHD11*, *MYB94*), stress induced enzymes (*ASPG1*, *PAL2*), proteins that stabilize RNA (*RH25*, *RH26*), and chaperones (*HSP15.7*, *Cpn10*, *FKBP62*). At the NAC transcription factor *NAC055* (Fig. 4b) we identified an inframe 3bp insertion at residue 251, which translates into the insertion of a phenylalanine (details in Supplementary Materials). Furthermore, we found a transposable element (TE) insertion 892 bp upstream of the transcription start site in all and only the individuals that lacked the 3bp insertion. The 3bp insertion was fixed in populations ES17 and ES05, which have the two most arid climates in July, and its allelic dosage was significantly correlated with the aridity index in July (Supplementary Fig. S5; linear regression with four principal coordinates from genome-wide structure as covariates; Wald p-value = 4.48 ∗ 10*^−^*^9^; more detail in Supplementary Materials). The 3bp insertion and the TE are candidate genetic variants that might change the function of the gene, and that might underlie adaptation to drought.

In order to identify adaptive genetic variants potentially involved in flowering regulation and vernalization, we inspected the 0.5% CLR tails of all sets and branches. We identified three candidate genes associated with vernalization responses, *VERNALIZATION INDEPENDENCE 4* (*VIP4*) in the terminal branch leading to W-CAN2, *VIN3-LIKE PROTEIN 1* (*VIL1*) in the terminal branch leading to ES17, and *FRIGIDA INTERACTING PROTEIN 1* (*FIP1*) in the terminal branch leading to W-CAN2 (Supplementary Table S7; Supplementary Fig. S6). *VIP4* is a positive regulator of *FLC* expression, and mutants flower even earlier than *FLC* null alleles (Zhang and Van Nocker 2002). At *VIP4*, we identified two non-conservative missense mutations (E212V) and (D184H) located in a disordered region of the protein. Notably, these substitutions change an acidic glutamic acid (E) and an aspartic acid (D) residues, which are conserved in *Arabidopsis* (UniProtKB:Q9FNQ0), *Brassica napus* (EnsemblPlants GeneID:Bra035940), and in *Arabis montbretiana* to a hydrophobic and a positively charged residues, respectively. The two mutations were nearly fixed in W-CAN2 (at 96.6% and 97.8% frequency, respectively), they were segregating in W-CAN1 (at 64.2%, and 66.9% frequency, respectively), and were absent in both CE-CAN and French populations. *VIL1* interacts with *VIN3* to epigenetically silence *FLC* expression (Sung et al. 2006). At *VIL1*, we identified a deletion of residues Asp472–Asn473 within a disordered region upstream of the VIN3-interacting domain. This variant segregated at higher frequency in W-CAN2 (54.9%) than in W-CAN1 (7.4%), and in France (5.9%), and was absent in CE-CAN. *FIP1* encodes a known interactor of *FRI*, however its mutant phenotype in *A. thaliana* flowers at the same time as wild type (Sung et al. 2006).

### Molecular evolution statistics identify genes with a different selection regime in the Cantabrian Mountains compared to the Alps

To detect general differences in the selection regime between populations from the Cantabrian Mountains and from the Alps, we used molecular evolution statistics. In particular, we compared the numbers of non-synonymous and synonymous substitutions (*Dn*, *Ds*), and their rate ratio (*d_N_ /d_S_*), and polymorphisms (*Ps*, *Ps*) and their rate ratio (*p_N_ /p_S_*), across all 28745 annotated genes, using Fisher’s exact test and *A. montbretiana* as the outgroup. The null hypothesis was that the mode and strength of selection acting on genes is the same in the Cantabrian Mountains and in the Alps, including genes that evolve nearly neutrally. Among the genes that significantly differed in substitution rates, fourteen genes had at least partially characterized orthologs in *A. thaliana* or in other species (Table 1), including eight genes with a lower *d_N_ /d_S_*in the Cantabrian Mountains, and six with a higher *d_N_ /d_S_* in the Cantabrian Mountains than in the Alps. These genes were associated with abiotic stress responses, defense mechanisms, genome maintenance, and epigenetic regulation of chromatin (Table 1). Genes related to abiotic stress, in particular, had more specific functions in cell wall biosynthesis, protein folding, transcriptional regulation, and epigenetic control. For instance, one of the genes with a significantly different substitution rate was a hexosyltransferase-encoding gene with homology to *MUCI70*, which is involved in seed coat mucilage biosynthesis in *A. thaliana* (Voiniciuc et al. 2018). In plants, seed mucilage contributes to traits such as water retention, dormancy regulation, soil adhesion, and protection against pathogens, which are thought to contribute to adaptation to environmental conditions (Yang et al. 2012). Another gene that differed in substitution rate was *SDC*, an imprinted gene that is epigenetically silenced during vegetative growth through non-CG DNA methylation mediated by the methyltransferases DRM1, DRM2, and CMT3. De-repression of *SDC* can lead to developmental abnormalities (Henderson and Jacobsen 2008), its epigenetic silencing can be reversed by heat stress, and its expression has been implicated in heat stress recovery (Sanchez and Paszkowski 2014). None of the genes that significantly differed in substitution rates were involved in flowering regulation. Because divergence from *A. montbretiana* is not independent between populations from the Cantabrian Mountains and from the Alps, this test can be considered conservative.

**Table 1:**
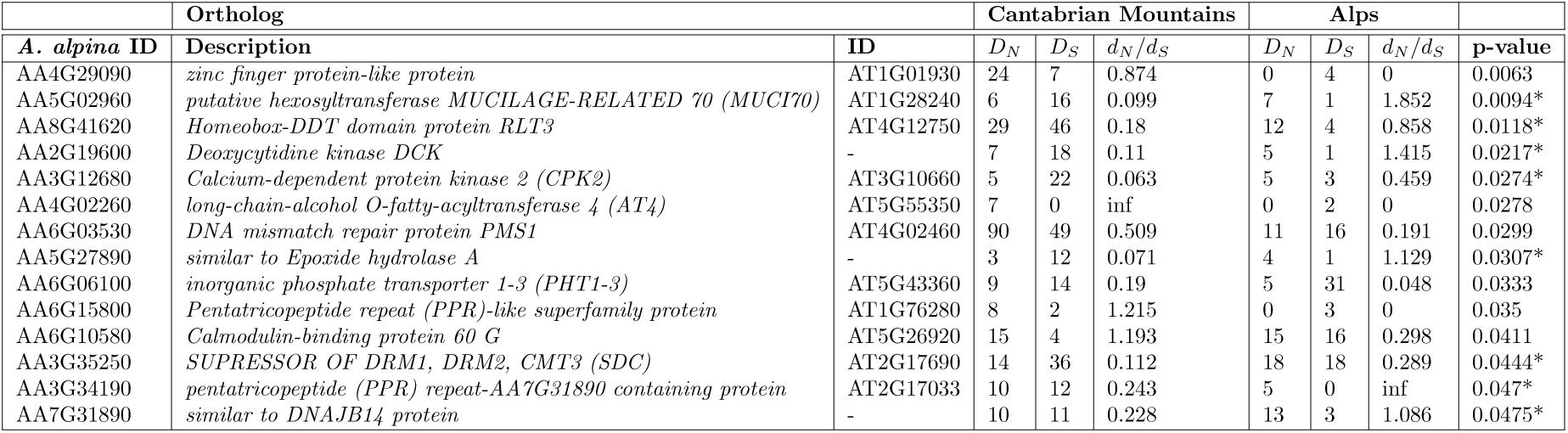
Genes with significantly different rate ratios of non-synonymous to synonymous substitutions in populations from the Cantabrian Mountains and from the Alps. The table shows the Gene Identifier in *A. thaliana* (ID), the number of derived non-synonymous (*D_N_*) and synonymous substitutions (*D_S_*), and their rate ratios (*d_N_* /*d_S_*) in the Cantabrian Mountains and in the Alps, sorted by p-value. Stars indicate lower *d_N_* /*d_S_* in the Cantabrian Mountains compared to the Alps.

Based on the rate ratios of non-synonymous to synonymous polymorphisms, we identified 481 genes with significant differences between the populations from the Cantabrian Mountains and from the Alps. The majority of these genes (426 genes, 88.57%), had a higher rate ratio in the Cantabrian Mountains than in the Alps, including the ten genes with the most significant differences. Their functions were related to abiotic stress and defense response, cell cycle control, and cell division (Table 2). The genes with significantly different rate ratios were enriched for GO categories related to protein phosphorylation, response to cyclopentenone, intermembrane phospholipid transfer, cell surface receptor protein serine/threonine kinase signaling, and regulation of endocytosis (Supplementary Table S8). Among the 41 genes associated with protein phosphorylation, 10 were also involved in cell surface receptor protein serine/threonine kinase signaling, and are primarily involved in stress- and defense-related signaling pathways. 40 of these 41 genes (97.6%) had higher rate ratios in populations from the Cantabrian Mountains than in populations from the Alps. The second most enriched GO category was response to cyclopentenone. Cyclopentenone-containing compounds such as OPDA (12-oxo-phytodienoic acid) regulate growth and various stress responses, including defense, and also serve as precursors to jasmonic acid (Yi et al. 2024). Eight genes were associated with this category, with primary functions in membrane stability and transport, cytoskeleton organization, and cell division and cytokinesis, and seven among them (87.5%) had a higher rate ratio in populations from the Cantabrian Mountains than in populations from the Alps. The genes involved in the negative regulation of endocytosis were two paralogs of *FT-INTERACTING PROTEIN 7* (*FTIP7*), which are located on two different chromosomes and have higher rate ratios in the Cantabrian Mountains compared to the Alps.

**Table 2:**
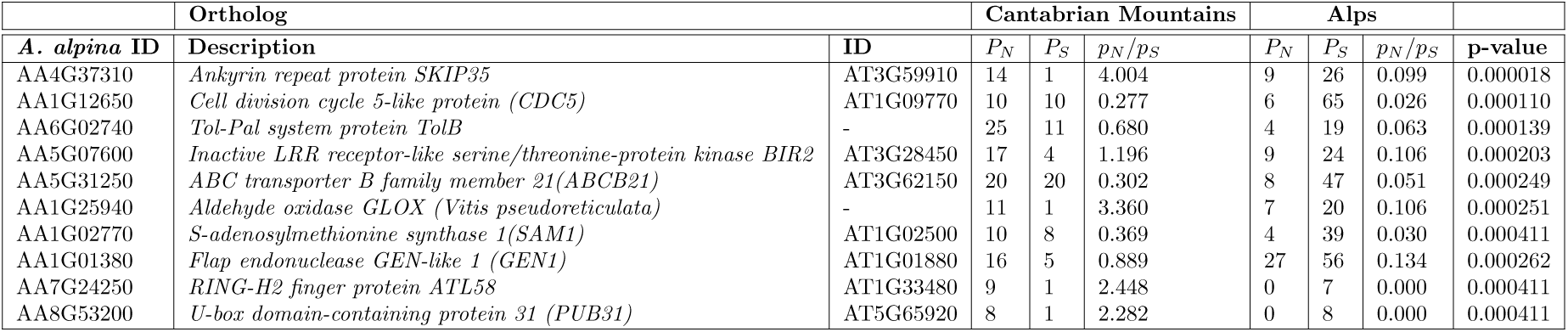
Top ten genes with the most significantly different rate ratio of non-synonymous to synonymous polymorphisms in populations from the Cantabrian Mountains and from the Alps. The table shows the Gene Identifier in *A. thaliana* (ID), the number of derived non-synonymous (*P_N_*) and synonymous polymorphisms (*P_S_*) and their rate ratios (*p_N_* /*p_S_*) in the Cantabrian Mountains and in the Alps, sorted by p-value.

Among the genes with significantly different rate ratios of polymorphisms, we identified as candidates that are potentially involved in flowering regulation the genes *FTIP7* (mentioned above), *CELL DIVISION CYCLE 5-LIKE* (*CDC5*), and *FRIGIDA-LIKE 1* (*FRL1*). *FTIP7* has been proposed to act redundantly with its homolog *FTIP1* in facilitating transport of FT into the phloem in multiple plant species, and FT positively regulates flowering (Orel et al. 2025). *FTIP7* has three paralogs in *A. alpina*, and two of them significantly differed in rate ratios (on chromosomes one: AA1G04680, and six: AA6G10170). At these two loci, we found a large number of derived alleles encoding semi- and non-conservative amino acid changes and presence-absence variants (Supplementary Materials and Supplementary Table S9). *CDC5* had the second most significant rate ratio among all genes.

In *A. thaliana*, *CDC5* is involved in transcriptional regulation, splicing, and epigenetic regulation of *FLC* and *MAF* genes, which are floral repressors, and mutations at *CDC5* accelerate flowering (Xin et al. 2025). *CDC5* had a total of 10 missense variants private to groups from the Cantabrian Mountains, which segregated at low frequencies (2.0%–13.2%), and which were mostly conservative or semi-conservative (8/10). It is possible that purifying selection is relaxed at *FTIP7* and *CDC5* in the Cantabrian Mountains, although we can not exclude that any of these variants may be functional. The candidate gene *FRL1* also significantly differed in the rate ratios of polymorphisms. In *A. thaliana*, FRL1 interacts with FRI as part of the FRI-C protein complex which contributes to the activation of *FLC* expression (Choi et al. 2011), until the plant experiences a cold treatment (vernalization), and FRI-C forms nuclear condensates (Zhu et al. 2021). *FRL1* had 21 and 15 non-synonymous substitutions respectively in the populations from the Cantabrian Mountains and from the Alps, compared to the outgroup *A. montbretiana*, and was therefore highly diverged. Near the C-terminus of the gene, we found a cluster of six non-synonymous mutations that were fixed derived in the Cantabrian Mountains compared to *A. montbretiana* (Fig. 5a). These mutations were linked into a haplotype that also segregated in the Alps at 23.7% frequency (Supplementary Table S10). To better understand the history of this polymorphism, we extracted from Relate the genealogies at and around *FRL1*. A single, very ancient genealogy overlapped most of this cluster of mutations (five out of six mutations; Fig. 5b). Divergence between haplotypes dated back to around 734 kya, and was therefore rather ancient. Finally, this genealogy was unusually balanced in comparison with neutral coalescence trees, and with genome-wide genealogies (Colless Index = 2933, empirical p-value=0.037). The genealogy that overlapped the sixth mutation was also balanced and ancient (Supplementary Fig. S7).

**Figure 5.**
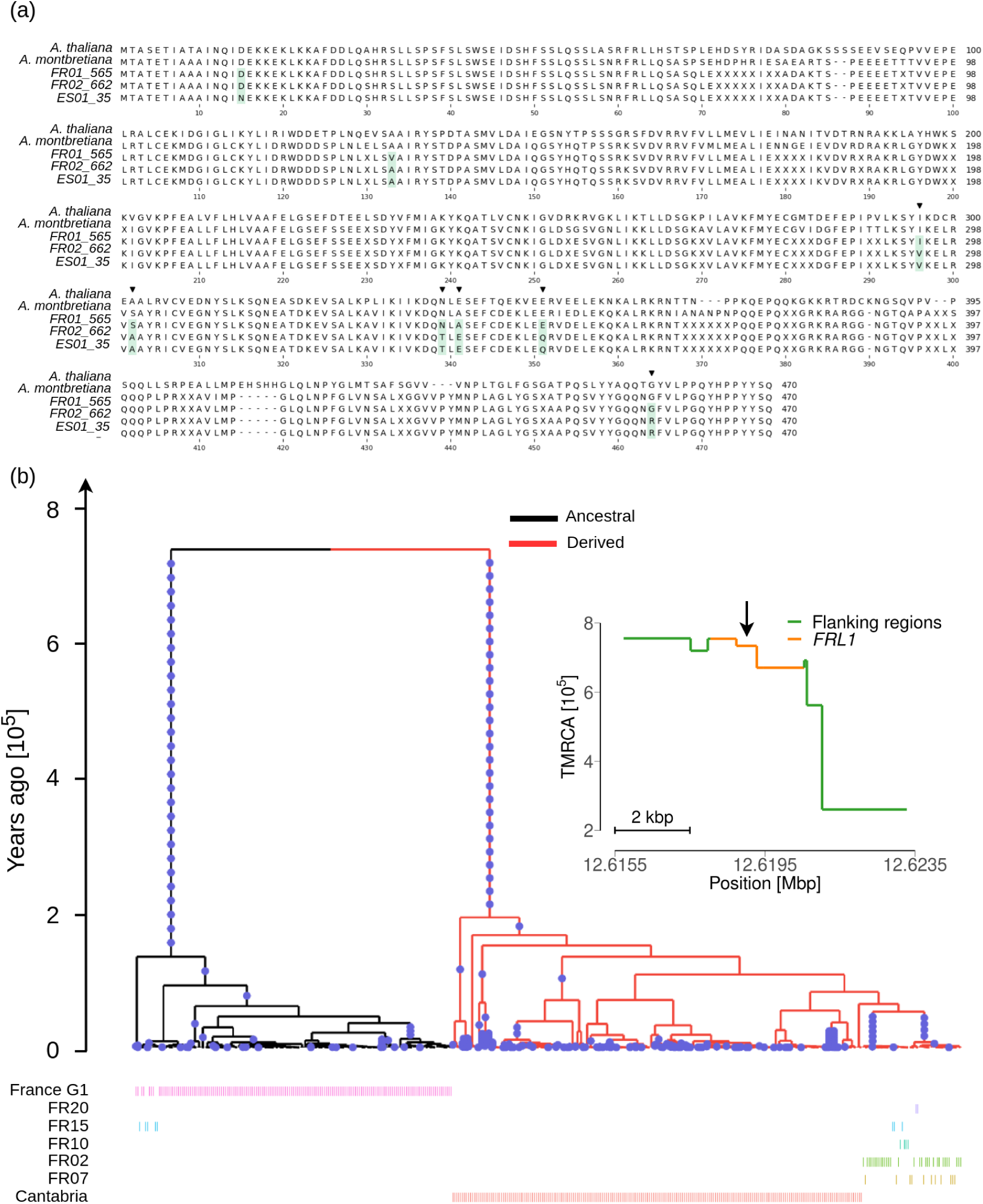
Genetic variation at the *FRL1* locus. **a:** Alignment of the FRL1 protein sequence from *A. thaliana*, *A. montbretiana*, and *A. alpina* from the Alps and from the Cantabrian Mountains. The six amino acid differences marked with black triangles are fixed derived in populations from the Cantabrian Mountains, and the two amino acid differences highlighted in green are segregating. **b:** The genealogy at the *FRL1* locus that spans five of the six non-synonymous mutations. Red branches indicate the lineages that carry the derived alleles at these five mutations, and black branches represent lineages that carry the ancestral alleles. France G1 (group 1) includes populations FR01, FR03, FR04, FR06, FR08, FR16, FR17, FR18, FR19, and FR21. The inset shows the time to the most recent common ancestor (tmrca) distribution across genealogies at the *FRL1* locus. The tmrca of the genealogy depicted here is marked by an arrow.

### Accessions from the Cantabrian Mountains flower late or are unable to flower without vernalization

Accessions from the Cantabrian Mountains were previously identified as outliers for flowering behavior across European populations of *A. alpina* (Wunder et al. 2023). Without vernalization, they were unable to flower within the course of the experiments, and they flowered for a short duration after vernalization. To confirm that the accessions used here had the same flowering behavior, we grew accessions from the Cantabrian Mountains and from the French Alps in controlled greenhouse conditions with and without a vernalization treatment (Supplementary Fig. S8). Consistent with previous results, most of the accessions from the Cantabrian Mountains were unable to flower without vernalization (65 accessions out of 71, or 91.6%, did not flower in any replicate). A single accession flowered in all four replicates, and five additional accessions flowered in at least one replicate, and they all flowered very late (on average 194.2 days from sowing). In comparison, all of the 55 accessions from the Alps flowered in at least one replicate, and most flowered in all replicates (51 accessions out of 55, or 92.7%). Accessions from the Alps varied widely between early and late onset of flowering (range: 48 to 247 days; on average after 115.8 days excluding non-flowering plants). After vernalization, all replicates of all accessions from the Cantabrian Mountains and from the Alps flowered, and did so mostly within 20-30 days from vernalization.

Consistent with an effect of the *FRL1* polymorphism on flowering regulation, the six non-synonymous mutations fixed in Spanish populations and segregating in the Alps were significantly associated with late flowering in this experiment (linear mixed-effect regression with four principal coordinates from genome-wide structure as covariates; Wald p-value = 3.96 ∗ 10*^−^*^3^; more detail in Supplementary Materials). Because the six mutations are in linkage disequilibrium with each other, these correlations are not independent, and represent a correlation of the entire haplotype.

## Discussion

Plants with an Arctic-alpine distribution are challenged by abiotic factors, such as extreme temperatures, low water availability, and short favorable seasons, and can reveal the dynamics of adaptation to abiotic stress. Moreover, these species experience habitat loss and fragmentation due to climate change, and studying their evolution and genomic diversity can reveal possible responses to a warming climate. In this study, we focused on the perennial, Arctic-alpine herb *A. alpina* and in particular on populations from the Cantabrian Mountains in Northern Spain, where the growing season is longer, warmer, and drier than in other regions of the range. Furthermore, these populations have been previously identified as outliers for flowering phenology, because they have a strong requirement for vernalization, and they flower for a short duration after vernalization (Wunder et al. 2023). In order to understand the evolutionary history of these populations and the adaptive value of flowering behavior, we characterized genomic diversity, we reconstructed their demographic history, and we analyzed genomic signatures of adaptation in comparison to outgroup populations from the Alps.

### *A. alpina* populations from the Cantabrian Mountains are a diverged genetic lineage that colonized Northern Spain during the last glacial period

The analysis of genomic diversity revealed strong differentiation between populations from the Cantabrian Mountains and from the French Alps, which was as ancient as 216 kya (198-230 kya) in our inference. This divergence dates back to the start of the penultimate glacial period between 194 and 135 kya, and is older than the most ancient splits species wide in the model plant *A. thaliana* (around 90 kya), and much older than its colonization of Eurasia (around 20 - 40 kya) (Durvasula et al. 2017). Therefore, in *A. alpina*, the Iberian Peninsula harbors a highly diverged genetic group, similar to the so-called Iberian relicts in *A. thaliana* (The 1001 Genomes Consortium 2016). However, the relicts in *A. thaliana* co-occur with other ancestry groups of different origin in Iberia (Castilla et al. 2020), while in *A. alpina* this single genetic group populated the Cantabrian Mountains. In the populations from the Cantabrian Mountains and from the Alps, historic effective population sizes have been mainly influenced by the last glacial episode, between approximately 115 and 12 kya. During this period, the Alps were covered by an ice shield (Ivy-Ochs et al. 2009) and the Iberian peninsula was a refugium for many species like bear, hedgehog, grasshoppers, and oak (Hewitt 2000; Petit et al. 2002). During this glacial episode, we detected a widespread increase in effective population size in populations from the Cantabrian Mountains and from the Alps. This suggests that the suitable habitat for *A. alpina* was larger at that time, and likely located at lower altitude, which may have aided the colonization of Northern Spain. Different from our results here, effects of glacial-interglacial cycles on genetic diversity pattern were not detected across seven tree species (Milesi et al. 2024).

Within the Cantabrian Mountains, we detected a pronounced pattern of isolation-by-distance, and marked differentiation among three genetic groups, found respectively in Central-Eastern Cantabrian Mountains (CE-CAN), and in Western Cantabrian Mountains around locality Angliru (W-CAN1), and around locality La Cueva (W-CAN2). Split times among populations aligned in an east-to-west gradient, with older splits for population pairs that include more eastern populations. Split times ranged between 150 kya and present, mostly overlapping the last glacial episode and the current interglacial. This suggests a colonization of the Cantabrian Mountains, or a range expansion, that possibly started from a region around the Alps around 200 kya, and progressed westward in Northern Spain through the last glacial period. These analyses cannot rule out that these populations may have inhabited other regions at the time they split from each other, and then shifted their range to the Cantabrian Mountains more recently, however the clear east-to-west cline in split times suggests a single range expansion wave. Our analyses do not detect any recent recolonization of the French Alps after the LGM from the Cantabrian Mountains, therefore the refugium of the ancestral population now inhabiting the Alps was probably located somewhere else. Finally, recent split times between populations in close proximity, and a widespread decline in effective population size in the recent past suggest that populations became increasingly fragmented since the LGM, corresponding to increasing average temperatures across Europe in the present interglacial.

### Candidate genes with signatures of positive selection have diverse physiological functions and may be involved in adaptation to abiotic stress

The demographic model that results from our inference suggests that the colonization of the Cantabrian Mountains happened during a glacial phase, when climatic conditions were likely colder and moister at lower altitude in Northern Spain, compared to present. In the current interglacial, since the last glacial maximum, temperatures have been rising on average, and especially in the Cantabrian Mountains the growing season has become longer and warmer, and summer has become dryer, compared to the rest of the European range. Because *A. alpina* is adapted to Arctic-alpine environments, the increasing temperatures in the current interglacial may have resulted in novel selective pressures. These new challenging conditions might have driven the evolution of new combinations of traits that together increased fitness in warm and dry conditions.

Enriched GO terms among genes in the top 5% CLR peaks and the candidate genes with the strongest selection signatures revealed three major functional associations: hormone regulation (e.g., *SC5D*, *GA20OX4* ; GO:0010358, GO:0010075), RNA metabolism and splicing (e.g., *BRR2C*, *SDN2* ; GO:0000381, GO:0045944), and lipid- and cell wall component metabolism (e.g., *KAS2*, *CASP*, *HPGT3* ; GO:0001676, GO:0035652, GO:0042538). This suggests that broad physiological pathways that might affect multiple plant traits are involved in adaptation. Furthermore, some ontologies were associated with selection signatures only on the ancestral, or on the terminal branches, and others were associated with both. The GO terms enriched on the ancestral branches included developmental processes (GO:0010075, GO:0046580, GO:0010358) and nutrient acquisition (GO:0071281, GO:0031670), different from the terminal branches on which multiple enriched GOs were involved in energy metabolism (GO:0001676, GO:0042775, GO:0009150). Both the ancestral and terminal branches were enriched for GO terms associated with abiotic stress responses. On the ancestral branches, these included responses to reactive oxygen species (ROS) (GO:0071452, GO:0046283), and on the terminal branches they included responses to hyperosmotic stress (GO:0042538), calcium-mediated signaling (GO:0050848, GO:1901019), which plays a key role in rapid stress responses, and long-chain fatty acid metabolism (GO:0001676), which is linked to cuticle biosynthesis. Additionally, enriched GO terms related to RNA metabolism included transcriptional initiation (GO:0051123, GO:0045944) on the ancestral branches, and mRNA splicing processes (GO:0000381) on the terminal branches. These results suggest that positive selection on the ancestral branches may have affected growth related phenotypes, nutrient responses, and adaptation to different light intensities, which might have been driven by shifts in altitude during glacial/interglacial cycles, and during the initial colonization of the Cantabrian Mountains. More recent selection in populations from the Cantabrian Mountains, however, likely involved adaptation to water deficiency.

Sixteen genes with selection signatures were directly associated with drought-related GO terms (response to water deprivation and heat). Among these, eleven were on terminal branches, and five on the ancestral branch, suggesting that selection on drought responses was recent. The most prominent candidate gene was *NAC055*, at which we found a 3bp insertion and a TE, which might change the function of the gene, and which are candidate genetic variants for adaptation to drought. The insertion was within a sequence motif at the C-terminus of the gene, which is shared by at least two other stress responsive NAC transcription factors (Tran et al. 2004), and which has been demonstrated to possess transactivation activity (Bu et al. 2008). Interestingly, *NAC055* and another candidate gene, *ZHD11*, are both induced by drought, salinity, and ABA, and overexpression lines have increased drought tolerance (Tran et al. 2004; Tran et al. 2007). Furthermore, they have been shown to cooperatively bind to and induce transcription of the gene *EARLY RESPONSIVE TO DEHYDRATION STRESS 1* (*ERD1*), which is downstream in the ABA independent drought response pathway (Tran et al. 2007). This suggests that positive selection may target specific transcription factor hubs at different regulatory nodes.

Finally, selection signatures were associated with many more genes with various functions, which suggests that selection was multivariate, and that various other traits may have been involved in adaptation. All the genes and genetic variants mentioned here are candidates that might have been involved in adaptation to climatic changes, however the causative mutations and their adaptive value remain to be validated.

### An ancient polymorphism at *FRL1* is a candidate for conferring variation in flowering behavior

Accessions from Cantabrian Mountains have been previously identified as outliers for flowering phenology (Wunder et al. 2023). These accessions have a strong requirement for vernalization to flower, and they flower for a short duration after vernalization, compared to other European populations that largely vary in flowering behavior (Wunder et al. 2023). Because a functioning vernalization pathway was likely present in ancestral populations in Brassicaceae, it is possible that the strong vernalization requirement is ancestral, and may have remained conserved in the Cantabrian Mountains through stronger stabilizing selection than in other European regions. Positive selection may have further contributed to the evolution of this trait, by strengthening the ancestral vernalization requirement. Consistent with selection on flowering regulation, four genes in the vernalization pathway that regulate *FLC* expression, *VIP4*, *VIL1*, *FRL1*, and *CDC5*, and one gene in the photoperiod pathway, *FTIP7*, have emerged as candidates to regulate flowering in the Cantabrian Mountains.

At *FRL1*, we detected very ancient divergence between two alternative haplotypes with a cluster of six non-synonymous mutations near the C-terminus of the gene. One of the haplotypes was fixed in Spain and it segregated at low frequency in the Alps, which suggests a local, soft sweep from standing genetic variation in the Cantabrian Mountains. In *A. thaliana*, *frl1* mutants flower very early, similar to *fri* mutants (Michaels et al. 2004). FRL1 is thought to stabilise the FRI-C complex, and it binds to FRI at its N-terminus, and to SUF4 (Choi et al. 2011), possibly at its C-terminus. *SUF4* binds to the promoter of *FLC* and likely has a crucial role on regulating *FLC* expression and flowering.

After a cold treatment, FRI-C forms nuclear condensates and is sequestered from the *FLC* promoter, and flowering is enabled (Zhu et al. 2021). Because we did not observe any high-impact mutation at *FRL1*, we hypothesize that the two alternative alleles are functional, and that they may differ in their effect of stabilizing FRI-C. Consistent with an effect of *FRL1* on flowering regulation, also the plants from the Alps with the *FRL1* haplotype found in the Cantabrian Mountains flowered late without vernalization (FR02: 171.67 ± 29.27 days; FR07: 178.85 ± 34.24 days; FR10: 220.0 ± 20.78 days), similar to populations from the Cantabrian Mountains. Conversely, plants with the haplotype that is prevalent in the Alps varied largely in flowering behavior, from early to late flowering, which was likely due to other genetic variants. Overall, the polymorphism at *FRL1* was very ancient, its origin largely predated the colonization of Cantabrian Mountains (at most as ancient as the split from populations in the Alps, 198-230 kya), and the genealogies associated with it were more balanced than expected, with long internal branches and short external branches. These results suggest that variation in flowering behavior evolved under balancing selection in perennial *A. alpina*, perhaps because of spatially or temporally varying selection, mediated by alternative, functional alleles at *FRL1*. In contrast, in *A. thaliana* and other annual species, the adaptive value of flowering regulation is often associated with positive selection for early flowering mediated by nonsense mutations at *FRI*, and rarely at *FLC*, which confer an escape strategy (Austen et al. 2017; Johanson et al. 2000; Méndez-Vigo et al. 2011; Fulgione et al. 2022; Stinchcombe et al. 2004; Zhang and Jiménez-Gómez 2020).

### A strong vernalization requirement and short flowering duration contribute to adaptation to summer drought in a perennial plant

The adaptive value of flowering regulation is established for annual plants, but it may differ in perennials. Perennial plants, like annuals, transition from vegetative growth to flowering, and they can vary in whether they flower in the first growing season, or only after vernalization. However, perennials do not senesce after flowering, and they can revert to vegetative growth and arrest flowering (Wang et al. 2009). This reversion is mediated by cycles in the expression of genes that can have pleiotropic effects on both the time of flowering onset and the time of flowering arrest, and therefore on the duration of flowering, such as *PEP1*, *MAF8.1*, and *MAF8.3* (Wang et al. 2009; Madrid et al. 2021), as well as genes cycling in opposition, such as *COOLAIR* and *VIN3* (Castaings et al. 2014). Different from annuals, flowering arrest in perennial plants can be linked to a trade-off between maximizing the reproductive output in the current growing season and maximizing survival until the next growing season and future reproduction. A negative correlation between the time of flowering onset and flowering duration has been observed across plants (Hendry and Day 2005), and specifically in *A. alpina* (Wunder et al. 2023). It is therefore unclear to which extent these traits evolve under direct selection or due to selection on correlated traits. Here, we argue that a strong vernalization requirement and a short duration of flowering align into a strategy characterized by flowering avoidance during summer. In Arctic-alpine environments, seeds can germinate earliest after snow melt, which happens around May at Spanish sites and around June to July in the Alps and Scandinavia, based on climatic data recorded at the sites where *A. alpina* grows (Wunder et al. 2023). The growing season (here defined as the snow-free season) is about six months long in Spain, and around four months long in the Alps and Scandinavia. An exception is the site near Granon in the Alps (F2 in (Wunder et al. 2023), and FR02 and FR07 here), where the snow melts around May and the growing season is around six months long, similar to Spanish sites. Plants with a weak (or without any) vernalization requirement can flower a minimum of around seven to eight weeks after germination in greenhouse conditions (Wang et al. 2009; Albani et al. 2012; Wunder et al. 2023) and likely later than that in the field. In nature, these plants could therefore flower in the first growing season around mid- to late summer (July to August in Spain, and August to September in the Alps and Scandinavia). Plants with a strong requirement for vernalization are unlikely to flower in the first year, especially when the growing season is short, as in Arctic-alpine environments. As a consequence, they effectively avoid flowering in mid- to late summer. In the following years, all plants are fully vernalized and they readily flower after winter. Plants with a weak requirement for vernalization also flower for a long duration, or perpetually, after vernalization, therefore they can flower until the end of the growing season. Conversely, plants with a strong vernalization requirement have a short flowering duration and arrest flowering after approximately 40-60 days in greenhouse experiments (Wunder et al. 2023). At the Spanish sites, these plants would arrest flowering around July, which prevents flowering after mid- to late summer also in the growing seasons that follow the first one. We note that genotype by environment (GxE) interactions may affect these scenarios, however the short duration of flowering of accessions from the Cantabrian Mountains was previously confirmed also for plants growing *in situ* (Wunder et al. 2023), and during our visits at the field sites. Overall, variation in flowering behavior in a perennial plant results in alternative seasonal strategies that mainly differ in whether plants flower in mid- to late summer or not, both in the first growing season and in the following years. When summers are long, warm, and relatively dry, as in Cantabrian Mountains, plants that avoid flowering through the summer may reduce water stress and escape drought. Interestingly, at the site near Granon in the Alps, the climatic conditions are similar to Spanish sites, accessions collected there regulate flowering in a similar way as Spanish accessions, and the *FRL1* haplotype found in Spain is also locally fixed. Consistent with a fitness cost of flowering through a dry summer, flowering in the first year was associated with an increase in mortality rate in the Cantabrian Mountains, but not in the Alps or in Scandinavia (Wunder et al. 2023). In annual plants, dehydration due to drought is escaped by reducing water consumption to a minimum, and surviving as seeds through the summer. In a perennial, avoiding processes that require high water consumption, like flowering, during the dry season may be close to a dehydration escape or avoidance strategy. Dehydration escape in the sense of completing the reproductive phase as early in the season as possible in water-limited environments, and avoidance in the sense of minimizing water loss (Volaire 2018). Counter-intuitively, while the principal escape strategy in annuals is associated with early flowering, the closest counterpart in a perennial is associated with late flowering and with a strong vernalization requirement.

### Concluding remarks

The study of plants with an Arctic-alpine distribution can contribute to our understanding of how plants adapt to abiotic stress, and can reveal possible responses to a warming climate. Our study reveals ancient divergence in the perennial, Arctic-alpine herb *A. alpina* from the Cantabrian Mountains in Northern Spain, and an east-to-west range expansion that started around 200 kya from the Alps and proceeded with the colonization of Northern Spain during the last glacial period. Since the LGM, in the current interglacial, we find evidence of population fragmentation, likely mediated by warming temperatures that resulted in selective pressures for the evolution of a drought-related trait syndrome, especially in the Cantabrian Mountains where the climate is warm and dry in summer. Among the genes with a signature of selection involved in drought responses and growth, we identified *SC5D* and *NAC055* as the strongest candidates for future molecular validation. Furthermore, we identified an ancient polymorphism at the gene *FRL1* which may contribute to variation in vernalization requirement and flowering duration. We argue here, that the combination of a strong vernalization requirement and a short duration of flowering in *A. alpina* populations from the Cantabrian Mountains results in drought escape, by avoiding increased water stress due to flowering during the warm, dry summer. Conversely, plants from other regions of the range vary between early, late, and no onset of flowering, possibly because of relaxation of purifying selection, or because of spatially or temporally varying selection. Resolving the adaptive value of variation in flowering behavior in these populations is an avenue for future research. Overall, our study suggests that the combination of two different evolutionary processes, positive selection at drought response genes and the maintenance of ancestral variation in flowering phenology, is a possible avenue for the evolution of new, synergistic trait combinations, or new trait syndromes, which can result in functional ecological strategies.

## Materials and Methods

### Plant material, short reads sequencing and variant calling

We collected cuttings and seeds from 211 individual *A. alpina* plants across 14 populations in the Cantabrian Mountains in Northern Spain, and from 215 individual plants across 15 populations in the French Alps (Supplementary Table S1). Some of these accessions were sourced from previous studies (Wunder et al. 2023; Tjeng et al. 2024). We extracted DNA and sequenced whole genomes with 150 bp paired end short reads nano ball sequencing at the Beijing Genomics Institute (BGI). More details on DNA extraction and sequencing can be found in the Supplementary Materials.

SNP calling was performed using the GATK pipeline *v*4.2.0 (Depristo et al. 2011). Read groups were assigned, adapters were soft clipped, duplicates were marked and reads were aligned to the reference genome *Arabis alpina v5.1* (Jiao et al. 2017). GATK haplotypecaller was used to call SNPs and short indels by performing local realignments. To avoid biases in variant frequencies, genotypes were called for each sample separately. Genotypes with genome-wide average depth of 10x or less were excluded from the analyses. SNPs with depth less than 5x and genotype quality less than 30 were set to missing values. Indels were removed and only biallelic sites were retained. We identified and filtered multicopy genomic regions in the samples from the Cantabrian Mountains and from the Alps, after filtering sites with more than 30% missing values, using default settings in ParaMask (Tjeng et al. 2024). Furthermore, we retained only uniquely mapping genomic regions inferred with SNPable (https://lh3lh3.users.sourceforge.net/snpable.shtml), using K-mer length of 150 and intermediate stringency (r=0.5).

### Genomic diversity and population structure analyses

We obtained a set of unrelated individuals by applying a greedy search algorithm to the empirical distribution of pairwise differences, in comparison to expected values assuming known relatedness status (detail in Supplementary Materials). We calculated *θ_π_*, *θ_W_*, Tajima’s D, and *F_IS_* on the populations with sample sizes greater than four using custom scripts available in GitHub (details in Supplementary Materials). To quantify isolation by distance (IBD), we calculated the Pearson product moment Mantel correlation between the matrix of pairwise differences and the matrix of geographical distances among samples using 10,000 bootstrap replicates. For the principal coordinate analysis (PCoA), we performed ordination of 1 - IBS distances in a 2-dimensional space using the metric scaling implemented in PLINK *v*1.9.0 (Chang et al. 2015). We inferred the proportion of shared genetic ancestry for two to 20 ancestry groups using ADMIXTURE *v*1.3.0 (Alexander et al. 2009), after pruning singleton SNPs, SNPs with more than 10% missing genotypes, SNPs in linkage disequilibrium (*r*^2^ *>* 0.2), and related individuals. The best number of ancestry groups was chosen based on the lowest average cross-validation (CV) error among 10 replicates (details in Supplementary Materials).

### Demographic inference

To reconstruct genome-wide genealogies and calculate coalescent rates, we used the software relate *v.*1.2.2 (Speidel et al. 2019). We inferred the ancestral states for genome-wide SNPs on the basis of an alignment between the *A. alpina* reference genome, and the genome of the outgroup species *A. montbretiana* (described in Supplementary Materials). We imputed and phased genomic data in two steps (described in Supplementary Materials), and we pruned annotated genes and their flanking regions (2kbp). We accounted for missing data and preserved the SNP densities by adjusting the distances between SNPs, and we accounted for inbreeding by selecting randomly one haplotype per individual. We assumed a mutation rate of 7 × 10*^−^*^9^ (Ossowski et al. 2010), a generation time of 1.5 years per generation (Laenen et al. 2018), an initial effective population size of 2 × 10^5^, and we used a scaled recombination map from crosses (Supplementary Materials). Confidence intervals were constructed from 100 bootstraps, by randomly re-sampling 5 Mbp regions, and constructing pseudo chromosomes of the same size as the actual chromosomes. For the inference of split times, we used a threshold of 0.5 for the relative cross-coalescent rate, after fitting a piecewise cubic polynomial function to the point estimates and 95% confidence intervals on a logarithmic time scale using the *smooth.spline* function from the stats *v*4.0.4 package in R.

### Identification of genomic signatures of selection with *3P-CLR*

We identified signatures of positive selection with *3P-CLR* (Racimo 2016). As focus clades, we used different combinations of the three major clusters from the Cantabrian Mountains, W-CAN1, W-CAN2, and ES17 (as a non-admixed representative of CE-CAN, details in Supplementary Materials). We used as outgroups the pooled populations from the Alps, and CE-CAN (details in Supplementary Materials). For each combination of clades, we excluded sites with more than 10% missing values within groups. We calculated drift rates on the basis of F3 statistics after pruning multicopy regions, regions that do not map uniquely inferred with SNPable, and regions two kbp up- and down-stream of genes. We calculated CLR scores in genomic windows of 0.00025 Morgans, including genes and multicopy regions, and excluding SNPs with a minor allele frequency smaller than 0.01 in the outgroup. The number of sampled SNPs was set to 100, with a minimum distance of 20 bp among them. CLR scores where normalized by mean and standard deviation for every branch and set of samples. The 0.5% highest CLR peaks in every branch were used for Gene Ontology enrichment analysis and the genes that overlapped the top five peaks of every group of branches were manually analyzed. For the analysis of candidate genes, we additionally reconstructed genealogies including annotated genes with relate using the same parameters described above.

### Inference of synonymous and non-synonymous divergence and polymorphism

Prior to the inference of molecular evolution statistics, we improved the annotation of the reference genome, and we predicted the effects of genetic variants (described in Supplementary Materials). We quantified the numbers of non-synonymous and synonymous substitutions (*Dn*, *Ds*), and their rate ratio (*d_N_ /d_S_*), and polymorphisms (*Ps*, *Ps*) and their rate ratio (*p_N_ /p_S_*), across all 28745 annotated genes in the genome using custom scripts available in GitHub. To calculate the total number of n-fold degenerate sites per gene, we used degenotate (Mirchandani et al. 2023). We compared the rate ratios in the populations from the Cantabrian Mountains and from the Alps using Fisher’s exact test.

### Gene ontology (GO) enrichment analyses

The enrichment of GO categories among genes with signatures of selection were tested using topGO (Alexa et al. 2006) with the elim algorithm and Fisher’s exact test. We tested the set of genes that overlapped the 0.5% high tail of *3P-CLR* for multiple combinations of branches and analyses (terminal or ancestral branches, and using populations from the Alps or from CE-CAN as outgroup), and the set of genes with a significantly different *p_N_ /p_S_* in populations from the Cantabrian Mountains and from the Alps. We only considered GO categories that included a minimum of two genes and more than one locus with signatures of positive selection, and an enrichment with a minimum significance level of 0.5% without multiple hypothesis testing.

### Data and resource availability

All code used in data analyses and visualization is available in GitHub at: https://github.com/Fulgione-group/Spanish_Arabis_Alpina. The sequencing dataset supporting the conclusions of this article is available in the European Nucleotide Archive (ENA) under the accession numbers PRJEB73825 and PRJEB96335. The new annotation of the reference genome is available at http://www.arabis-alpina.org/. All data supporting conclusions, that are not included in the Supplementary Materials and scripts for analyses are available in Zenodo at: https://zenodo.org/records/17053709?token=eyJhbGciOiJIUzUxMiJ9.eyJpZCI6IjZhMDQ5Mjg[]Nl87OvIgjKtTIabbsI-cqVuunoBoVSPEr3gzzVy0W054TJBOKdqCT03PjYPx7LA.

## Author contributions

A.F., G.C. and B.T. conceived and designed the study. B.T. performed most of the analyses. M.M. performed the *d_N_ /d_S_* and *p_N_ /p_S_* analyses. H.B.G. performed the greenhouse experiment. J.Z. annotated the reference genome. C.A.B., A.D.L., B.T. and J.W. collected the plants, A.D.L. and B.T. propagated them and extracted DNA for sequencing. B.T. and A.F. wrote the manuscript, with contributions from all authors.

## Acknowledgements

This study was supported by funding from the Deutsche Forschungsgemeinschaft, DFG, German Research Foundation (Project-ID 514187363 - FU 1387/2-1, and Project-ID 456082119 - TRR 341/1), and by funding from the Max-Planck-Gesellschaft, all to AF. Support for plant growth and for computational analyses were provided by the Max Planck Institute for Plant Breeding Research (MPIPZ). The CAB laboratory was funded by grant PID2022-136893NB-I00 from the MCIN / MCIU / AEI / 10.13039/501100011033 and FEDER (EU).

