## Supplementary Material for "Adaptation to seasonal drought in *Arabis alpina* is linked to the demographic history and climatic changes since the last glacial maximum"

September 8, 2025

[1]: Max Planck Institute for Plant Breeding Research, Carl-von-Linne-Weg 10, 50829, Cologne, Germany

[2]: Nunhems Netherlands BV, Napoleonsweg 152, 6083 AB Nunhem, The Netherlands

[3]: Centro Nacional de Biotecnologia (CNB), Consejo Superior de Investigaciones Científicas (CSIC), Madrid 28049, Spain

### Supplementary Text

#### Plant material, short reads sequencing and variant calling

We collected cuttings and seeds from 211 individual *A. alpina* plants across 14 populations in the Cantabrian Mountains in Northern Spain, and from 215 individual plants across 15 populations in the French Alps (Supplementary Table S1). Some of these accessions were sourced from previous studies (Wunder et al. 2023; Tjeng et al. 2024). We rooted the cuttings in the greenhouse, and we extracted DNA from 100 mg of young leaves and meristems, following the DNeasy Plant Mini Kit protocol with minor adjustments. Whole-genome sequencing with short reads was performed at the Beijing Genomics Institute (BGI) for all samples. DNA concentration was quantified by fluorometer or microplate reader (e.g. qubit fluorometer, invitrogen). Sample integrity and purity were detected by agarose gel electrophoresis (concentration of agarose gel: 1%, voltage: 150 V, electrophoresis time: 40 minutes). 1 $\mu g$ of genomic DNA was randomly fragmented with covaris. The DNA fragments were selected by magnetic beads to an average size of 200-400 bp. Fragments were end repaired and then 3' adenylated. Adaptors were ligated to the ends of these 3' adenylated fragments. Fragments were amplified with PCR using the adaptors from the previous step. PCR products were purified by the magnetic beads. The double stranded PCR products were heat denatured and circularized by the splint oligo sequence. The single strand circle DNA (ssCir DNA) were formatted as the final library. The library was amplified with phi29 to make DNA nanoball (DNB). The DNBs were load into the patterned nanoarray and paired-end 150 base pairs reads were generated with combinatorial probe-anchor synthesis (cPAS).

#### Analysis of genetic diversity and population structure

To obtain a set of unrelated individuals, we calculated the average pairwise difference for any second-degree relationship ($C_{pd}$). This calculation was based on the average within population pairwise differences ($\overline{pd}$) and mean heterozygosity ($\overline{het}$), assuming an outcrossing rate of $t = 0.2$, which reflects conservative estimates from mother-offspring relationships reported by Toräng et al. (2017). Considering a grandmother-grandchild relationship, with either two generations of selfing, one generation of selfing followed by outcrossing (or vice versa), or two generations of outcrossing, the formula is as follows:

$$C_{pd} = (1-t)^2 \cdot 0.5 \cdot \overline{het} + 2t(1-t) \cdot \big(0.25 \cdot \overline{het} + 0.5 \cdot \overline{pd}\big) + t^2 \cdot \big(0.125 \cdot \overline{het} + 0.75 \cdot \overline{pd}\big)$$

We then applied a greedy search algorithm to retain pairs of individuals with pairwise differences higher than that estimate.

To estimate diversity and $F_{IS}$, we used the set of unrelated individuals, and we did not prune for linkage. To calculate $F_{IS}$, we used populations with sample sizes ($\geq 10$) to avoid high error rates in allele frequency estimates, and we calculated Nei's $Ho$ and $Hs$ per site, across populations (Nei 1987). We then calculated $F_{IS}$ genome-wide and in windows of 10,000 SNPs as $1 - \sum Ho / \sum Hs$. To account for missing values, we randomly projected the number of genotypes down to 90% of the total sample size per population.

In the ADMIXTURE analysis, the most representative replicate for each number of ancestry groups, and the best alignment of clusters between the most representative replicates was chosen after post processing the Q-matrices with pong (Behr et al. 2016).

#### Identification of ancestral states based on a whole-genome alignment of the outgroup *Arabis montbretiana* to the *A. alpina* reference genome

We aligned the *Arabis montbretiana* genome, accessed at NCBI with accession number PRJNA258048, to the *A. alpina* reference genome *v*5.1 (Jiao et al. 2017) using lastz aligner *v*1.04.15 (Harris 2007), kentools *v*302 (Nassar et al. 2023) and GenomeAlignmentTools (Sharma and Hiller 2017; Osipova et al. 2019). Each chromosome of the *A. montbretiana* reference was split in ten segments, and each segment was aligned to the *A. alpina* reference. We used lastz parameters $-gap = 400, 30$, $-gappedthresh = 3000$, and $-inner = 2200$. The alignments of each segment were chained using axtChain from kentools with parameter $-linearGap = medium$. To fill missing alignments in repetitive regions we used the RepeatFiller script from GenomeAlignmentTools with lastz parameters $O = 400$, $E = 30$, $L = 3000$, $T = 1$ and $H = 2200$. The patchChain script in GenomeAlignmentTools was used to perform highly sensitive local realignment after chaining. Segments before and after realignment were merged, chained again and netted. The axt alignment files were converted into a fasta file with the sequence coordinates of the *A. alpina* reference genome.

#### Demographic inference

For demographic analyses, we phased genomic data in two steps. First, we prephased haplotypes using information of sequencing read pairs in the bam files with WhatsHap *v*.1.1 (Martin et al. 2023). Next, the data set was divided into two subsets, one for the Cantabrian Mountains populations and one for the populations from the Alps. For each subset we pruned sites with more than 20% missing genotypes, and multicopy regions identified with ParaMask (Tjeng et al. 2024). We then jointly phased the data, and imputed missing genotypes on each sample set using the HMM-based approach implemented in shapeit *v*.4.2 (Delaneau et al. 2019). In shapeit, we used recombination maps from a F2 crossing population between individuals from population FR01 (Wötzel 2016). Recombination rates were scaled to $(1 - F_{IS})$ to adjust for inbreeding (Nordborg 2000), using the average $F_{IS}$ that we inferred in previous analyses. We then merged the subsets and pruned annotated genes and their flanking regions (2kbp).

#### Acquisition of climatic data

Monthly temperature and precipitation records were obtained from https://www.worldclim.org/ at a resolution of 30" for the period 1970–2000 for all collection sites in the Cantabrian Mountains and in the Alps, as well as for sites in Scandinavia described in Wunder et al. (2023). The monthly Aridity Index at the collection sites was sourced from Zomer et al. (2022).

#### Improvement of the annotation of the *A. alpina* reference genome

We constructed an NCBI assembly BLAST database using rmblast *v*2.10.0+ and performed a *de novo* search for repetitive elements with RepeatModeler *v*2.0.1 (Flynn et al. 2020). We hard-masked repeats longer than 5bp, and we soft-masked shorter ones using RepeatMasker (http://www.repeatmasker.org). The structural annotation was generated using Maker *v*3.01.03 with the proteome of *A. alpina* (UniProt identifier: UP000029120), and of *A. thaliana* (UniProt identifier: UP000006548), as well as a previous annotation supplied as evidence files (Cantarel et al. 2008). The gene predictor Augustus was used as part of the pipeline, with the *A. thaliana* data set as HMM training set. Additionally, we employed the deep neural network gene prediction software Helixer $v0.3.0$ (Stiehler et al. 2021; Holst et al. 2023). Post-processing of the gene annotations was conducted with the AGAT suite $v1.0.0$ (Dainat 2023), which resulted in manually curated GFF3 files. To further enhance the annotation, we integrated homology information using Orthorbb $v2.2$ from GenomeAnnotation (`https://github.com/darencard/GenomeAnnotation`) with default parameters. Functional annotation, including gene ontologies, was performed using EggNOG $v2.1.9$ with default settings (Huerta-Cepas et al. 2019; Cantalapiedra et al. 2021).

To predict the effects of genetic variants, we first constructed a novel database for the reference genome by utilizing the new genome annotation GTF file. Then, we annotated a VCF file that included outgroup data from the whole-genome alignment of *A. montbretiana* for SNPs, and outgroup data from short reads of *A. montbretiana* for indels, using the newly created database described above, and the default settings in SNPeff $v5.2.a$ (Cingolani et al. 2012).

#### Selection scans with *3P-CLR*

For the *3P-CLR* analysis, we used six combinations of populations, each consisting of two subject populations and one outgroup (Supplementary Table S3). As subject populations, we used the three Cantabrian Mountains structure groups, combined in all possible pairs (three combinations in total), and used the pooled French population as the outgroup. However, the resulting negative drift rate estimates between CE-CAN and the W-CAN1 or W-CAN2 groups (–0.066 and –0.063, respectively) indicated potential admixture, violating the tree like assumptions of the method. To address this, we replaced the pooled CE-CAN group with only the westernmost CE-CAN population, ES17, which is genetically closest to the W-CAN groups and also the most distinct within CE-CAN. This adjustment resulted in positive drift rate estimates between ES17 and both W-CAN groups (Supplementary Table S3). Given the high divergence between Cantabrian Mountains and French populations, we also performed additional *3P-CLR* analyses using CE-CAN as the outgroup. Specifically, we used the full CE-CAN group, including ES17, as outgroup for the analyses focused on W-CAN1 and W-CAN2, and we used CE-CAN excluding ES17 as the outgroup when analyzing combinations that included W-CAN1 or W-CAN2 with ES17 as subject populations.

#### Correlations between allele dosage at the 3bp insertion at *NAC055* and the aridity index

After we identified a 3bp insertion at *NAC055* as a candidate genetic variant with a selection signature and possibly related to drought responses, we tested whether this variant was correlated with the aridity index at the collection site. We repeated the analysis on the full data set (n=211), and after excluding populations ES02, ES10, and ES15 (n = 200), which were collected at sites with micro climates very distinct from the surrounding climate. In particular, ES02 was on a roadside and shaded by buildings in an otherwise warm and dry environment, ES10 was inside a cave, likely moister and cooler than the surroundings, and ES15 was on a roadside and near buildings in a canyon near a small river.

The aridity index in July was a highly significant predictor of allele dosage at the 3bp insertion, based on a binomial regression with four principal coordinates from genome-wide structure as covariates (on the whole data set, Wald p-value $= 6.26 * 10^{-5}$; excluding populations with putatively distinct microclimate, Wald p-value $= 5.14 * 10^{-4}$; Supplementary fig. S5). Similarly, the allele dosage at the 3bp insertion was a highly significant predictor of the aridity index in July, based on a linear regression with four principal coordinates from genome-wide structure as covariates (on the whole data set, Wald p-value $= 4.48 * 10^{-9}$; excluding populations with putatively distinct microclimate, Wald p-value $= 5.78 * 10^{-15}$).

#### Characterization of the mutations at *FTIP7*

Two of the three paralog copies of *FTIP7* had a significantly different rate ratio of polymorphisms between populations from the Cantabrian Mountains and from the Alps. Specifically, at AA1G04680, we found four non-conservative amino acid changes and four in-frame deletions and duplications segregating in Cantabrian Mountains and mostly fixed for the ancestral allele in France (Supplementary Table S9). At AA6G10170, we found in total five mutations with semi- or non-conservative effects, of which four were segregating in Cantabrian Mountains and fixed in French populations, and one was segregating in both regions (Supplementary Table S9).

#### Greenhouse experiment

We grew 71 accessions from the Cantabrian Mountains and 55 from the Alps in a controlled greenhouse experiment to characterize their flowering behavior. Accessions were grown in four replicates with a randomized block design, with one replicate per block, and with a photoperiod of 16 h of light, and 8 h of darkness. Seeds were stratified in cold and moist conditions for five days, then planted in 9x9 cm pots with a standard soil mix. Germination time was uniform across accessions. The time of flowering onset (flowering time) was scored as the number of days between sowing and the opening of the first flower, and was scored twice per week. Plants that had not flowered by the end of the experiment at 250 days after sowing, were considered non-flowering. Plants that died before flowering were excluded from the experiment. Accessions for which one replicate died were included in the analyses, and for no accession more than one replicate died in the experiment.

#### Correlations between *FRL1* haplotype and flowering time

We tested for an association between the six non-synonymous substitutions at the C-terminus of *FRL1* and flowering time without vernalization scored in the greenhouse experiment on all the 126 accessions from populations in the Cantabrian Mountains and in the Alps. For this, we fitted a linear mixed model in the R package lme4 *v*1.1.28, using the allele dosage as predictor, four principal coordinates as covariates, and the accessions as random effect. Fixed effects were tested with an analysis of deviance using the Wald chi-squared test. For the four principal coordinates, we used the multidimensional scaling function in PLINK, with 1-IBS distances on genome-wide SNPs. Prior to this, SNPs with more than 10% missing genotypes, unmappable SNPs identified by SNPable, and multicopy regions identified with ParaMask in the populations from Northern Spain and from the Alps were filtered out. Flowering time of individuals that did not flower by the end of the experiment was set to 250 days (the end of the experiment).

### Supplementary Tables

**Table S1:** *Arabis alpina* accessions used in this study and their metadata, including sample ID, country, location, coordinates, collection date, and Source (CL = Coupland lab; FL = Fulgione Lab; CAB = Carlos Alonso-Blanco). The sample ID is composed of the population ID, accession number, a letter indicating collection as cuttings (C) or seeds (S), and the number of generations selfed before sequencing.

| ID | Country | Location | Coordinates | Collection date | Source |
|---|---|---|---|---|---|
| ES01-001-C-00 | Spain | Léon - Puerto de la Cubilla | N42.9931°, W5.9100° | 8/9/2020 | FL |
| ES01-001-S-00 | Spain | Léon - Puerto de la Cubilla | N42.9931°, W5.9103° | 8/9/2020 | FL |
| ES01-004-C-00 | Spain | Léon - Puerto de la Cubilla | N42.9935°, W5.9102° | 8/9/2020 | FL |
| ES01-005-C-00 | Spain | Léon - Puerto de la Cubilla | N42.9929°, W5.9094° | 8/9/2020 | FL |
| ES01-007-C-00 | Spain | Léon - Puerto de la Cubilla | N42.9932°, W5.9103° | 8/9/2020 | FL |
| ES01-009-C-00 | Spain | Léon - Puerto de la Cubilla | N42.9927°, W5.9101° | 8/9/2020 | FL |
| ES01-010-C-00 | Spain | Léon - Puerto de la Cubilla | N42.9928°, W5.9102° | 8/9/2020 | FL |
| ES01-011-C-00 | Spain | Léon - Puerto de la Cubilla | N42.9927°, W5.9103° | 8/9/2020 | FL |
| ES01-012-C-00 | Spain | Léon - Puerto de la Cubilla | N42.9926°, W5.9107° | 8/9/2020 | FL |
| ES01-014-S-00 | Spain | Léon - Puerto de la Cubilla | N42.9927°, W5.91155° | 8/9/2020 | FL |
| ES01-015-S-00 | Spain | Léon - Puerto de la Cubilla | N42.9927°, W5.9115° | 8/9/2020 | FL |
| ES01-016-S-00 | Spain | Léon - Puerto de la Cubilla | N42.9937°, W5.9123° | 8/9/2020 | FL |
| ES01-017-C-00 | Spain | Léon - Puerto de la Cubilla | N42.9933°, W5.9110° | 8/9/2020 | FL |
| ES01-018-C-00 | Spain | Léon - Puerto de la Cubilla | N42.9934°, W5.9109° | 8/9/2020 | FL |
| ES01-019-C-00 | Spain | Léon - Puerto de la Cubilla | N42.9901°, W5.9059° | 8/9/2020 | FL |
| ES01-020-S-00 | Spain | Léon - Puerto de la Cubilla | N42.9901°, W5.9144° | 8/9/2020 | FL |
| ES01-046-S-01 | Spain | Asturias - Puerto de la Cubilla | N42.9936°, W5.9089° | 9/13/2010 | CL |
| ES01-049-S-01 | Spain | Asturias - Puerto de la Cubilla | N42.9936°, W5.9089° | 3/27/2015 | CL |
| ES01-059-S-01 | Spain | Asturias - Puerto de la Cubilla | N42.9936°, W5.9089° | 8/6/2013 | CL |
| ES01-065-S-01 | Spain | Asturias - Puerto de la Cubilla | N42.9936°, W5.9089° | 4/1/2015 | CL |
| ES01-074-S-01 | Spain | Asturias - Puerto de la Cubilla | N42.9936°, W5.9089° | 9/1/2012 | CL |
| ES01-084-S-01 | Spain | Asturias - Puerto de la Cubilla | N42.9936°, W5.9089° | 9/25/2014 | CL |
| ES01-103-S-01 | Spain | Asturias - Puerto de la Cubilla | N42.9936°, W5.9089° | 4/1/2015 | CL |
| ES01-108-S-01 | Spain | Asturias - Puerto de la Cubilla | N42.9918°, W5.9081° | 9/2/2012 | CL |
| ES02-001-S-00 | Spain | Asturias - Puerto de Pajares | N42.9923°, W5.7604° | 8/8/2020 | FL |
| ES02-002-S-00 | Spain | Asturias - Puerto de Pajares | N42.9921°, W5.7605° | 8/8/2020 | FL |
| ES02-003-S-01 | Spain | Asturias - Puerto de Pajares | N42.9924°, W5.7597° | 9/2/2012 | CL |
| ES02-004-S-00 | Spain | Asturias - Puerto de Pajares | N42.9932°, W5.7450° | 8/8/2020 | FL |
| ES02-005-C-00 | Spain | Asturias - Puerto de Pajares | N42.9930°, W5.7440° | 8/8/2020 | FL |
| ES03-001-S-00 | Spain | Asturias - Angliru | N43.2276°, W5.9367° | 8/7/2020 | FL |
| ES03-002-C-00 | Spain | Asturias - Angliru | N43.2279°, W5.9373° | 8/7/2020 | FL |
| ES03-003-C-00 | Spain | Asturias - Angliru | N43.2293°, W5.9379° | 8/7/2020 | FL |
| ES03-005-C-00 | Spain | Asturias - Angliru | N43.2298°, W5.9385° | 8/7/2020 | FL |
| ES03-006-C-00 | Spain | Asturias - Angliru | N43.2298°, W5.9385° | 8/7/2020 | FL |
| ES03-007-C-00 | Spain | Asturias - Angliru | N43.2299°, W5.9410° | 8/7/2020 | FL |
| ES03-008-C-00 | Spain | Asturias - Angliru | N43.2300°, W5.9412° | 8/7/2020 | FL |
| ES03-038-S-01 | Spain | Asturias - Angliru | N43.2298°, W5.9390° | 3/27/2015 | CL |
| ES03-009-C-00 | Spain | Asturias - Angliru | N43.2298°, W5.9423° | 8/7/2020 | FL |
| ES03-010-C-00 | Spain | Asturias - Angliru | N43.2304°, W5.9438° | 8/7/2020 | FL |
| ES03-011-C-00 | Spain | Asturias - Angliru | N43.2303°, W5.9440° | 8/7/2020 | FL |
| ES03-035-S-01 | Spain | Asturias - Angliru | N43.2298°, W5.9390° | 9/6/2012 | CL |
| ES03-012-C-00 | Spain | Asturias - Angliru | N43.2305°, W5.9443° | 8/7/2020 | FL |
| ES03-014-C-00 | Spain | Asturias - Angliru | N43.2310°, W5.9443° | 8/7/2020 | FL |
| ES03-015-C-00 | Spain | Asturias - Angliru | N43.2319°, W5.94504° | 8/7/2020 | FL |
| ES03-016-C-00 | Spain | Asturias - Angliru | N43.2319°, W5.9450° | 8/7/2020 | FL |
| ES03-037-S-01 | Spain | Asturias - Angliru | N43.2298°, W5.9390° | 2/20/2014 | CL |
| ES03-017-C-00 | Spain | Asturias - Angliru | N43.2218°, W5.9429° | 8/7/2020 | FL |
| ES03-018-C-00 | Spain | Asturias - Angliru | N43.2219°, W5.9430° | 8/7/2020 | FL |
| ES03-036-S-01 | Spain | Asturias - Angliru | N43.2298°, W5.9390° | 7/3/2014 | CL |
| ES03-019-C-00 | Spain | Asturias - Angliru | N43.2220°, W5.9431° | 8/7/2020 | FL |
| ES03-020-C-00 | Spain | Asturias - Angliru | N43.2298°, W5.9390° | 8/7/2020 | FL |
| ES03-021-C-00 | Spain | Asturias - Angliru | N43.2231°, W5.9446° | 8/7/2020 | FL |
| ES03-022-C-00 | Spain | Asturias - Angliru | N43.2233°, W5.9437° | 8/7/2020 | FL |
| ES03-023-C-00 | Spain | Asturias - Angliru | N43.2234°, W5.9435° | 8/7/2020 | FL |
| ES03-024-S-01 | Spain | Asturias - Angliru | N43.2298°, W5.9390° | 9/25/2014 | CL |
| ES03-026-S-00 | Spain | Asturias - Angliru | N43.2281°, W5.9345° | 8/7/2020 | FL |

| | | | | | |
|---|---|---|---|---|---|
| ES03-028-C-00 | Spain | Asturias - Angliru | N43.2315°, W5.9373° | 8/7/2020 | FL |
| ES03-029-C-00 | Spain | Asturias - Angliru | N43.2317°, W5.9373° | 8/7/2020 | FL |
| ES03-030-S-00 | Spain | Asturias - Angliru | N43.2316°, W5.9370° | 8/7/2020 | FL |
| ES03-031-C-00 | Spain | Asturias - Angliru | N43.2292°, W5.9385° | 9/8/2021 | FL |
| ES03-032-C-00 | Spain | Asturias - Angliru | N43.2291°, W5.9386° | 9/8/2021 | FL |
| ES03-033-C-00 | Spain | Asturias - Angliru | N43.2288°, W5.9371° | 9/8/2021 | FL |
| ES03-047-S-01 | Spain | Asturias - Angliru | N43.2298°, W5.9390° | 2/20/2014 | CL |
| ES03-048-S-01 | Spain | Asturias - Angliru | N43.2298°, W5.9390° | 2/20/2014 | CL |
| ES03-055-S-01 | Spain | Asturias - Angliru | N43.2298°, W5.9390° | 2/20/2014 | CL |
| ES03-034-S-01 | Spain | Asturias - Angliru | N43.2298°, W5.9390° | 2/20/2014 | CL |
| ES04-001-C-00 | Spain | Asturias - Lagos de Saliencia | N43.0559°, W6.1016° | 8/10/2020 | FL |
| ES04-002-C-00 | Spain | Asturias - Lagos de Saliencia | N43.0518°, W6.1036° | 8/10/2020 | FL |
| ES04-003-C-00 | Spain | Asturias - Lagos de Saliencia | N43.0517°, W6.1036° | 8/10/2020 | FL |
| ES04-004-C-00 | Spain | Asturias - Lagos de Saliencia | N43.0517°, W6.1037° | 8/10/2020 | FL |
| ES04-005-C-00 | Spain | Asturias - Lagos de Saliencia | N43.0533°, W6.1014° | 8/10/2020 | FL |
| ES04-006-C-00 | Spain | Asturias - Lagos de Saliencia | N43.0533°, W6.1014° | 8/10/2020 | FL |
| ES04-008-C-00 | Spain | Asturias - Lagos de Saliencia | N43.0519°, W6.1040° | 8/10/2020 | FL |
| ES04-009-C-00 | Spain | Asturias - Lagos de Saliencia | N43.0519°, W6.1041° | 8/10/2020 | FL |
| ES04-010-C-00 | Spain | Asturias - Lagos de Saliencia | N43.0519°, W6.1038° | 8/10/2020 | FL |
| ES04-011-C-00 | Spain | Asturias - Lagos de Saliencia | N43.0512°, W6.1052° | 8/10/2020 | FL |
| ES04-012-C-00 | Spain | Asturias - Lagos de Saliencia | N43.0512°, W6.1053° | 8/10/2020 | FL |
| ES04-013-C-00 | Spain | Asturias - Lagos de Saliencia | N43.0513°, W6.1053° | 8/10/2020 | FL |
| ES04-014-C-00 | Spain | Asturias - Lagos de Saliencia | N43.0514°, W6.1052° | 8/10/2020 | FL |
| ES04-015-C-00 | Spain | Asturias - Lagos de Saliencia | N43.0514°, W6.1051° | 8/10/2020 | FL |
| ES04-016-C-00 | Spain | Asturias - Lagos de Saliencia | N43.0516°, W6.1051° | 8/10/2020 | FL |
| ES04-017-C-00 | Spain | Asturias - Lagos de Saliencia | N43.0513°, W6.1041° | 8/10/2020 | FL |
| ES04-018-C-00 | Spain | Asturias - Lagos de Saliencia | N43.0513°, W6.1040° | 8/10/2020 | FL |
| ES04-019-C-00 | Spain | Asturias - Lagos de Saliencia | N43.0512°, W6.1040° | 8/10/2020 | FL |
| ES04-020-C-00 | Spain | Asturias - Lagos de Saliencia | N43.0514°, W6.1053° | 8/10/2020 | FL |
| ES04-021-S-00 | Spain | Asturias - Lagos de Saliencia | N43.0512°, W6.1025° | 8/10/2020 | FL |
| ES04-022-C-00 | Spain | Asturias - Lagos de Saliencia | N43.0512°, W6.1025° | 8/10/2020 | FL |
| ES04-041-S-01 | Spain | Asturias - Lagos de Saliencia | N43.0513°, W6.1042° | 8/5/2013 | CL |
| ES04-024-S-00 | Spain | Asturias - Lagos de Saliencia | N43.0511°, W6.1026° | 8/10/2020 | FL |
| ES04-025-C-00 | Spain | Asturias - Lagos de Saliencia | N43.0511°, W6.1023° | 8/10/2020 | FL |
| ES04-026-C-00 | Spain | Asturias - Lagos de Saliencia | N43.0509°, W6.1024° | 8/10/2020 | FL |
| ES04-027-C-00 | Spain | Asturias - Lagos de Saliencia | N43.0510°, W6.1023° | 8/10/2020 | FL |
| ES04-028-C-00 | Spain | Asturias - Lagos de Saliencia | N43.0510°, W6.1022° | 8/10/2020 | FL |
| ES04-029-C-00 | Spain | Asturias - Lagos de Saliencia | N43.0510°, W6.1026° | 8/10/2020 | FL |
| ES04-042-S-01 | Spain | Asturias - Lagos de Saliencia | N43.0513°, W6.1042° | 2/20/2014 | CL |
| ES04-030-C-00 | Spain | Asturias - Lagos de Saliencia | N43.0512°, W6.1038° | 8/10/2020 | FL |
| ES04-031-C-00 | Spain | Asturias - Lagos de Saliencia | N43.0511°, W6.1039° | 8/10/2020 | FL |
| ES04-032-C-00 | Spain | Asturias - Lagos de Saliencia | N43.0511°, W6.1037° | 8/10/2020 | FL |
| ES04-033-C-00 | Spain | Asturias - Lagos de Saliencia | N43.0510°, W6.1040° | 8/10/2020 | FL |
| ES04-043-S-01 | Spain | Asturias - Lagos de Saliencia | N43.0513°, W6.1042° | 3/27/2015 | CL |
| ES04-034-C-00 | Spain | Asturias - Lagos de Saliencia | N43.0510°, W6.1038° | 8/10/2020 | FL |
| ES04-035-C-00 | Spain | Asturias - Lagos de Saliencia | N43.0514°, W6.1053° | 9/7/2021 | FL |
| ES04-036-C-00 | Spain | Asturias - Lagos de Saliencia | N43.0513°, W6.1015° | 9/7/2021 | FL |
| ES04-037-C-00 | Spain | Asturias - Lagos de Saliencia | N43.0498°, W6.1026° | 9/7/2021 | FL |
| ES04-057-S-01 | Spain | Asturias - Lagos de Saliencia | N43.0513°, W6.1042° | nan | CL |
| ES04-038-C-00 | Spain | Asturias - Lagos de Saliencia | N43.0519°, W6.1087° | 9/7/2021 | FL |
| ES04-039-C-00 | Spain | Asturias - Lagos de Saliencia | N43.0519°, W6.1087° | 9/7/2021 | FL |
| ES04-045-S-01 | Spain | Asturias - Lagos de Saliencia | N43.0513°, W6.1042° | 2/20/2014 | CL |
| ES04-046-S-01 | Spain | Asturias - Lagos de Saliencia | N43.0513°, W6.1042° | 2/20/2014 | CL |
| ES04-050-S-01 | Spain | Asturias - Lagos de Saliencia | N43.0513°, W6.1042° | 2/20/2014 | CL |
| ES04-051-S-01 | Spain | Asturias - Lagos de Saliencia | N43.0513°, W6.1042° | 9/4/2012 | CL |
| ES04-052-S-01 | Spain | Asturias - Lagos de Saliencia | N43.0513°, W6.1042° | 4/20/2015 | CL |
| ES04-064-S-01 | Spain | Asturias - Lagos de Saliencia | N43.0513°, W6.1042° | 9/5/2012 | CL |
| ES04-072-S-01 | Spain | Asturias - Lagos de Saliencia | N43.0513°, W6.1042° | 8/5/2013 | CL |
| ES05-001-C-00 | Spain | Asturias - El Coto | N43.0828°, W6.2359° | 8/11/2020 | FL |
| ES05-002-S-00 | Spain | Asturias - El Coto | N43.0826°, W6.2363° | 8/11/2020 | FL |
| ES05-003-C-00 | Spain | Asturias - El Coto | N43.0833°, W6.2378° | 8/11/2020 | FL |
| ES05-004-C-00 | Spain | Asturias - El Coto | N43.0833°, W6.2378° | 8/11/2020 | FL |
| ES05-005-C-00 | Spain | Asturias - El Coto | N43.0835°, W6.2380° | 8/11/2020 | FL |
| ES05-006-C-00 | Spain | Asturias - El Coto | N43.0800°, W6.2348° | 8/11/2020 | FL |
| ES05-007-C-00 | Spain | Asturias - El Coto | N43.0800°, W6.2348° | 8/11/2020 | FL |
| ES05-009-S-00 | Spain | Asturias - El Coto | N43.0779°, W6.2342° | 8/11/2020 | FL |
| ES05-010-C-00 | Spain | Asturias - El Coto | N43.0775°, W6.2339° | 8/11/2020 | FL |

| | | | | | |
|---|---|---|---|---|---|
| ES05-011-S-00 | Spain | Asturias - El Coto | N43.0828°, W6.2365° | 8/11/2020 | FL |
| ES05-012-S-00 | Spain | Asturias - El Coto | N43.0828°, W6.2364° | 8/11/2020 | FL |
| ES05-013-S-00 | Spain | Asturias - El Coto | N43.0828°, W6.2363° | 8/11/2020 | FL |
| ES05-014-S-00 | Spain | Asturias - El Coto | N43.0830°, W6.2371° | 8/11/2020 | FL |
| ES05-016-S-00 | Spain | Asturias - El Coto | N43.0832°, W6.2377° | 8/11/2020 | FL |
| ES05-017-C-00 | Spain | Asturias - El Coto | N43.0834°, W6.2380° | 8/11/2020 | FL |
| ES05-018-C-00 | Spain | Asturias - El Coto | N43.0835°, W6.2387° | 8/11/2020 | FL |
| ES05-019-C-00 | Spain | Asturias - El Coto | N43.0799°, W6.2347° | 8/11/2020 | FL |
| ES05-020-C-00 | Spain | Asturias - El Coto | N43.0798°, W6.2346° | 8/11/2020 | FL |
| ES05-021-C-00 | Spain | Asturias - El Coto | N43.0779°, W6.2343° | 8/11/2020 | FL |
| ES05-022-C-00 | Spain | Asturias - El Coto | N43.0779°, W6.2344° | 8/11/2020 | FL |
| ES05-023-S-00 | Spain | Asturias - El Coto | N43.0780°, W6.2346° | 8/11/2020 | FL |
| ES06-001-S-01 | Spain | Asturias - Puerto de la Cubilla | N43.0007°, W5.9269° | 9/10/2014 | CL |
| ES06-002-S-01 | Spain | Asturias - Puerto de la Cubilla | N43.0007°, W5.9269° | 7/2/2014 | CL |
| ES06-003-C-00 | Spain | Asturias - Puerto de la Cubilla | N42.9993°, W5.9280° | 8/9/2020 | FL |
| ES06-004-S-00 | Spain | Asturias - Puerto de la Cubilla | N42.9995°, W5.9282° | 8/9/2020 | FL |
| ES06-005-S-00 | Spain | Asturias - Puerto de la Cubilla | N42.9995°, W5.9282° | 8/9/2020 | FL |
| ES06-006-C-00 | Spain | Asturias - Puerto de la Cubilla | N42.9998°, W5.9283° | 8/9/2020 | FL |
| ES06-007-C-00 | Spain | Asturias - Puerto de la Cubilla | N43.0001°, W5.9288° | 8/9/2020 | FL |
| ES06-008-C-00 | Spain | Asturias - Puerto de la Cubilla | N43.0003°, W5.9288° | 8/9/2020 | FL |
| ES06-008-S-01 | Spain | Asturias - Puerto de la Cubilla | N43.0007°, W5.9269° | 9/18/2014 | CL |
| ES06-009-C-00 | Spain | Asturias - Puerto de la Cubilla | N43.0004°, W5.9289° | 8/9/2020 | FL |
| ES06-010-S-01 | Spain | Asturias - Puerto de la Cubilla | N43.0007°, W5.9269° | 6/20/2014 | CL |
| ES06-011-C-00 | Spain | Asturias - Puerto de la Cubilla | N43.0026°, W5.9283° | 8/9/2020 | FL |
| ES06-012-S-00 | Spain | Asturias - Puerto de la Cubilla | N42.9995°, W5.92821° | 8/9/2020 | FL |
| ES06-015-C-00 | Spain | Asturias - Puerto de la Cubilla | N43.0008°, W5.9274° | 9/6/2021 | FL |
| ES06-016-C-00 | Spain | Asturias - Puerto de la Cubilla | N43.0007°, W5.9278° | 9/6/2021 | FL |
| ES06-017-C-00 | Spain | Asturias - Puerto de la Cubilla | N43.0008°, W5.9287° | 9/6/2021 | FL |
| ES06-018-C-00 | Spain | Asturias - Puerto de la Cubilla | N43.0009°, W5.9285° | 9/6/2021 | FL |
| ES06-020-S-01 | Spain | Asturias - Puerto de la Cubilla | N43.0007°, W5.9269° | nan | CL |
| ES06-021-C-00 | Spain | Asturias - Puerto de la Cubilla | N43.0017°, W5.9307° | 9/6/2021 | FL |
| ES06-022-C-00 | Spain | Asturias - Puerto de la Cubilla | N43.0014°, W5.9311° | 9/6/2021 | FL |
| ES06-023-C-00 | Spain | Asturias - Puerto de la Cubilla | N43.0013°, W5.9312° | 9/6/2021 | FL |
| ES06-026-C-00 | Spain | Asturias - Puerto de la Cubilla | N42.9989°, W5.9301° | 9/6/2021 | FL |
| ES06-041-S-01 | Spain | Asturias - Puerto de la Cubilla | N43.0007°, W5.9269° | 9/19/2014 | CL |
| ES06-045-S-01 | Spain | Asturias - Puerto de la Cubilla | N43.0007°, W5.9269° | 7/3/2014 | CL |
| ES06-050-S-01 | Spain | Asturias - Puerto de la Cubilla | N43.0007°, W5.9269° | 9/25/2014 | CL |
| ES08-001-S-01 | Spain | Asturias - Naranjo de Bulnes | N43.2071°, W4.8149° | 7/17/2014 | CL |
| ES08-002-S-01 | Spain | Asturias - Naranjo de Bulnes | N43.2071°, W4.8149° | 7/2/2014 | CL |
| ES08-004-S-01 | Spain | Asturias - Naranjo de Bulnes | N43.2071°, W4.8149° | 9/25/2014 | CL |
| ES08-005-S-01 | Spain | Asturias - Naranjo de Bulnes | N43.2071°, W4.8149° | 7/3/2014 | CL |
| ES08-006-S-01 | Spain | Asturias - Naranjo de Bulnes | N43.2071°, W4.8149° | 9/18/2014 | CL |
| ES08-007-S-01 | Spain | Asturias - Naranjo de Bulnes | N43.2071°, W4.8149° | 8/10/2009 | CL |
| ES08-008-S-01 | Spain | Asturias - Naranjo de Bulnes | N43.2071°, W4.8149° | 7/1/2014 | CL |
| ES08-009-S-01 | Spain | Asturias - Naranjo de Bulnes | N43.2071°, W4.8149° | 8/10/2009 | CL |
| ES08-010-S-01 | Spain | Asturias - Naranjo de Bulnes | N43.2071°, W4.8149° | 8/10/2009 | CL |
| ES09-001-S-01 | Spain | Cantabria - Fuente Dé | N43.1657°, W4.8126° | 8/4/2013 | CL |
| ES09-003-S-01 | Spain | Cantabria - Fuente Dé | N43.1657°, W4.8126° | 7/20/2015 | CL |
| ES09-004-S-01 | Spain | Cantabria - Fuente Dé | N43.1490°, W4.8075° | 8/4/2013 | CL |
| ES10-001-C-00 | Spain | Asturias - Cueva Huerta | N43.1236°, W6.0596° | 8/11/2020 | FL |
| ES10-002-C-00 | Spain | Asturias - Cueva Huerta | N43.1237°, W6.0586° | 8/11/2020 | FL |
| ES10-003-C-00 | Spain | Asturias - Cueva Huerta | N43.1237°, W6.0586° | 8/11/2020 | FL |
| ES10-004-C-00 | Spain | Asturias - Cueva Huerta | N43.1237°, W6.0586° | 8/11/2020 | FL |
| ES10-005-S-01 | Spain | Asturias - Cueva Huerta | N43.1237°, W6.0586° | 8/11/2020 | FL |
| ES15-001-S-00 | Spain | Asturias - Entrepeñas | N43.1426°, W5.5792° | 8/8/2020 | FL |
| ES17-001-C-00 | Spain | Léon - Puebla de Lillo | N43.0098°, W5.2775° | 8/12/2020 | FL |
| ES17-002-C-00 | Spain | Léon - Puebla de Lillo | N43.0120°, W5.2788° | 8/12/2020 | FL |
| ES17-004-C-00 | Spain | Léon - Puebla de Lillo | N43.0126°, W5.2792° | 8/12/2020 | FL |
| ES17-005-C-00 | Spain | Léon - Puebla de Lillo | N43.0126°, W5.2793° | 8/12/2020 | FL |
| ES17-006-C-00 | Spain | Léon - Puebla de Lillo | N43.0127°, W5.2793° | 8/12/2020 | FL |
| ES17-007-C-00 | Spain | Léon - Puebla de Lillo | N43.0127°, W5.2794° | 8/12/2020 | FL |
| ES17-009-C-00 | Spain | Léon - Puebla de Lillo | N43.0095°, W5.2780° | 8/12/2020 | FL |
| ES17-010-C-00 | Spain | Léon - Puebla de Lillo | N43.0126°, W5.2794° | 8/12/2020 | FL |
| ES17-011-C-00 | Spain | Léon - Puebla de Lillo | N43.0132°, W5.2794° | 8/12/2020 | FL |
| ES17-012-C-00 | Spain | Léon - Puebla de Lillo | N43.0094°, W5.2763° | 8/12/2020 | FL |
| ES17-014-C-00 | Spain | Léon - Puebla de Lillo | N43.0156°, W5.2836° | 8/12/2020 | FL |
| ES17-015-C-00 | Spain | Léon - Puebla de Lillo | N43.0155°, W5.2835° | 8/12/2020 | FL |

| | | | | | |
|---|---|---|---|---|---|
| ES17-016-C-00 | Spain | Léon - Puebla de Lillo | N43.0155°, W5.2835° | 8/12/2020 | FL |
| ES17-017-C-00 | Spain | Léon - Puebla de Lillo | N43.0155°, W5.2834° | 8/12/2020 | FL |
| ES17-018-C-00 | Spain | Léon - Puebla de Lillo | N43.0155°, W5.2834° | 8/12/2020 | FL |
| ES23-001-S-00 | Spain | Asturias - Puerto de Leitariegos | N43.0016°, W6.4242° | 8/15/2020 | CAB |
| ES23-003-S-00 | Spain | Asturias - Puerto de Leitariegos | N43.0016°, W6.4242° | 8/15/2020 | CAB |
| ES23-005-S-00 | Spain | Asturias - Puerto de Leitariegos | N43.0016°, W6.4242° | 8/15/2020 | CAB |
| ES23-006-S-00 | Spain | Asturias - Puerto de Leitariegos | N43.0016°, W6.4242° | 8/15/2020 | CAB |
| ES23-007-S-00 | Spain | Asturias - Puerto de Leitariegos | N43.0016°, W6.4242° | 8/15/2020 | CAB |
| ES24-001-S-01 | Spain | Cantabria - Brañavieja | N43.0397°, W4.3631° | nan | CAB |
| ES24-002-S-01 | Spain | Cantabria - Brañavieja | N43.0397°, W4.3631° | nan | CAB |
| ES24-004-S-01 | Spain | Cantabria - Brañavieja | N43.0397°, W4.3631° | nan | CAB |
| ES24-005-S-01 | Spain | Cantabria - Brañavieja | N43.0397°, W4.3631° | nan | CAB |
| ES25-001-S-01 | Spain | Burgos - Portillo de la Sia | N43.1532°, W3.5699° | nan | CAB |
| ES25-002-S-01 | Spain | Burgos - Portillo de la Sia | N43.1532°, W3.5699° | nan | CAB |
| ES25-003-S-01 | Spain | Burgos - Portillo de la Sia | N43.1532°, W3.5699° | nan | CAB |
| ES25-004-S-01 | Spain | Burgos - Portillo de la Sia | N43.1532°, W3.5699° | nan | CAB |
| ES25-005-S-01 | Spain | Burgos - Portillo de la Sia | N43.1532°, W3.5699° | nan | CAB |
| ES25-006-S-01 | Spain | Burgos - Portillo de la Sia | N43.1532°, W3.5699° | nan | CAB |
| ES25-007-S-01 | Spain | Burgos - Portillo de la Sia | N43.1532°, W3.5699° | nan | CAB |
| ES25-008-S-01 | Spain | Burgos - Portillo de la Sia | N43.1532°, W3.5699° | nan | CAB |
| ES25-009-S-01 | Spain | Burgos - Portillo de la Sia | N43.1532°, W3.5699° | nan | CAB |
| FR01-001-S-01 | France | Savoy - Grand Galibier | N45.0550°, E6.44995° | 9/15/2018 | CL |
| FR01-002-S-01 | France | Savoy - Grand Galibier | N45.0550°, E6.44995° | 9/15/2018 | CL |
| FR01-003-S-01 | France | Savoy - Grand Galibier | N45.0559°, E6.44644° | 9/15/2018 | CL |
| FR01-004-S-01 | France | Savoy - Grand Galibier | N45.0584°, E6.44095° | 9/15/2018 | CL |
| FR01-005-S-01 | France | Savoy - Grand Galibier | N45.0607°, E6.43910° | 9/15/2018 | CL |
| FR01-006-S-01 | France | Savoy - Grand Galibier | N45.0636°, E6.43435° | 9/15/2018 | CL |
| FR01-007-S-01 | France | Savoy - Grand Galibier | N45.0634°, E6.43489° | 9/15/2018 | CL |
| FR01-008-S-01 | France | Savoy - Grand Galibier | N45.0639°, E6.43389° | 9/15/2018 | CL |
| FR01-009-S-01 | France | Hautes-Alpes - Col du Galibier | N45.0594°, E6.40532° | 9/23/2012 | CL |
| FR01-013-S-01 | France | Hautes-Alpes - Col du Galibier | N45.0594°, E6.40532° | 9/23/2012 | CL |
| FR01-015-S-01 | France | Hautes-Alpes - Col du Galibier | N45.0594°, E6.40532° | nan | CL |
| FR01-016-S-01 | France | Hautes-Alpes - Col du Galibier | N45.0594°, E6.40532° | nan | CL |
| FR01-018-S-01 | France | Hautes-Alpes - Col du Galibier | N45.0594°, E6.40532° | nan | CL |
| FR01-020-S-01 | France | Hautes-Alpes - Col du Galibier | N45.0594°, E6.40532° | nan | CL |
| FR01-028-S-01 | France | Hautes-Alpes - Col du Galibier | N45.0594°, E6.40532° | 9/23/2012 | CL |
| FR01-035-S-01 | France | Hautes-Alpes - Col du Galibier | N45.0594°, E6.40532° | nan | CL |
| FR01-036-S-01 | France | Hautes-Alpes - Col du Galibier | N45.0594°, E6.40532° | nan | CL |
| FR01-038-S-01 | France | Hautes-Alpes - Col du Galibier | N45.0594°, E6.40532° | nan | CL |
| FR01-039-S-01 | France | Hautes-Alpes - Col du Galibier | N45.0594°, E6.40532° | nan | CL |
| FR01-040-S-01 | France | Hautes-Alpes - Col du Galibier | N45.0594°, E6.40532° | nan | CL |
| FR01-046-S-01 | France | Hautes-Alpes - Col du Galibier | N45.0594°, E6.40532° | nan | CL |
| FR01-047-S-01 | France | Hautes-Alpes - Col du Galibier | N45.0594°, E6.40532° | nan | CL |
| FR01-055-S-01 | France | Hautes-Alpes - Col du Galibier | N45.0594°, E6.40532° | 9/23/2012 | CL |
| FR01-062-S-01 | France | Hautes-Alpes - Col du Galibier | N45.0594°, E6.40532° | 9/23/2012 | CL |
| FR01-070-S-01 | France | Hautes-Alpes - Col du Galibier | N45.0594°, E6.40532° | nan | CL |
| FR01-072-S-01 | France | Hautes-Alpes - Col du Galibier | N45.0601°, E6.40347° | 9/23/2012 | CL |
| FR01-073-S-01 | France | Hautes-Alpes - Col du Galibier | N45.0602°, E6.40344° | 9/23/2012 | CL |
| FR01-075-S-01 | France | Hautes-Alpes - Col du Galibier | N45.0602°, E6.40344° | nan | CL |
| FR01-079-S-01 | France | Hautes-Alpes - Col du Galibier | N45.0602°, E6.40344° | 9/23/2012 | CL |
| FR01-081-S-01 | France | Hautes-Alpes - Col du Galibier | N45.0602°, E6.40344° | 9/23/2012 | CL |
| FR01-083-S-01 | France | Hautes-Alpes - Col du Galibier | N45.0602°, E6.40344° | 9/23/2012 | CL |
| FR01-086-S-01 | France | Hautes-Alpes - Col du Galibier | N45.0602°, E6.40344° | 9/23/2012 | CL |
| FR01-090-S-01 | France | Hautes-Alpes - Col du Galibier | N45.0604°, E6.40303° | 9/23/2012 | CL |
| FR01-098-S-01 | France | Hautes-Alpes - Col du Galibier | N45.0604°, E6.40303° | 9/23/2012 | CL |
| FR01-100-S-01 | France | Hautes-Alpes - Col du Galibier | N45.0604°, E6.40285° | 9/23/2012 | CL |
| FR01-104-S-01 | France | Hautes-Alpes - Col du Galibier | N45.0604°, E6.40285° | nan | CL |
| FR01-105-S-01 | France | Hautes-Alpes - Col du Galibier | N45.0604°, E6.40285° | 9/23/2012 | CL |
| FR01-111-S-01 | France | Hautes-Alpes - Col du Galibier | N45.0604°, E6.40285° | nan | CL |
| FR01-115-S-01 | France | Hautes-Alpes - Col du Galibier | N45.0607°, E6.40171° | 9/23/2012 | CL |
| FR01-116-S-01 | France | Hautes-Alpes - Col du Galibier | N45.0605°, E6.40154° | 9/22/2012 | CL |
| FR01-118-S-01 | France | Hautes-Alpes - Col du Galibier | N45.0606°, E6.40130° | 9/22/2012 | CL |
| FR01-119-S-01 | France | Hautes-Alpes - Col du Galibier | N45.0605°, E6.40128° | nan | CL |
| FR01-120-S-01 | France | Hautes-Alpes - Col du Galibier | N45.0605°, E6.40112° | 9/22/2012 | CL |
| FR01-122-S-01 | France | Hautes-Alpes - Col du Galibier | N45.0607°, E6.40104° | 9/22/2012 | CL |
| FR01-124-S-01 | France | Hautes-Alpes - Col du Galibier | N45.0609°, E6.40074° | 9/22/2012 | CL |
| FR01-130-S-01 | France | Hautes-Alpes - Col du Galibier | N45.0609°, E6.40222° | nan | CL |

| | | | | | |
|---|---|---|---|---|---|
| FR01-131-S-01 | France | Hautes-Alpes - Col du Galibier | N45.0609°, E6.40222° | 9/22/2012 | CL |
| FR01-133-S-01 | France | Hautes-Alpes - Col du Galibier | N45.0609°, E6.40222° | 9/22/2012 | CL |
| FR01-135-S-01 | France | Hautes-Alpes - Col du Galibier | N45.0609°, E6.40222° | nan | CL |
| FR01-137-S-01 | France | Hautes-Alpes - Col du Galibier | N45.0609°, E6.40222° | 9/22/2012 | CL |
| FR01-139-S-01 | France | Hautes-Alpes - Col du Galibier | N45.0609°, E6.40222° | 9/22/2012 | CL |
| FR01-400-S-01 | France | Hautes-Alpes - Col du Galibier | N45.0606°, E6.40083° | 9/22/2012 | CL |
| FR01-404-S-01 | France | Hautes-Alpes - Col du Galibier | N45.0611°, E6.40030° | 10/11/2014 | CL |
| FR01-407-S-01 | France | Hautes-Alpes - Col du Galibier | N45.0616°, E6.40013° | 9/22/2012 | CL |
| FR01-409-S-01 | France | Hautes-Alpes - Col du Galibier | N45.0619°, E6.39974° | 9/22/2012 | CL |
| FR02-015-S-01 | France | Hautes-Alpes - Col du Granon | N44.9663°, E6.58244° | 9/19/2014 | CL |
| FR02-016-S-01 | France | Hautes-Alpes - Col du Granon | N44.9660°, E6.58242° | 9/19/2012 | CL |
| FR02-018-S-01 | France | Hautes-Alpes - Col du Granon | N44.9663°, E6.58244° | 8/27/2014 | CL |
| FR02-026-S-01 | France | Hautes-Alpes - Col du Granon | N44.9663°, E6.58244° | 2/20/2014 | CL |
| FR02-029-S-01 | France | Hautes-Alpes - Col du Granon | N44.9663°, E6.58244° | 2/20/2014 | CL |
| FR02-033-S-00 | France | Hautes-Alpes - Col du Granon | N44.9663°, E6.58244° | 8/24/2011 | CL |
| FR02-037-S-01 | France | Hautes-Alpes - Col du Granon | N44.9663°, E6.58244° | 8/27/2014 | CL |
| FR02-049-S-01 | France | Hautes-Alpes - Col du Granon | N44.9663°, E6.58244° | 9/19/2014 | CL |
| FR02-071-S-01 | France | Hautes-Alpes - Col du Granon | N44.9663°, E6.58244° | 5/15/2017 | CL |
| FR02-079-S-01 | France | Hautes-Alpes - Col du Granon | N44.9663°, E6.58244° | 9/25/2014 | CL |
| FR02-099-S-01 | France | Hautes-Alpes - Col du Granon | N44.9660°, E6.58242° | 7/20/2012 | CL |
| FR03-165-S-01 | France | Savoy - Pic Blanc du Galibier | N45.0639°, E6.38459° | 8/14/2012 | CL |
| FR03-175-S-01 | France | Savoy - Pic Blanc du Galibier | N45.0639°, E6.38459° | 9/17/2012 | CL |
| FR03-176-S-01 | France | Savoy - Pic Blanc du Galibier | N45.0639°, E6.38459° | 9/17/2012 | CL |
| FR03-178-S-01 | France | Savoy - Pic Blanc du Galibier | N45.0639°, E6.38459° | 9/17/2012 | CL |
| FR03-180-S-01 | France | Savoy - Pic Blanc du Galibier | N45.0639°, E6.38459° | 9/17/2012 | CL |
| FR03-181-S-01 | France | Savoy - Pic Blanc du Galibier | N45.0641°, E6.38475° | 9/17/2012 | CL |
| FR03-182-S-01 | France | Savoy - Pic Blanc du Galibier | N45.0639°, E6.38507° | 9/17/2012 | CL |
| FR03-307-S-01 | France | Savoy - Pic Blanc du Galibier | N45.0642°, E6.38467° | 9/17/2012 | CL |
| FR03-308-S-01 | France | Savoy - Pic Blanc du Galibier | N45.0643°, E6.38463° | 9/22/2013 | CL |
| FR03-309-S-01 | France | Savoy - Pic Blanc du Galibier | N45.0643°, E6.38463° | 9/22/2013 | CL |
| FR03-311-S-01 | France | Savoy - Pic Blanc du Galibier | N45.0643°, E6.38463° | 9/17/2012 | CL |
| FR03-312-S-01 | France | Savoy - Pic Blanc du Galibier | N45.0643°, E6.38463° | 9/17/2012 | CL |
| FR03-313-S-01 | France | Savoy - Pic Blanc du Galibier | N45.0643°, E6.38463° | 9/17/2012 | CL |
| FR03-315-S-01 | France | Savoy - Pic Blanc du Galibier | N45.0642°, E6.38438° | 9/22/2013 | CL |
| FR03-316-S-01 | France | Savoy - Pic Blanc du Galibier | N45.0642°, E6.38438° | 9/17/2012 | CL |
| FR03-317-S-01 | France | Savoy - Pic Blanc du Galibier | N45.0642°, E6.38438° | 9/17/2012 | CL |
| FR03-318-S-01 | France | Savoy - Pic Blanc du Galibier | N45.0642°, E6.38438° | 9/17/2012 | CL |
| FR03-319-S-01 | France | Savoy - Pic Blanc du Galibier | N45.0642°, E6.38438° | 8/14/2012 | CL |
| FR03-320-S-01 | France | Savoy - Pic Blanc du Galibier | N45.0642°, E6.38438° | 9/17/2012 | CL |
| FR03-323-S-01 | France | Savoy - Pic Blanc du Galibier | N45.0641°, E6.38408° | 8/14/2012 | CL |
| FR03-326-S-01 | France | Savoy - Pic Blanc du Galibier | N45.0640°, E6.38442° | 9/17/2012 | CL |
| FR03-327-S-01 | France | Savoy - Pic Blanc du Galibier | N45.0640°, E6.38442° | 8/14/2012 | CL |
| FR03-328-S-01 | France | Savoy - Pic Blanc du Galibier | N45.0640°, E6.38439° | 9/22/2013 | CL |
| FR04-253-S-01 | France | Savoy - Col du Galibier | N45.0618°, E6.39436° | 8/15/2012 | CL |
| FR04-254-S-01 | France | Savoy - Col du Galibier | N45.0618°, E6.39436° | 8/15/2012 | CL |
| FR04-256-S-01 | France | Savoy - Col du Galibier | N45.0618°, E6.39436° | 9/18/2012 | CL |
| FR04-258-S-01 | France | Savoy - Col du Galibier | N45.0618°, E6.39436° | 9/18/2012 | CL |
| FR04-259-S-01 | France | Savoy - Col du Galibier | N45.0618°, E6.39434° | 9/18/2012 | CL |
| FR04-260-S-01 | France | Savoy - Col du Galibier | N45.0618°, E6.39434° | 9/18/2012 | CL |
| FR04-261-S-01 | France | Savoy - Col du Galibier | N45.0623°, E6.39290° | 9/21/2013 | CL |
| FR04-263-S-01 | France | Savoy - Col du Galibier | N45.0629°, E6.39099° | 9/18/2012 | CL |
| FR04-264-S-01 | France | Savoy - Col du Galibier | N45.0630°, E6.39098° | 9/18/2012 | CL |
| FR04-265-S-01 | France | Savoy - Col du Galibier | N45.0630°, E6.39098° | 9/18/2012 | CL |
| FR04-267-S-01 | France | Savoy - Col du Galibier | N45.0630°, E6.39098° | 9/18/2012 | CL |
| FR04-272-S-01 | France | Savoy - Col du Galibier | N45.0630°, E6.39098° | 9/18/2012 | CL |
| FR04-274-S-01 | France | Savoy - Col du Galibier | N45.0641°, E6.38916° | 9/21/2013 | CL |
| FR04-302-S-01 | France | Savoy - Col du Galibier | N45.0641°, E6.38897° | 8/15/2012 | CL |
| FR04-303-S-01 | France | Savoy - Col du Galibier | N45.0641°, E6.38894° | 8/15/2012 | CL |
| FR04-304-S-01 | France | Savoy - Col du Galibier | N45.0641°, E6.38894° | 9/21/2013 | CL |
| FR04-305-S-01 | France | Savoy - Col du Galibier | N45.0641°, E6.38890° | 9/21/2013 | CL |
| FR04-350-S-01 | France | Savoy - Col du Galibier | N45.0641°, E6.38890° | 9/18/2012 | CL |
| FR04-351-S-01 | France | Savoy - Col du Galibier | N45.0639°, E6.38897° | 9/18/2012 | CL |
| FR06-004-S-01 | France | Savoy - Vallon du Fond | N45.0767°, E6.40833° | 8/13/2012 | CL |
| FR06-009-S-01 | France | Savoy - Vallon du Fond | N45.0763°, E6.39617° | 9/19/2013 | CL |
| FR06-010-S-01 | France | Savoy - Vallon du Fond | N45.0751°, E6.39122° | 8/13/2012 | CL |
| FR06-011-S-01 | France | Savoy - Vallon du Fond | N45.0751°, E6.39122° | 9/19/2013 | CL |
| FR06-012-S-01 | France | Savoy - Vallon du Fond | N45.0751°, E6.39124° | 9/19/2013 | CL |

| | | | | | |
|---|---|---|---|---|---|
| FR06-013-S-01 | France | Savoy - Vallon du Fond | N45.0751°, E6.39130° | 9/19/2013 | CL |
| FR06-014-S-01 | France | Savoy - Vallon du Fond | N45.0750°, E6.39126° | 9/19/2013 | CL |
| FR06-016-S-01 | France | Savoy - Vallon du Fond | N45.0752°, E6.38694° | 9/19/2013 | CL |
| FR06-029-S-01 | France | Savoy - Vallon du Fond | N45.0730°, E6.38124° | nan | CL |
| FR06-031-S-01 | France | Savoy - Vallon du Fond | N45.0730°, E6.38116° | nan | CL |
| FR06-037-S-01 | France | Savoy - Vallon du Fond | N45.0755°, E6.40015° | 9/19/2013 | CL |
| FR07-002-S-01 | France | Hautes-Alpes - Col du Granon | N44.9660°, E6.58242° | 7/20/2012 | CL |
| FR07-003-S-01 | France | Hautes-Alpes - Col du Granon | N44.9660°, E6.58242° | 7/20/2012 | CL |
| FR07-007-S-01 | France | Hautes-Alpes - Col du Granon | N44.9660°, E6.58242° | 7/20/2012 | CL |
| FR07-008-S-01 | France | Hautes-Alpes - Col du Granon | N44.9660°, E6.58242° | 7/20/2012 | CL |
| FR07-010-S-01 | France | Hautes-Alpes - Col du Granon | N44.9661°, E6.58259° | 7/20/2012 | CL |
| FR07-012-S-01 | France | Hautes-Alpes - Col du Granon | N44.9661°, E6.58259° | 7/20/2012 | CL |
| FR07-027-S-01 | France | Hautes-Alpes - Col du Granon | N44.9661°, E6.58259° | 8/10/2012 | CL |
| FR07-030-S-01 | France | Hautes-Alpes - Col du Granon | N44.9662°, E6.58246° | 8/10/2012 | CL |
| FR07-032-S-01 | France | Hautes-Alpes - Col du Granon | N44.9663°, E6.58251° | 8/10/2012 | CL |
| FR07-038-S-01 | France | Hautes-Alpes - Col du Granon | N44.9663°, E6.58249° | 8/10/2012 | CL |
| FR07-046-S-01 | France | Hautes-Alpes - Col du Granon | N44.9664°, E6.58243° | 8/10/2012 | CL |
| FR07-047-S-01 | France | Hautes-Alpes - Col du Granon | N44.9664°, E6.58243° | 8/10/2012 | CL |
| FR07-078-S-01 | France | Hautes-Alpes - Col du Granon | N44.9663°, E6.58249° | 8/10/2012 | CL |
| FR07-081-S-01 | France | Hautes-Alpes - Col du Granon | N44.9664°, E6.58243° | 8/10/2012 | CL |
| FR07-083-S-01 | France | Hautes-Alpes - Col du Granon | N44.9664°, E6.58243° | 8/10/2012 | CL |
| FR07-085-S-01 | France | Hautes-Alpes - Col du Granon | N44.9664°, E6.58243° | 8/10/2012 | CL |
| FR07-086-S-01 | France | Hautes-Alpes - Col du Granon | N44.9664°, E6.58243° | 8/10/2012 | CL |
| FR07-089-S-01 | France | Hautes-Alpes - Col du Granon | N44.9664°, E6.58243° | 8/10/2012 | CL |
| FR07-091-S-01 | France | Hautes-Alpes - Col du Granon | N44.9664°, E6.58243° | 7/20/2012 | CL |
| FR07-092-S-01 | France | Hautes-Alpes - Col du Granon | N44.9664°, E6.58243° | 8/10/2012 | CL |
| FR07-093-S-01 | France | Hautes-Alpes - Col du Granon | N44.9664°, E6.58243° | 7/20/2012 | CL |
| FR07-094-S-01 | France | Hautes-Alpes - Col du Granon | N44.9664°, E6.58243° | 8/10/2012 | CL |
| FR07-095-S-01 | France | Hautes-Alpes - Col du Granon | N44.9664°, E6.58243° | 7/31/2013 | CL |
| FR07-097-S-01 | France | Hautes-Alpes - Col du Granon | N44.9664°, E6.58243° | 7/20/2012 | CL |
| FR07-099-S-01 | France | Hautes-Alpes - Col du Granon | N44.9664°, E6.58243° | 7/20/2012 | CL |
| FR07-101-S-01 | France | Hautes-Alpes - Col du Granon | N44.9664°, E6.58243° | 7/20/2012 | CL |
| FR07-102-S-01 | France | Hautes-Alpes - Col du Granon | N44.9693°, E6.58112° | 7/20/2012 | CL |
| FR07-103-S-01 | France | Hautes-Alpes - Col du Granon | N44.9693°, E6.58112° | 7/20/2012 | CL |
| FR07-104-S-01 | France | Hautes-Alpes - Col du Granon | N44.9689°, E6.58104° | 7/20/2012 | CL |
| FR07-105-S-01 | France | Hautes-Alpes - Col du Granon | N44.9692°, E6.58104° | 7/20/2012 | CL |
| FR07-106-S-01 | France | Hautes-Alpes - Col du Granon | N44.9692°, E6.58104° | 7/20/2012 | CL |
| FR08-001-S-01 | France | Savoy - Col de la Croix de Fer | N45.2239°, E6.19116° | 9/20/2012 | CL |
| FR08-002-S-01 | France | Savoy - Col de la Croix de Fer | N45.2174°, E6.18842° | 9/20/2012 | CL |
| FR08-003-S-01 | France | Savoy - Col de la Croix de Fer | N45.2174°, E6.18842° | 8/16/2012 | CL |
| FR08-004-S-01 | France | Savoy - Col de la Croix de Fer | N45.2174°, E6.18842° | 8/16/2012 | CL |
| FR08-005-S-01 | France | Savoy - Col de la Croix de Fer | N45.2174°, E6.18842° | 8/16/2012 | CL |
| FR08-006-S-01 | France | Savoy - Col de la Croix de Fer | N45.2174°, E6.18842° | 9/20/2012 | CL |
| FR08-008-S-01 | France | Savoy - Col de la Croix de Fer | N45.2174°, E6.18842° | 9/20/2012 | CL |
| FR08-010-S-01 | France | Savoy - Col de la Croix de Fer | N45.2174°, E6.18842° | 8/16/2012 | CL |
| FR08-011-S-01 | France | Savoy - Col de la Croix de Fer | N45.2137°, E6.18544° | 9/20/2012 | CL |
| FR08-012-S-01 | France | Savoy - Col de la Croix de Fer | N45.2137°, E6.18544° | 9/20/2012 | CL |
| FR08-016-S-01 | France | Savoy - Col de la Croix de Fer | N45.2137°, E6.18544° | 9/20/2012 | CL |
| FR08-020-S-01 | France | Savoy - Col de la Croix de Fer | N45.2137°, E6.18544° | 9/20/2012 | CL |
| FR08-022-S-01 | France | Savoy - Col de la Croix de Fer | N45.2147°, E6.18452° | 9/20/2012 | CL |
| FR08-024-S-01 | France | Savoy - Col de la Croix de Fer | N45.2147°, E6.18452° | 9/20/2012 | CL |
| FR08-027-S-01 | France | Savoy - Col de la Croix de Fer | N45.2143°, E6.18429° | 9/20/2012 | CL |
| FR08-030-S-01 | France | Savoy - Col de la Croix de Fer | N45.2141°, E6.18404° | 9/20/2012 | CL |
| FR08-032-S-01 | France | Savoy - Col de la Croix de Fer | N45.2141°, E6.18404° | 9/20/2012 | CL |
| FR08-036-S-01 | France | Savoy - Col de la Croix de Fer | N45.2111°, E6.18240° | 9/20/2012 | CL |
| FR08-037-S-01 | France | Savoy - Col de la Croix de Fer | N45.2111°, E6.18240° | 9/20/2012 | CL |
| FR08-038-S-01 | France | Savoy - Col de la Croix de Fer | N45.2111°, E6.18240° | 8/16/2012 | CL |
| FR08-039-S-01 | France | Savoy - Col de la Croix de Fer | N45.2111°, E6.18240° | 9/20/2012 | CL |
| FR08-040-S-01 | France | Savoy - Col de la Croix de Fer | N45.2111°, E6.18208° | 9/20/2012 | CL |
| FR08-041-S-01 | France | Savoy - Col de la Croix de Fer | N45.2111°, E6.18208° | 9/21/2012 | CL |
| FR08-042-S-01 | France | Savoy - Col de la Croix de Fer | N45.2111°, E6.18208° | 9/21/2012 | CL |
| FR08-044-S-01 | France | Savoy - Col de la Croix de Fer | N45.1992°, E6.17089° | 9/21/2012 | CL |
| FR08-045-S-01 | France | Savoy - Col de la Croix de Fer | N45.1990°, E6.17082° | 9/21/2012 | CL |
| FR08-047-S-01 | France | Savoy - Col de la Croix de Fer | N45.1980°, E6.16993° | 9/21/2012 | CL |
| FR08-049-S-01 | France | Savoy - Col de la Croix de Fer | N45.1980°, E6.16988° | 9/21/2012 | CL |
| FR08-050-S-01 | France | Savoy - Col de la Croix de Fer | N45.1980°, E6.16988° | 9/21/2012 | CL |
| FR08-051-S-01 | France | Savoy - Col de la Croix de Fer | N45.1958°, E6.17311° | 9/21/2012 | CL |

| | | | | | |
|---|---|---|---|---|---|
| FR08-052-S-01 | France | Savoy - Col de la Croix de Fer | N45.1958°, E6.17311° | 9/21/2012 | CL |
| FR08-054-S-01 | France | Savoy - Col de la Croix de Fer | N45.1958°, E6.17311° | 9/21/2012 | CL |
| FR08-055-S-01 | France | Savoy - Col de la Croix de Fer | N45.1957°, E6.17295° | 9/24/2013 | CL |
| FR08-060-S-01 | France | Savoy - Col de la Croix de Fer | N45.1957°, E6.17295° | 9/21/2012 | CL |
| FR08-061-S-01 | France | Savoy - Col de la Croix de Fer | N45.1714°, E6.16838° | 9/24/2013 | CL |
| FR10-001-S-01 | France | Hautes-Alpes - Briançon | N44.9004°, E6.6430° | 3/7/2010 | CL |
| FR10-003-S-01 | France | Hautes-Alpes - Briançon | N44.9004°, E6.6430° | 3/7/2010 | CL |
| FR10-004-S-01 | France | Hautes-Alpes - Briançon | N44.9004°, E6.6430° | 3/7/2010 | CL |
| FR10-005-S-01 | France | Hautes-Alpes - Briançon | N44.9004°, E6.6430° | 3/7/2010 | CL |
| FR15-325-S-01 | France | Savoy - La Valloirette | N45.0882°, E6.43233° | 8/13/2012 | CL |
| FR15-326-S-01 | France | Savoy - La Valloirette | N45.0882°, E6.43233° | 8/13/2012 | CL |
| FR15-327-S-01 | France | Savoy - La Valloirette | N45.0882°, E6.43233° | 8/13/2012 | CL |
| FR15-328-S-01 | France | Savoy - La Valloirette | N45.0882°, E6.43233° | 9/19/2013 | CL |
| FR15-329-S-01 | France | Savoy - La Valloirette | N45.0882°, E6.43233° | 9/19/2013 | CL |
| FR15-330-S-01 | France | Savoy - La Valloirette | N45.0811°, E6.42592° | 8/13/2012 | CL |
| FR15-331-S-01 | France | Savoy - La Valloirette | N45.0811°, E6.42584° | 8/13/2012 | CL |
| FR15-332-S-01 | France | Savoy - La Valloirette | N45.0814°, E6.42587° | 8/13/2012 | CL |
| FR16-001-C-00 | France | Hautes-Alpes - Col de la Trancoulette | N44.8917°, E6.55038° | 7/11/2021 | CL |
| FR16-002-C-00 | France | Hautes-Alpes - Col de la Trancoulette | N44.8917°, E6.55038° | 7/11/2021 | CL |
| FR16-004-C-00 | France | Hautes-Alpes - Col de la Trancoulette | N44.8917°, E6.55038° | 7/11/2021 | CL |
| FR17-001-C-00 | France | Hautes-Alpes - Col de la Pisse | N44.9038°, E6.52796° | 7/12/2021 | CL |
| FR17-002-C-00 | France | Hautes-Alpes - Col de la Pisse | N44.9038°, E6.52796° | 7/12/2021 | CL |
| FR17-003-C-00 | France | Hautes-Alpes - Col de la Pisse | N44.9038°, E6.52796° | 7/12/2021 | CL |
| FR17-004-C-00 | France | Hautes-Alpes - Col de la Pisse | N44.9038°, E6.52796° | 7/12/2021 | CL |
| FR18-005-C-00 | France | Hautes-Alpes - Col des Marseilles | N44.8322°, E6.81208° | 7/14/2021 | CL |
| FR19-001-C-00 | France | Hautes-Alpes - Cime de la Condamine | N44.8933°, E6.51945° | 7/20/2021 | CL |
| FR19-002-C-00 | France | Hautes-Alpes - Cime de la Condamine | N44.8933°, E6.51945° | 7/20/2021 | CL |
| FR19-003-C-00 | France | Hautes-Alpes - Cime de la Condamine | N44.8933°, E6.51945° | 7/20/2021 | CL |
| FR20-001-C-00 | France | Hautes-Alpes - Les Combes | N44.8944°, E6.56460° | 7/21/2021 | CL |
| FR20-004-C-00 | France | Hautes-Alpes - Les Combes | N44.8944°, E6.56460° | 7/21/2021 | CL |
| FR20-005-C-00 | France | Hautes-Alpes - Les Combes | N44.8944°, E6.56460° | 7/21/2021 | CL |
| FR21-001-C-00 | France | Hautes-Alpes - Col d'Arsine | N44.9749°, E6.41017° | 7/22/2021 | CL |
| FR21-002-C-00 | France | Hautes-Alpes - Col d'Arsine | N44.9749°, E6.41017° | 7/22/2021 | CL |
| FR21-003-C-00 | France | Hautes-Alpes - Col d'Arsine | N44.9740°, E6.41544° | 7/22/2021 | CL |
| FR21-004-C-00 | France | Hautes-Alpes - Col d'Arsine | N44.9740°, E6.41544° | 7/22/2021 | CL |

**Table S2:** Split times among population pairs (measured in thousands of years ago, and including all populations from Spain, and FR01 as a representative from the Alps), on the basis of the rCCR analysis with Relate.

| Pop | ES01 | ES02 | ES03 | ES04 | ES05 | ES06 | ES08 | ES09 | ES10 | ES17 | ES23 | ES24 | ES25 |
|---|---|---|---|---|---|---|---|---|---|---|---|---|---|
| **ES01** | 0 | 9.2 (7.4 - 12.4) | 19.7 (17.2 - 25.2) | 17.8 (15.5 - 23.3) | 32 (27.1 - 37.9) | 6.4 (5.4 - 7.5) | 130.6 (109.3 - 136.5) | 109 (87.3 - 115.7) | 27.5 (21.4 - 36.4) | 64.7 (47.4 - 85.2) | 37.9 (25.7 - 46.6) | 107.4 (89.9 - 114.3) | 131.7 (113.3 - 135.3) |
| **ES02** | 9.2 (7.4 - 12.4) | 0 | 13.1 (1.2 - 17) | 18.1 (13.2 - 28.3) | 29.3 (22.5 - 34.1) | 10.4 (7.7 - 13.9) | 132.9 (113.3 - 140.2) | 109 (86.5 - 119.8) | 25.9 (17.9 - 37.7) | 48.2 (32.9 - 72) | 43 (31.2 - 53.7) | 98.2 (73.3 - 111.3) | 134.1 (116.3 - 137.7) |
| **ES03** | 19.7 (17.2 - 25.2) | 13.1 (1.2 - 17) | 0 | 19.1 (15.6 - 24.1) | 21.9 (16.6 - 26.4) | 19.1 (15.5 - 24.5) | 137.7 (117.4 - 145.3) | 119.8 (100.1 - 126.1) | 22 (15.2 - 26.8) | 78 (53.7 - 90.7) | 41.1 (29.8 - 49.1) | 115.3 (100 - 122.7) | 142.7 (124.9 - 146.6) |
| **ES04** | 17.8 (15.5 - 23.3) | 18.1 (13.2 - 28.3) | 19.1 (15.6 - 24.1) | 0 | 9.6 (7.3 - 11.5) | 19.3 (14.9 - 23.7) | 136.5 (116.3 - 145.3) | 113.8 (92.6 - 122.9) | 25 (18.1 - 33.4) | 70.1 (43.4 - 82.2) | 15.7 (11.6 - 21.3) | 113.3 (91.5 - 124.9) | 140.2 (120.5 - 147.9) |
| **ES05** | 32 (27.1 - 37.9) | 29.3 (22.5 - 34.1) | 21.9 (16.6 - 26.4) | 9.6 (7.3 - 11.5) | 0 | 33.5 (27.1 - 38.3) | 135.3 (113.3 - 142.7) | 110.9 (89.5 - 119.8) | 22 (17.3 - 26.5) | 68.8 (43 - 80.1) | 20.2 (15.7 - 24.8) | 112.3 (93.1 - 122.7) | 141.4 (120.5 - 147.9) |
| **ES06** | 6.4 (5.4 - 7.5) | 10.4 (7.7 - 13.9) | 19.1 (15.5 - 24.5) | 19.3 (14.9 - 23.7) | 33.5 (27.1 - 38.3) | 0 | 130.6 (108.3 - 137.7) | 104.4 (82.9 - 112.8) | 34.3 (26.8 - 41.8) | 52.3 (41.1 - 67) | 40.7 (28.8 - 48.7) | 98.2 (74.6 - 109.3) | 135.3 (117.4 - 138.9) |
| **ES08** | 130.6 (109.3 - 136.5) | 132.9 (113.3 - 140.2) | 137.7 (117.4 - 145.3) | 136.5 (116.3 - 145.3) | 135.3 (113.3 - 142.7) | 130.6 (108.3 - 137.7) | 0 | 23.1 (18.8 - 28.2) | 144.6 (118.8 - 157.5) | 89.1 (73.3 - 95.7) | 127.1 (107.4 - 137.7) | 117.4 (100 - 126) | 140.2 (121.6 - 146.6) |
| **ES09** | 109 (87.3 - 115.7) | 109 (86.5 - 119.8) | 119.8 (100.1 - 126.1) | 113.8 (92.6 - 122.9) | 110.9 (89.5 - 119.8) | 104.4 (82.9 - 112.8) | 23.1 (18.8 - 28.2) | 0 | 126.1 (99.2 - 137.4) | 54 (40.7 - 63.6) | 102.7 (79.4 - 114.8) | 74.2 (60.4 - 90.3) | 121.8 (105.3 - 127.2) |
| **ES10** | 27.5 (21.4 - 36.4) | 25.9 (17.9 - 37.7) | 22 (15.2 - 26.8) | 25 (18.1 - 33.4) | 22 (17.3 - 26.5) | 34.3 (26.8 - 41.8) | 144.6 (118.8 - 157.5) | 126.1 (99.2 - 137.4) | 0 | 82.9 (53.1 - 100.9) | 42.1 (30.2 - 48.7) | 127.2 (99.2 - 140.9) | 149.6 (123.9 - 157.5) |
| **ES17** | 64.7 (47.4 - 85.2) | 48.2 (32.9 - 72) | 78 (53.7 - 90.7) | 70.1 (43.4 - 82.2) | 68.8 (43 - 80.1) | 52.3 (41.1 - 67) | 89.1 (73.3 - 95.7) | 54 (40.7 - 63.6) | 82.9 (53.1 - 100.9) | 0 | 67 (50.4 - 79.4) | 61.9 (48.2 - 78.7) | 120.5 (103.6 - 128.3) |
| **ES23** | 37.9 (25.7 - 46.6) | 43 (31.2 - 53.7) | 41.1 (29.8 - 49.1) | 15.7 (11.6 - 21.3) | 20.2 (15.7 - 24.8) | 40.7 (28.8 - 48.7) | 127.1 (107.4 - 137.7) | 102.7 (79.4 - 114.8) | 42.1 (30.2 - 48.7) | 67 (50.4 - 79.4) | 0 | 94 (62.4 - 108.3) | 132.9 (116.3 - 138.9) |
| **ES24** | 107.4 (89.9 - 114.3) | 98.2 (73.3 - 111.3) | 115.3 (100 - 122.7) | 113.3 (91.5 - 124.9) | 112.3 (93.1 - 122.7) | 98.2 (74.6 - 109.3) | 117.4 (100 - 126) | 74.2 (60.4 - 90.3) | 127.2 (99.2 - 140.9) | 61.9 (48.2 - 78.7) | 94 (62.4 - 108.3) | 0 | 50 (41.5 - 56.6) |
| **ES25** | 131.7 (113.3 - 135.3) | 134.1 (116.3 - 137.7) | 142.7 (124.9 - 146.6) | 140.2 (120.5 - 147.9) | 141.4 (120.5 - 147.9) | 135.3 (117.4 - 138.9) | 140.2 (121.6 - 146.6) | 121.8 (105.3 - 127.2) | 149.6 (123.9 - 157.5) | 120.5 (103.6 - 128.3) | 132.9 (116.3 - 138.9) | 50 (41.5 - 56.6) | 0 |
| **FR01** | 228.6 (205.5 - 243.3) | 226.6 (1.9 - 239) | 230.6 (211 - 245.4) | 228.6 (209.2 - 243.3) | 226.6 (209.2 - 241.1) | 228.6 (209.2 - 241.1) | 211 (193.1 - 224.6) | 208.9 (193.5 - 218.1) | 227.6 (208.9 - 248) | 214.8 (196.5 - 226.6) | 220.6 (203.6 - 236.9) | 198.3 (186.3 - 212.9) | 200.1 (188 - 214.8) |

**Table S3:** Drift rates scaled by $2 \times Ne$ calculated based on F3 statistics. Populations A and B are subject populations and population C is the reference population. Drift rates from the ancestor of population A and B to population A (drift rate A), to population B (drift rate B) and to population C (drift rate C; through the ancestor of population A, B and C).

| **Set** | **Population A** | **Population B** | **Population C** | **Drift rate A** | **Drift rate B** | **Drift rate C** |
|---|---|---|---|---|---|---|
| Set1 | W-CAN2 | W-CAN1 | FR01 | 0.177 | 0.166 | 3.353 |
| Set2 | W-CAN2 | ES17 | FR01 | 0.653 | 0.319 | 2.778 |
| Set3 | W-CAN1 | ES17 | FR01 | 0.638 | 0.301 | 0.2846 |
| Set4 | W-CAN2 | W-CAN1 | CE-CAN | 0.082 | 0.089 | 0.414 |
| Set5 | W-CAN2 | ES17 | CE-CAN | 0.223 | 0.264 | 0.408 |
| Set6 | W-CAN1 | ES17 | CE-CAN | 0.218 | 0.250 | 0.419 |

**Table S4:** Candidate genes that overlap the highest *3P-CLR* peaks, analyzed with different outgroups. Gene names, *A. thaliana* orthologs, branches with signatures of positive selection (P = private, A = ancestral), maximum normalized CLR score, molecular function from UniProt entries (Consortium 2023), and outgroup (F = French populations and E-CAN = Eastern Cantabrian populations)

| **Gene name** | ***A.a. gene ID*** | ***A.t.* ortholog** | **Branch(es)** | **Normalized CLR** | **Function** | **Outgroup** |
|---|---|---|---|---|---|---|
| *SC5D* | AA6G27580 | At3g02580 | P(W-CAN2) | 10.53 | Involved in the biosynthesis of precursor of growth-promoting brassinosteroids | F |
| *HPGT3* | AA6G26170 | At2g25300 | P(ES17) | 8.16 | Essential for AGP glycosylation | F |
| NA | AA6G26180 | At3g58210 | P(ES17) | 8.16 | May be linked to secondary cell wall biosynthesis | F |
| NA | AA7G10040 | NA | P(W-CAN1) | 7.49 | Unknown function | F |
| *CYP57* | AA7G10050 | At4g33060 | P(W-CAN1) | 6.81 | Functions as a molecular chaperone | F |
| *TWD40-2* | AA8G45050 | At5g24710 | P(W-CAN2) | 6.95 | Involved in clathrin mediated endocytosis | F |
| *KAS2* | AA2G23850 | At5g24710 | P(ES17) | 6.60 | Involved in the regulation of fatty acids ratios and cold response | F |
| *ORC3* | AA8G20070 | At5g16690 | A(W-CAN1, W-CAN2, ES17) | 3.19 | Involved in DNA replication | F |
| *BIR1* (paralog1) | AA8G20080 | At5g48380 | A(W-CAN1, W-CAN2, ES17) | 3.19 | Involved in negative regulation of plant immunity | F |
| *GA20OX4* | AA2G06360 | At1g60980 | A(W-CAN1, W-CAN2, ES17) | 3.05 | Key oxidase enzyme in the biosynthesis of gibberellin | F |
| *KIB4* | AA8G20040 | At3g03730 | A(W-CAN1, W-CAN2, ES17) | 2.97 | Mediates the ubiquitination and proteasomal degradation of target proteins | F |
| NA | AA8G20050 | NA | A(W-CAN1, W-CAN2, ES17) | 2.97 | Unknown function | F |
| *BIR1* (paralog2) | AA8G20060 | At5g48380 | A(W-CAN1, W-CAN2, ES17) | 2.97 | Involved in negative regulation of plant immunity | F |
| *LTP2* | AA2G11850 | At2g38530 | A(W-CAN1, W-CAN2, ES17) | 2.92 | May play a role in wax or cutin deposition in the cell walls | F |
| *AKCS9* | AA2G11860 | At2g38530 | A(W-CAN1, W-CAN2, ES17) | 2.92 | Potential lipid transfer protein | F |
| NA | AA2G11870 | NA | A(W-CAN1, W-CAN2, ES17) | 2.92 | Unknown function | F |
| *XBAT35* | AA3G27210 | At3g23280 | A(W-CAN1, W-CAN2, ES17) | 2.86 | Associated with defense and abiotic stress response | F |
| *PRAF1* | AA3G27220 | At1g76950 | A(W-CAN1, W-CAN2, ES17) | 2.86 | Small transmembrane protein involved in vesicle trafficking | F |
| NA | AA3G27230 | NA | A(W-CAN1, W-CAN2, ES17) | 2.86 | Unknown function | F |
| *SC5D* | AA6G27580 | At3g02580 | P(W-CAN2) | 17.43 | Involved in the biosynthesis of precursor of growth-promoting brassinosteroids | E-CAN |
| *BRR2C* | AA8G47440 | At5g61140 | P(W-CAN2) | 15.70 | Involved in spliceosome assembly, activation and disassembly | E-CAN |
| *PCR4* | AA3G22360 | At3g18460 | P(W-CAN1) | 14.63 | May be involved in heavy metals transport | E-CAN |
| *CASP* | AA3G22370 | At3g18480 | P(W-CAN1) | 14.63 | May be involved in intra-Golgi transport | E-CAN |
| *ILR2* | AA3G22380 | At5g08620 | P(W-CAN1) | 14.63 | Probably involved in the metabolism of auxin conjugates | E-CAN |
| *ASPG1* | AA3G22390 | At5g08620 | P(W-CAN1) | 14.63 | Aspartic protease involved in drought avoidance through abscisic acid signaling | E-CAN |
| NA | AA7G12830 | NA | P(W-CAN2) | 14.57 | Unknown function | E-CAN |
| *SDN2* | AA7G12840 | At5g05540 | P(W-CAN2) | 14.57 | Regulates small RNAs turnover | E-CAN |
| *DET1* (paralog1) | AA7G12770 | At4g10180 | P(W-CAN2) | 14.038 | Component of light signal transduction machinery | E-CAN |
| *DET1* (paralog2) | AA7G12780 | At4g10180 | P(W-CAN2) | 14.038 | Component of light signal transduction machinery | E-CAN |
| *APX2* | AA7G12790 | At3g09640 | P(W-CAN2) | 14.038 | Plays a key role in hydrogen peroxide removal | E-CAN |
| NA | AA7G12800 | NA | P(W-CAN2) | 14.038 | Unknown function | E-CAN |
| *NLE1* | AA7G12810 | At5g52820 | P(W-CAN2) | 14.038 | Involved in growth and developmental regulation | E-CAN |
| *LAX2* | AA6G21520 | At2g21050 | A(W-CAN1, W-CAN2) | 7.82 | Mediates auxin uptake | E-CAN |
| *CSP4* | AA6G21530 | At2g21060 | A(W-CAN1, W-CAN2) | 7.82 | Chaperone that binds to and unwinds RNA | E-CAN |
| *VTL5* | AA3G25050 | At3g25190 | A(W-CAN1, W-CAN2, ES17) | 7.68 | Vacuolar iron transporter | E-CAN |
| NA | AA8G30270 | NA | A(W-CAN1, W-CAN2, ES17) | 7.60 | Unknown function | E-CAN |
| NA | AA8G30280 | NA | A(W-CAN1, W-CAN2, ES17) | 7.60 | Unknown function | E-CAN |
| *MLP31* | AA1G29520 | At1g70840 | A(W-CAN1, W-CAN2, ES17) | 7.32 | Involved in defense and abiotic stress response | E-CAN |
| *BSMT1* (paralog1) | AA1G29530 | At3g11480 | A(W-CAN1, W-CAN2, ES17) | 7.32 | Catalyzes the methylation of salicylic acid | E-CAN |
| *BSMT1* (paralog2) | AA1G29540 | At3g11480 | A(W-CAN1, W-CAN2, ES17) | 7.32 | Catalyzes the methylation of salicylic acid | E-CAN |
| *MSH4* | AA4G18360 | At3g11480 | A(W-CAN1, W-CAN2, ES17) | 7.30 | Promotes homologous recombination | E-CAN |

**Table S5:** Top GO categories associated with genes that overlap the 0.5% high tail of *3P-CLR* using different groups of branches (terminal or ancestral) and outgroups (Eastern Cantabrian populations, or French populations).

| **GO enrichment analysis: terminal branches within Cantabria, with E-CAN as the outgroup** | | | | | | | | |
|---|---|---|---|---|---|---|---|---|
| **Rank** | **GO.ID** | **Term** | **Annotated** | **Significant** | **Ratio** | **Expected** | **P-value** | **AA gene IDs** |
| 1 | 1902476 | chloride transmembrane transport | 8 | 3 | 0.375 | 0.10 | 0.000120 | AA8G27720, AA1G19000, AA1G19050 |
| 2 | 0035652 | clathrin-coated vesicle cargo loading | 2 | 2 | 1.000 | 0.030 | 0.000170 | AA8G32590, AA3G19790, AA3G19790 |
| 3 | 0006582 | melanin metabolic process | 3 | 2 | 0.667 | 0.04 | 0.000500 | AA3G29310, AA3G28530 |
| 4 | 0045921 | positive regulation of exocytosis | 32 | 4 | 0.125 | 0.42 | 0.000750 | AA3G28530, AA7G06300 |
| 5 | 0001676 | long-chain fatty acid metabolic process | 2 | 2 | 1.000 | 0.03 | 0.000170 | AA3G28530, AA4G07770, AA4G07780, AA4G25990 |
| 6 | 0043162 | ubiquitin-dependent protein catabolic process | 15 | 3 | 0.200 | 0.19 | 0.000880 | AA3G29360, AA3G22810, AA3G18860 |
| 7 | 0050848 | regulation of calcium-mediated signaling | 4 | 2 | 0.500 | 0.050 | 0.000990 | AA3G28530, AA7G12960 |
| 8 | 0000381 | regulation of alternative mRNA splicing, via spliceosome | 19 | 3 | 0.158 | 0.25 | 0.00179 | AA4G18110, AA4G18130, AA2G07210 |
| 9 | 0051209 | release of sequestered calcium ion into cytosol | 7 | 2 | 0.286 | 0.090 | 0.003370 | AA3G28530, AA7G12960 |
| 10 | 1901019 | regulation of calcium ion transmembrane transport | 7 | 2 | 0.286 | 0.09 | 0.00337 | AA3G28530, AA3G34510 |
| 11 | 0051345 | positive regulation of hydrolase activity | 2 | 2 | 0.060 | 1.090 | 0.004750 | AA3G29310, AA3G22810, AA8G15710, AA8G32590, AA7G06300 |
| **GO enrichment analysis: ancestral branch, with E-CAN as the outgroup** | | | | | | | | |
| **Rank** | **GO.ID** | **Term** | **Annotated** | **Significant** | **Ratio** | **Expected** | **P-value** | **AA gene IDs** |
| 1 | 0071281 | cellular response to iron ion | 64 | 5 | 0.078 | 0.31 | 0.000015 | AA3G24350, AA3G25020, AA3G25030, AA3G25040, AA3G25050 |
| 2 | 0071452 | cellular response to singlet oxygen | 5 | 2 | 0.400 | 0.02 | 0.000230 | AA6G21120, AA8G19800 |
| 3 | 0010358 | leaf shaping | 12 | 2 | 0.167 | 0.06 | 0.001500 | AA1G29450, AA5G19050 |
| 4 | 0031670 | cellular response to nutrient | 12 | 2 | 0.167 | 0.06 | 0.001500 | AA6G21120, AA8G19800 |
| 5 | 0010075 | regulation of meristem growth | 59 | 3 | 0.051 | 0.29 | 0.002980 | AA1G29450, AA5G19050, AA6G21120 |
| 6 | 0046283 | anthocyanin-containing compound metabolic process | 62 | 3 | 0.048 | 0.30 | 0.003430 | AA3G29350, AA6G21120, AA8G30040 |
| **GO enrichment analysis: terminal branches within Cantabria, with FR as the outgroup** | | | | | | | | |
| **Rank** | **GO.ID** | **Term** | **Annotated** | **Significant** | **Ratio** | **Expected** | **P-value** | **AA gene IDs** |
| 1 | 0042775 | mitochondrial ATP synthesis coupled electron transport | 36 | 5 | 0.1389 | 0.60 | 0.000320 | AA8G18520, AA3G20490, AA6G12340. AA7G10960, AA5G06640 |
| 2 | 0006857 | oligopeptide transport | 78 | 6 | 0.077 | 1.31 | 0.00201 | AA3G19710, AA4G25380, AA1G12910, AA8G14650, AA8G17860, AA8G31710 |
| 3 | 0009150 | purine ribonucleotide metabolic process | 462 | 17 | 0.037 | 7.76 | 0.002260 | AA8G18520, AA1G15040, AA1G23770, AA1G35170, AA3G06510, AA3G20490<br>AA4G33220, AA6G12340, AA7G10960, AA3G19400, AA3G28870, AA3G20560<br>AA3G29930, AA5G06640, AA7G10920, AA7G11030, AA7G11250 |
| 4 | 0042538 | hyperosmotic salinity response | 84 | 6 | 0.071 | 1.41 | 0.002920 | AA4G25380, AA7G11040, AA7G26010, AA7G26020, AA8G17870, AA2G15990 |
| **GO enrichment analysis: ancestral branch, with FR as the outgroup** | | | | | | | | |
| **Rank** | **GO.ID** | **Term** | **Annotated** | **Significant** | **Ratio** | **Expected** | **P-value** | ***A. alpina* gene IDs** |
| 1 | 0046580 | negative regulation of Ras protein signal transduction | 15 | 3 | 0.200 | 0.11 | 0.000150 | AA1G15580, AA4G16250, AA4G16260 |
| 2 | 0051123 | RNA polymerase II transcriptional preinitiation complex assembly | 33 | 3 | 0.091 | 0.23 | 0.0016 | AA1G22980, AA3G23040, AA3G23050 |
| 3 | 0045944 | Positive regulation of transcription by RNA polymerase II | 156 | 5 | 0.032 | 1.10 | 0.0016 | AA3G23040, AA3G23050, AA4G16250, AA4G16260, AA5G21780 |

**Table S6:** Candidate genes associated with response to water deprivation and heat. Gene names, *A. thaliana* orthologs, branches on which signatures are detected (T = terminal, A = ancestral), maximum normalized CLR score, GO term, and GO ID.

| Gene name | *A.a. gene ID* | *A.t.* ortholog | Branch(es) | Normalized CLR (max) | GO term | GO ID |
|---|---|---|---|---|---|---|
| *ASPG1* | AA3G22390 | At3g18490 | T(W-CAN1) | 14.63 | response to water deprivation | 0009414 |
| *NAC055* | AA3G18870 | At3g15500 | T(W-CAN1), T(ES17) | 11.02 | response to water deprivation | 0009414 |
| *PAL2* | AA5G19770 | At3g53260 | T(W-CAN2) | 8.90 | response to water deprivation | 0009414 |
| *RH26* | AA8G10500 | At5g08610 | T(W-CAN1) | 7.09 | response to water deprivation | 0009414 |
| *RH25* | AA8G10510 | At5g08620 | T(W-CAN1) | 7.09 | response to water deprivation | 0009414 |
| *PARG2* | AA6G27810 | At2g31865 | T(ES17), T(W-CAN1) | 6.94 | response to water deprivation | 0009414 |
| *ZHD11* | AA2G15790 | At1g69600 | T(W-CAN2) | 6.90 | response to water deprivation | 0009414 |
| *DRIPH* | AA3G27540 | At3g23060 | T(W-CAN2) | 6.39 | response to water deprivation | 0009414 |
| *ARR1* | AA3G20460 | At3g16857 | T(W-CAN2) | 6.22 | response to water deprivation | 0009414 |
| *RBG4* | AA3G26460 | At3g23830 | T(W-CAN2) | 6.03 | response to water deprivation | 0009414 |
| *HSP15.7* | AA6G27830 | At5g37670 | T(ES17), T(W-CAN1) | 6.55 | response to heat | 0009408 |
| *MYB94* | AA5G09300 | At3g47600 | A(W-CAN2, ES17) | 5.52 | response to water deprivation | 0009414 |
| *ASPG1* | AA3G22390 | At3g18490 | A(W-CAN1, W-CAN2, ES17) | 5.18 | response to water deprivation | 0009414 |
| *ERF011* | AA5G15450 | At3g50260 | A(W-CAN1, W-CAN2, ES17) | 5.18 | response to water deprivation | 0009414 |
| *Cpn10* | AA1G29480 | At1g14980 | A(W-CAN1, W-CAN2, ES17) | 6.55 | response to heat | 0009408 |
| *FKBP62* | AA3G24960 | At3g25230 | A(W-CAN1, W-CAN2, ES17) | 4.58 | response to heat | 0009408 |

**Table S7:** Candidate genes related to vernalization. Gene names, *A. thaliana* orthologs, branches on which signatures are detected (T = terminal, A = ancestral), maximum normalized CLR score, GO term, and GO ID.

| Gene name | *A.t.* ortholog | Branch(es) | Normalized CLR | GO term | GO ID |
|---|---|---|---|---|---|
| *VIP4* | At5g61150 | T(W-CAN2) | 6.27 | vernalization response | 0042538 |
| *VIL1* | At3g24440 | T(ES17) | 5.00 | vernalization response | 0042538 |
| *FIP1* | At2g06005 | T(W-CAN2) | 11.17 | protein binding | 0005515 |

**Table S8:** Top GO categories associated with genes that differ significantly in the rate ratio of non-synonymous to synonymous polymorphisms between populations from Cantabria and from the French Alps.

| **GO ID** | **GO Term** | **Annotated** | **Significant** | **Ratio** | **Expected** | **P-value** | ***A.t.* orthologs** |
|---|---|---|---|---|---|---|---|
| GO:0006468 | protein phosphorylation | 1224 | 41 | 0.033 | 23.80 | 0.00051 | AA1G09570,AA1G14670,AA1G14770, AA1G29200,AA1G40690,AA2G06560, AA2G11400,AA2G14830,AA2G15330, AA2G17730,AA2G25770,AA3G01250, AA3G12680,AA3G24460,AA3G34370, AA4G03060,AA4G16580,AA4G40200, AA4G40560,AA4G41130,AA4G41720, AA5G07600,AA5G13790,AA5G22880, AA6G21990,AA6G32380,AA7G08710, AA7G25120,AA8G15950,AA8G19910, AA8G25650,AA8G43370,AA8G45790, AA8G50650,AA1G33160,AA7G03270, AA8G16830,AA3G26300,AA3G30970, AA7G23150,AA8G02220 |
| GO:0010583 | response to cyclopentenone | 109 | 8 | 0.073 | 2.12 | 0.00131 | A1G10960,AA1G38120,AA2G25770, AA3G14410,AA5G31250,AA8G46010, AA3G02390,AA6G02740 |
| GO:0120010 | intermembrane phospholipid transfer | 13 | 3 | 0.23 | 0.25 | 0.00181 | AA2G24340,AA6G22040, AA7G02050 |
| GO:0007178 | cell surface receptor protein serine/threonine kinase signaling pathway | 183 | 10 | 0.055 | 3.56 | 0.00319 | AA1G09570,AA1G14770,AA2G06560, AA2G11400,AA3G01250,AA3G34370, AA4G16580,AA4G40560,AA5G22880, AA8G50650 |
| GO:0045806 | negative regulation of endocytosis | 5 | 2 | 0.4 | 0.10 | 0.00363 | AA1G04680,AA6G10170 |

**Table S9:** Allele frequencies of semi- and non-conservative amino acid changes and deletions in *FTIP7* paralogs for Cantabrian Mountains groups and French populations. Amino acid changes are indicated using one-letter amino acid codes and arrows and indels are indicated using one-letter amino acid codes followed by three-letter codes for deletions or duplications (del/dup). Stars denote non-conservative and circles denote semi-conservative amino acid changes or indels based on chemical properties and position.

| **Variant** | | | | | | | **Allele frequency [%]** | | | | |
|---|---|---|---|---|---|---|---|---|---|---|---|
| ***A. alpina* ID** | **Chr** | **Position** | **Ref** | **Alt** | **Substitution/Indel** | **Domain** | **W-CAN1** | **W-CAN2** | **ES17** | **CE-CAN** | **FRANCE** |
| AA1G04680 | chr1 | 1769334 | C | T | G → R (683)* | C2 | 97.94 | 100.00 | 33.00 | 44.00 | 0.00 |
| AA1G04680 | chr1 | 1769695 | ACTGTTATTGTTG | A | NNNS del (559-562)* | - | 3.09 | 0.00 | 66.67 | 4.00 | 0.00 |
| AA1G04680 | chr1 | 1770704 | A | C | M → R (226)* | - | 0.00 | 0.00 | 66.67 | 4.00 | 0.00 |
| AA1G04680 | chr1 | 1770838 | TTGC | T | Q del (179)° | - | 8.76 | 0.00 | 0.00 | 0.00 | 0.00 |
| AA1G04680 | chr1 | 1770838 | TTGC | TTGCTGC | Q dup (179)° | - | 3.09 | 0.00 | 66.67 | 4.00 | 43.02 |
| AA1G04680 | chr1 | 1770853 | CTGTTGTTGT | CTGTTGT | Q del (174)° | - | 0.00 | 22.30 | 0.00 | 0.00 | 0.00 |
| AA1G04680 | chr1 | 1770853 | CTGTTGTTGT | CTGTTGTTGTTGT | Q dup (173)° | | 0.00 | 0.00 | 0.00 | 0.04 | 0.00 |
| AA1G04680 | chr1 | 1770887 | A | G | L → S (165)* | - | 0.00 | 22.30 | 0.00 | 0.00 | 0.00 |
| AA6G10170 | chr6 | 5698025 | A | G | N → A (16)° | - | 2.06 | 0.00 | 66.67 | 100.00 | 83.26 |
| AA6G10170 | chr6 | 5698052 | G | A | T → A (26)° | - | 97.94 | 100.00 | 33.33 | 100.00 | 83.26 |
| AA6G10170 | chr6 | 5701344 | G | A | L → F (867)° | PRT-C | 7.73 | 17.57 | 66.67 | 0.00 | 0.00 |
| AA6G10170 | chr6 | 5701694 | C | T | T → I (984)* | PRT-C | 9.97 | 17.57 | 66.67 | 0.00 | 39.30 |
| AA6G10170 | chr6 | 5701769 | C | T | T → M (1009)* | PRT-C | 16.49 | 30.41 | 0.00 | 0.00 | 0.00 |

**Table S10:** Population frequencies of six non-synonymous mutations at *FRL1* in the Cantabrian Mountains and in France, and the flowering behavior of the corresponding populations. The proportion of missing genotype data is shown in brackets. Flowering behavior without vernalization for the corresponding populations (mentioned in brackets) from (Wunder et al. 2023) is categorized in early, late, and no onset of flowering. For the Cantabrian Mountains, we reported the range across populations (minimum-maximum).

| | **Allele frequency [%]** | | | | | | **flowering behavior [%]** | | |
|---|---|---|---|---|---|---|---|---|---|
| **Population(s)** | **12618926** | **12618944** | **12619056** | **12619062** | **12619091** | **12619412** | **early onset** | **late onset** | **no onset** |
| Cantabria | 100.0 | 100.0 | 100.0 | 100.0 | 100.0 | 100.0 | 0-2 | 0-1 | 97-100 |
| FR02(F2) | 100.0 | 100.0 | 100.0 | 100.0 | 100.0 | 100.0 | 1 | 23 | 76 |
| FR07(F7) | 100.0 | 100.0 | 100.0 | 100.0 (9.0) | 100.0 | 100.0 | 21 | 21 | 58 |
| FR10(F6) | 100.0 | 100.0 | 100.0 | 100.0 | 100.0 | 100.0 | 0 | 0 | 100 |
| FR20 | 83.3 | 83.3 | 83.3 | 83.3 | 83.3 (33.3) | 100.0 (67) | 4 | 46 | 50 |
| FR15 | 37.5 | 25.0 | 28.6 (12.5) | 28.6 (12.5) | 28.6 (12.5) | 37.5 | NA | NA | NA |
| FR08(F4) | 0.0 | 0.0 | 0.0 | 1.4 | 1.4 | 1.4 | 58 | 3 | 39 |
| FR01(F1a) | 0.0 | 0.0 | 0.0 | 0.0 | 0.0 | 0.0 | 84 | 11 | 5 |
| FR03(F1b) | 0.0 | 0.0 | 0.0 | 0.0 | 0.0 | 0.0 | 14 | 44 | 42 |

### Supplementary Figures

##### Figure S1

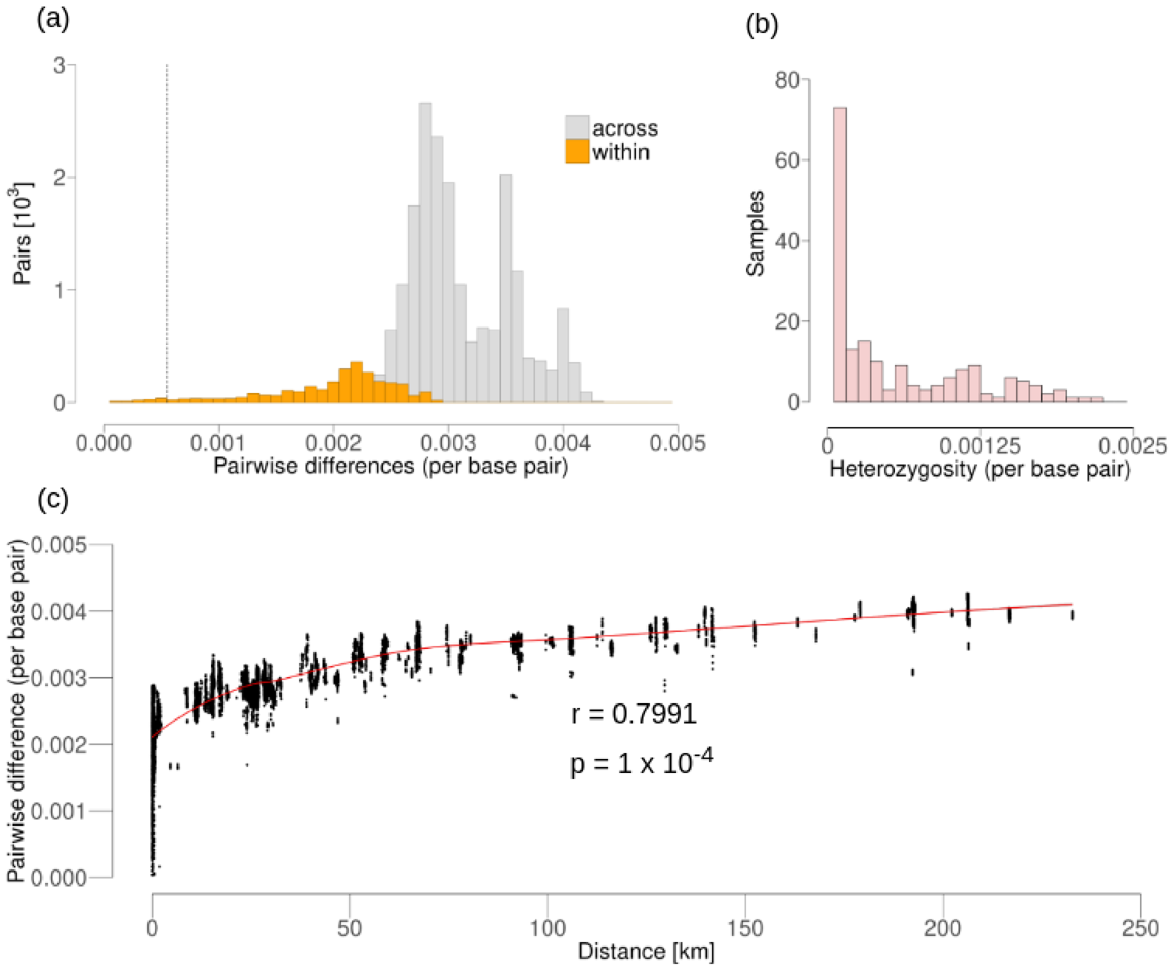


Distribution of genetic differences among pairs of individuals from the Cantabrian Mountains. **a:** Pairwise differences per base pair across (grey) and within (yellow) populations. The dashed line indicates the cutoff used to select unrelated samples. **b:** Distribution of heterozygote sites per base pair and per sample. **c:** Pairwise differences as a function of geographical distances. The figure shows the Mantel test correlation statistic (r) and p-value (p), and a local polynomial regression in red.

**Figure S2**

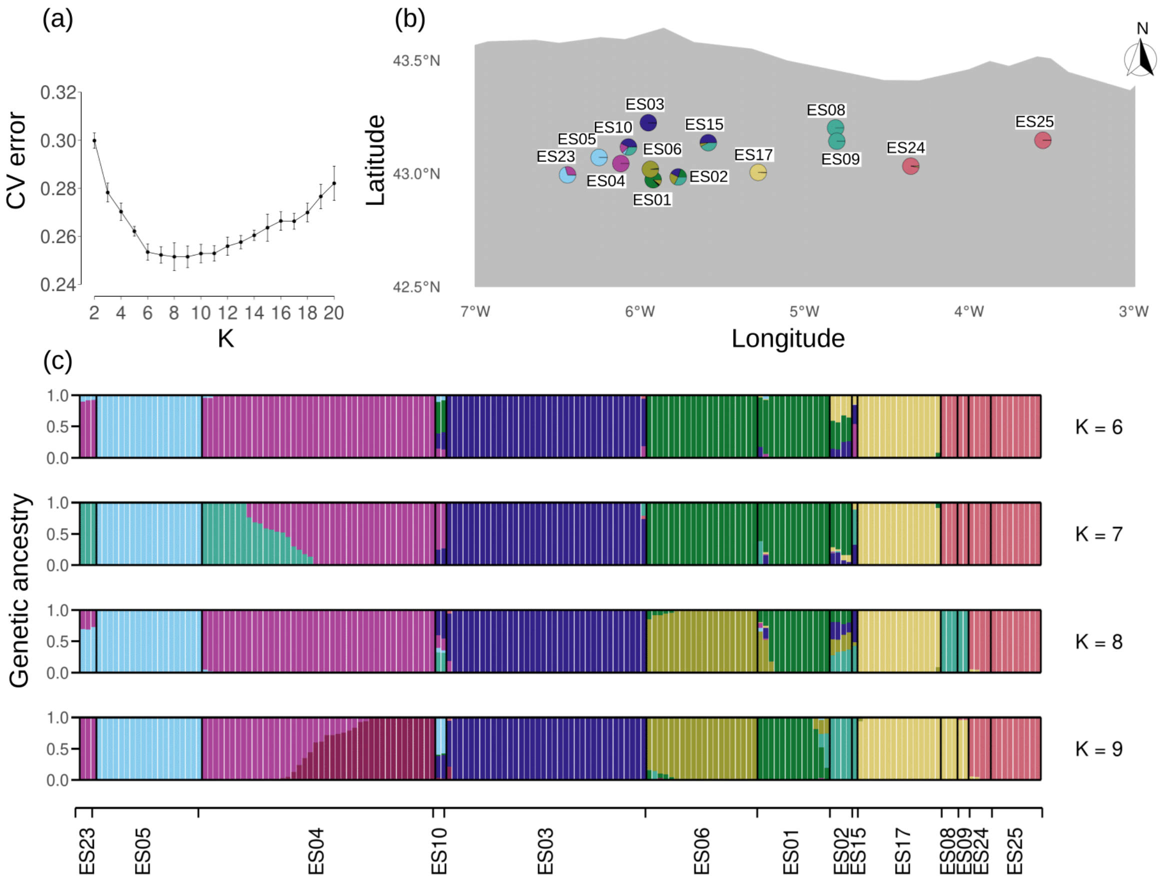


Genetic ancestry groups identified in Cantabrian Mountains *A. alpina*, on the basis of ADMIXTURE. **a:** Mean cross-validation (CV) errors with two to 20 ancestry groups (K). Bars show standard deviation across 10 replicates. **b:** Geographical distribution of ancestry groups inferred from the run with the lowest average CV error ($K = 8$) across replicates. Pie charts represent populations and colors represent ancestry groups. **c:**. Genetic ancestry proportions per individual for runs using six to nine ancestry groups.

**Figure S3**

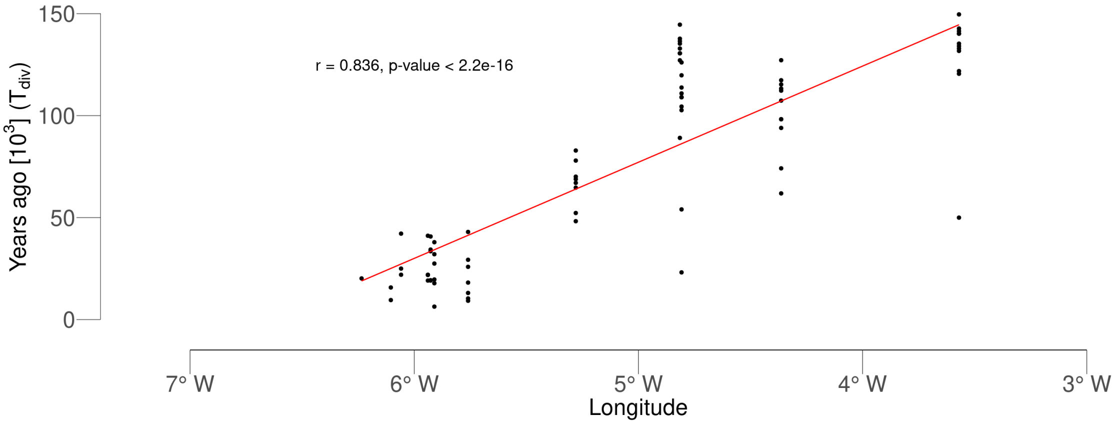


Pairwise split times between Cantabrian Mountains populations as a function of the longitude of the eastern-most population in each pair. The red line represents a linear fit, and the Pearson correlation coefficient (r) and p-value are shown within the figure.

**Figure S4**

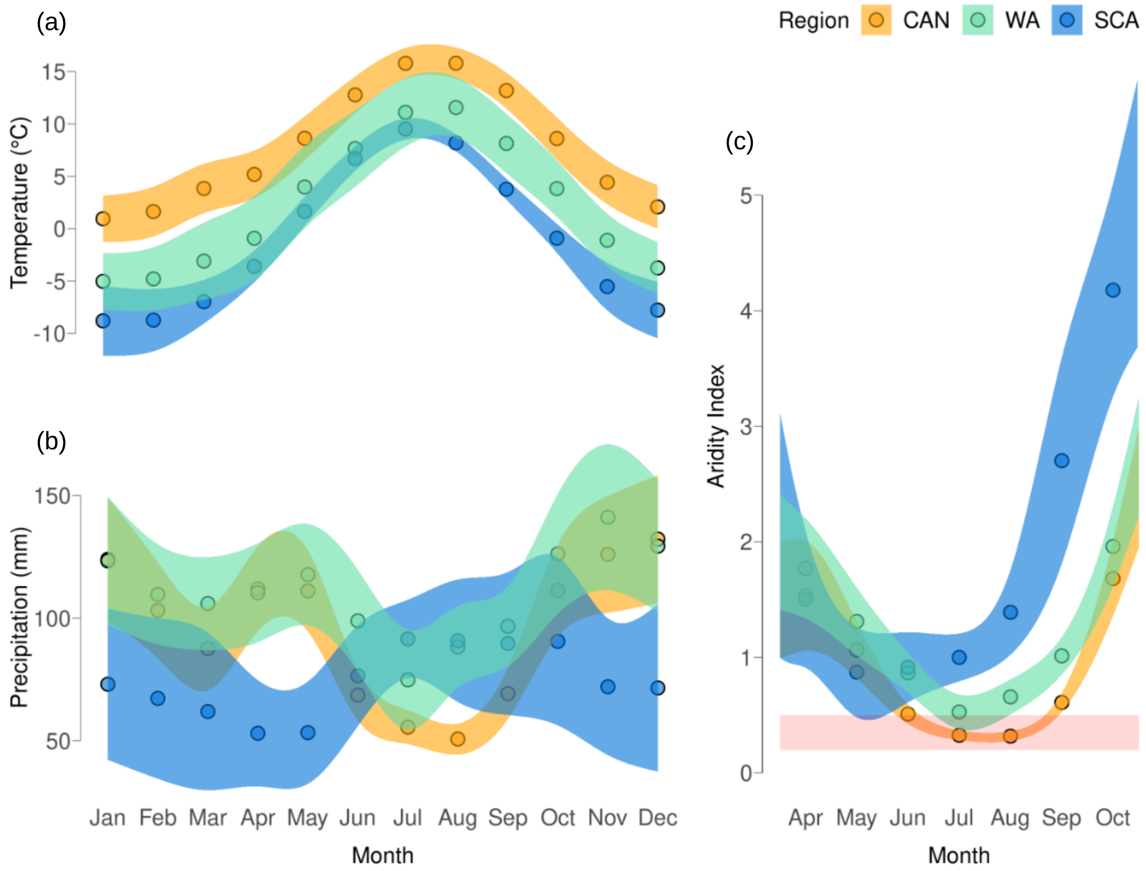


Yearly course of **a:** temperature, **b:** precipitation and **c:** aridity index (AI) across European populations from the Cantabrian Mountains (CAN), Western Alps (WA), and Scandinavia (SCA). Monthly averages across populations are indicated by points, and standard deviations are shown by smoothed ribbons. The red shaded area in (c) indicates semi-arid conditions defined as $0.2 \leq AI \leq 0.5$. Note that lower values for AI correspond to more arid climates that higher values.

**Figure S5**

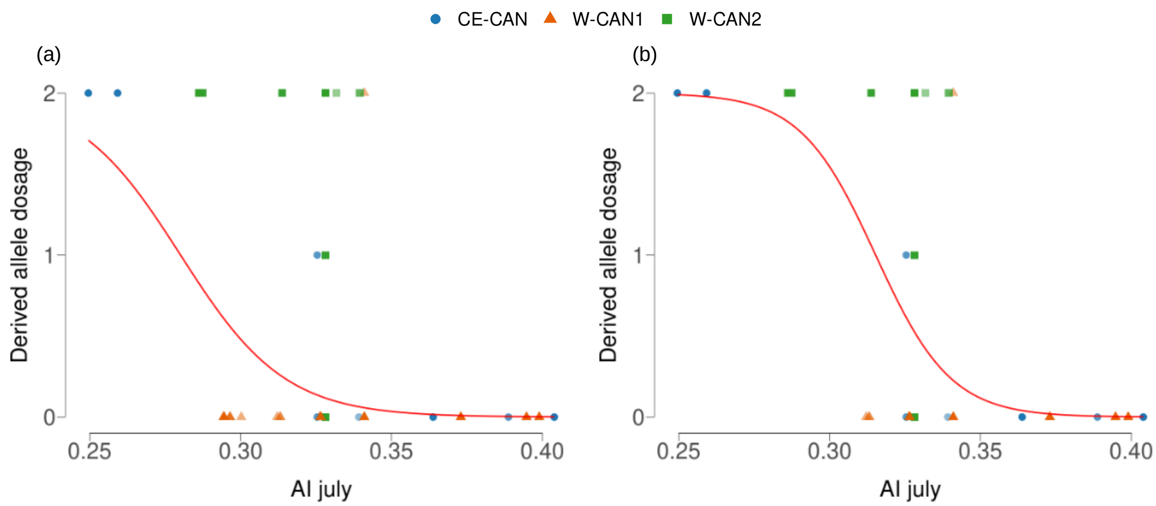


Binomial regression of the derived allele dosage at the 3bp insertion at *NAC055* and aridity index (AI) in July, with four principal coordinates from genome-wide structure analysis included as covariates. Data points represent single accessions, and they are shaped and colored by structure group. The red line shows the predicted relationship between allele dosage and AI. **a:** Binomial regression including all 211 accessions from the Cantabrian Mountains. **b:** Binomial regression excluding populations ES02, ES10, and ES15 (n = 200), which were collected at sites with micro climates very distinct from the surrounding climate (details in Supplementary Text).

**Figure S6**

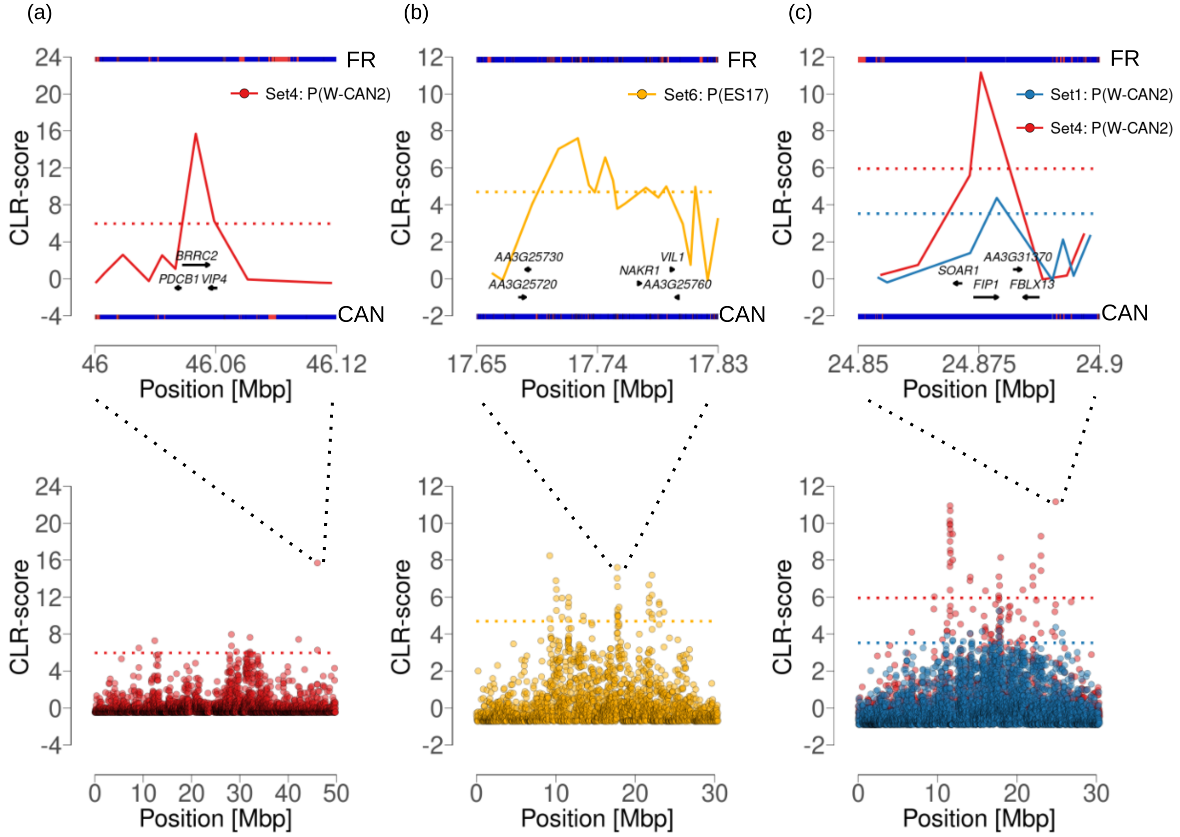


Standardized composite likelihood ratio scores for flowering genes loci *VIP4* (**a**), *VIL1* (**b**) and *FIP1* (**c**), shown zoomed in (top) and across the chromosome (bottom). Genes are indicated by arrows and labeled with their *A. thaliana* orthologs or, if no ortholog exists, by their *A. alpina* gene identifiers. Dotted lines show 0.5% most extreme CLR scores for each analysis. Segments in the top panels show multicopy (red) and single-copy (blue) regions in France (top segment) and in the Cantabrian Mountains (bottom segment) inferred by ParaMask. **a:** Scores for the private branch of W-CAN2 at the *VIP4* locus. **b:** Scores for the private branch of ES17 at the *VIL1* locus. **c:** Scores for the private branch of W-CAN2 and different *3P-CLR* analysis (blue - Set1; red - Set4) at the *FIP1* locus.

**Figure S7**

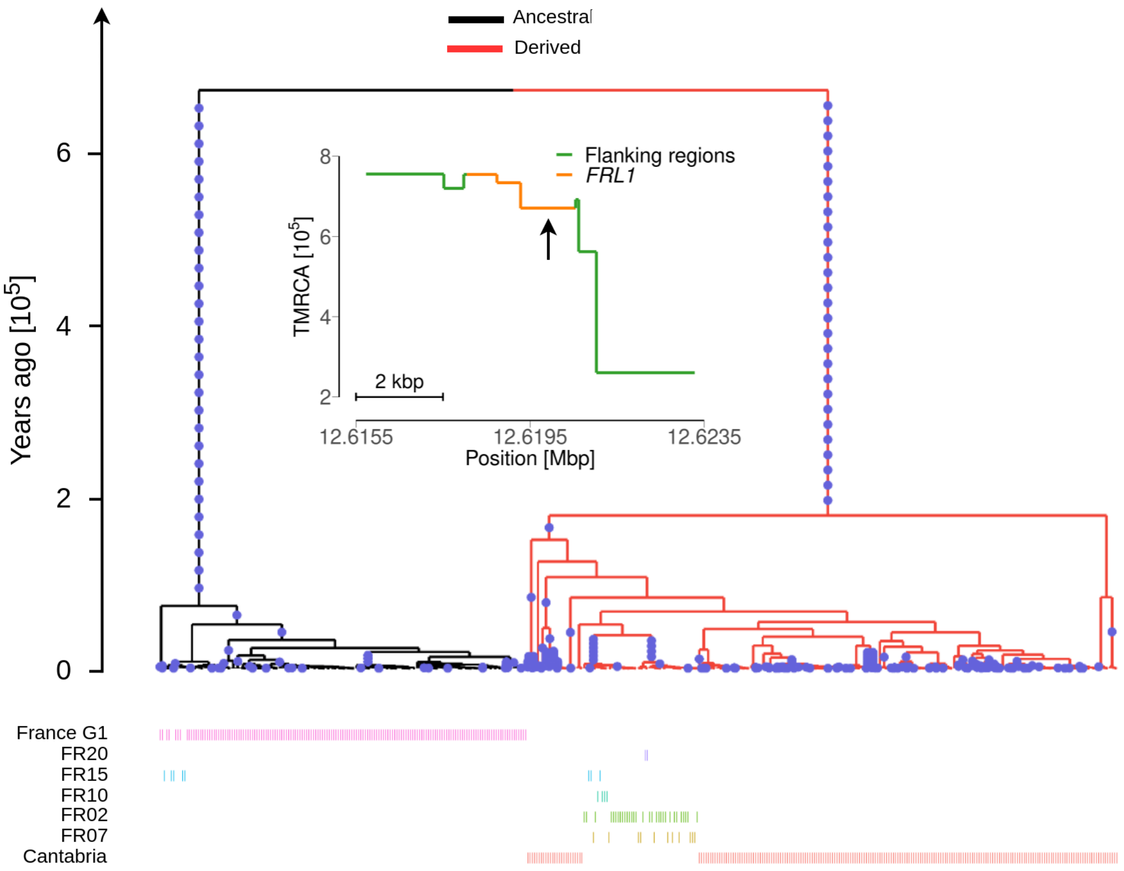


The genealogy at the *FRL1* locus that spans one of the six non-synonymous mutations. Red branches indicate the lineages that carry the derived alleles at this mutation, and black branches represent lineages that carry the ancestral allele. France G1 (group 1) includes populations FR01, FR03, FR04, FR06, FR08, FR16, FR17, FR18, FR19, and FR21. The inset shows the time to most recent common ancestor (tmrca) distribution across genealogies at the *FRL1* locus. The tmrca of the genealogy depicted here is marked by an arrow.

#### Figure S8

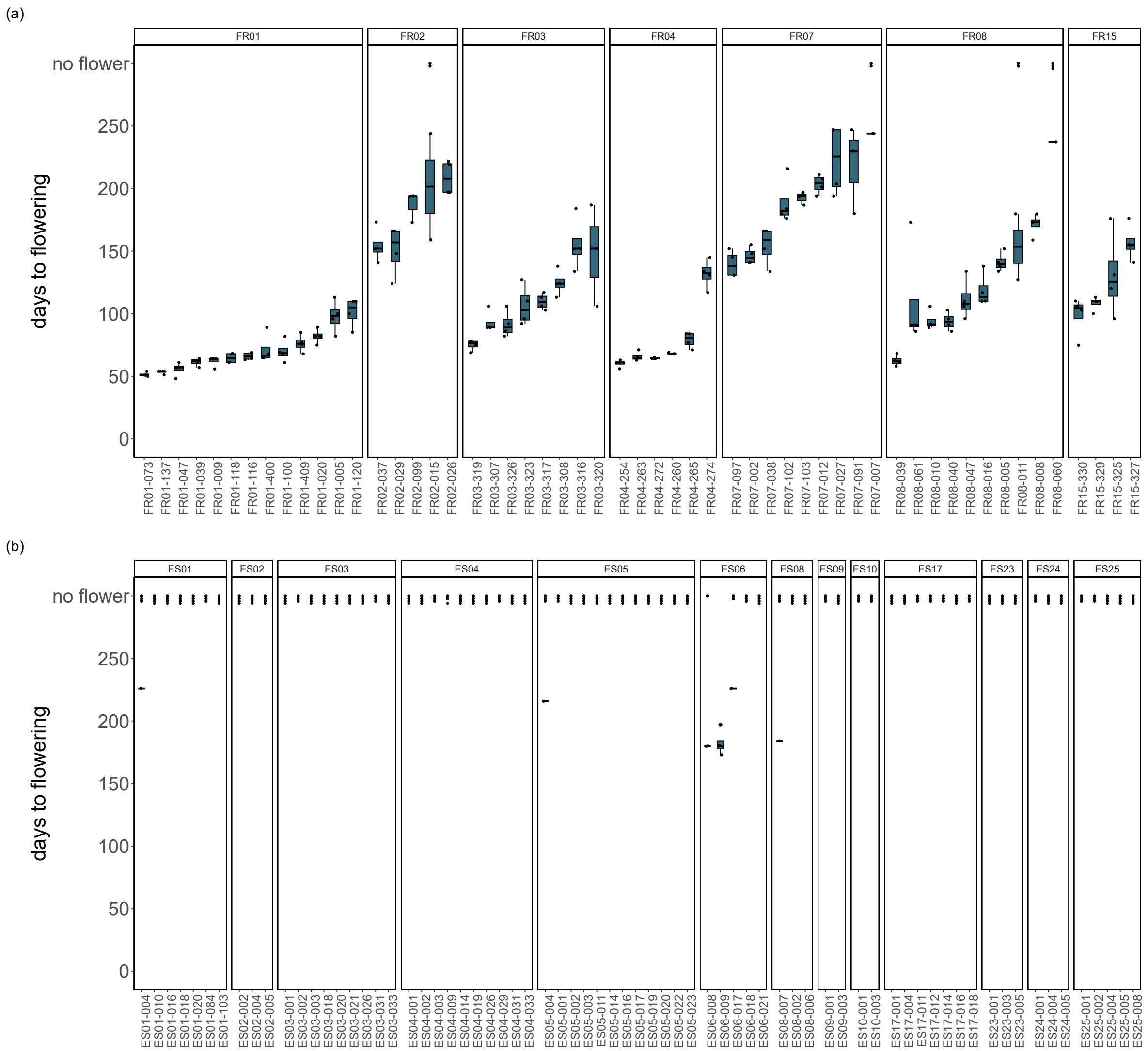


Time of flowering onset (days to flowering) for accessions from (**a**) French populations and (**b**) Cantabrian Mountains populations in greenhouse conditions without vernalization. Flowering time is measured as the number of days between sowing and the opening of the first flower.
